# MiniRead: a simple and inexpensive do-it-yourself device for multiple analyses of micro-organism growth kinetics

**DOI:** 10.1101/2023.12.21.572596

**Authors:** Matthieu Falque, Aurélie Bourgais, Fabrice Dumas, Mickaël de Carvalho, Célian Diblasi

**Author notes:** **TAKE AWAY:** We present a simple and inexpensive do-it-yourself turbidimeter for cell suspensions. It records automatically micro-organisms growth kinetics in 96-well plate without computer. Temperature is controlled in plate bottom and lid independently to avoid mist on the lids. An associated R package allows automatic kinetics analysis to estimate growth parameters.

## Abstract

Fitness in micro-organisms can be proxied by growth parameters on different media and/or temperatures. This is achieved done by measuring optical density at 600 nm using a spectrophotometer, which measures the effect of absorbance and side scattering due to turbidity of cells suspensions. However, when growth kinetics must be monitored in many 96-well plates at the same time, buying several 96-channels spectrophotometers is often beyond budgets. The MiniRead device presented here is a simple and inexpensive do-it-yourself 96-well temperature-controlled turbidimeter designed to measure the interception of white light *via* absorption or side scattering through liquid culture medium. Turbidity is automatically recorded in each well at regular time intervals for up to several days or weeks. Output tabulated text files are recorded into a micro-SD memory card to be easily transferred to a computer. We propose also an R package which allows (1) to compute the non-linear calibration curves required to convert raw readings into cell concentration values, and (2) to analyze growth kinetics output files to automatically estimate growth parameters such as lag time, maximum growth rate, or cell concentration at the plateau.

The MiniRead project is freely available under GPL license from https://forgemia.inra.fr/gqe-base/MiniRead. The project includes (1): user manual (Supplementary Material 1), firmware, list of electronic components and printed circuit board manufacturing files for the MiniRead device, and (2) a release containing the MiniRead R package for calibration and data analysis (Supplementary Material 2) with its tutorial (Supplementary Material 3). Detailed device building instructions are available at https://moulon.inrae.fr/materiel_labo/miniread/.

## INTRODUCTION

Although the ramifications of the concept of fitness can be quite bushy (Krimbas 2004), the individual absolute fitness of an organism may be defined in a simplified way as the number of descendants that this organism will produce in a given environment. In asexually-reproducing unicellular micro-organisms, different components of fitness can be experimentally measured quite directly by characterizing quantitatively the kinetics of mitotic cell divisions in different controlled environments. This may require to repeat growth experiments on many different chemical (composition of the culture media) and/or physical (e.g. growth temperature) environmental conditions. Therefore, fitness measurements often need to carry out many growth assays in parallel, even if the number of different sample strains or populations is initially limited. In that context, 96-well plates are appropriate to carry out growth experiments, and many manufacturers propose optical plate readers adapted to that format, which allow automatic growth kinetics monitoring.

However, most readers are designed to measure absorbance at precise wavelengths or even fluorescence, whereas monitoring growth kinetics does not require such sophisticated measurements. Growth experiments with micro-organisms are typically carried out on either solid gelled (e.g. see Barton et al. 2018) or liquid culture media, using 96 or even 384-wells plates (Jung et al. 2015). On solid medium, growth can be monitored *via* measuring either the size of colonies by image analysis, or the level of light interception through a small column of medium on top of which the colony is growing. In liquid medium, cell concentration can be simply proxied by measuring the turbidity of the broth for different kinds of micro-organisms (Hills 2017; Luong et al. 2011; Nwoba et al. 2022), related to the amount of light which is side-scattered or absorbed when a light beam is passing through a column of cell suspension. When using flat-bottom wells, the same holds true even if the cells have sedimented and form a layer at the bottom of the well. This makes agitation not necessary provided there is enough gas exchanges for cells to grow under the desired conditions. Long ago already, that turbidity approach had given rise to a simple device to monitor growth (Watson et al. 1969). However, by that time, the state-of-the-art of electronics offered limited possibilities to efficient home-made implementations, and since then, we are not aware of any other published simple device of the same type. Our objective here was to build on the same idea to design and assemble a small and inexpensive 96-well home-made turbidimeter, of which one could produce several units with a modest budget.

The main requirements for the MiniRead device were: (1) use 150 – 200 μL volume of liquid culture in 96-wells plates without agitation, (2) record light interception by the column of cell suspension in each well at regular time intervals, (3) accurately control the temperature of the bottom of the culture plate (up to 45°C), (4) control independently the temperature of the plate lid, which should be kept e.g. two degrees warmer than the bottom to avoid mist condensation – that point is critical for optical readings while keeping sterile conditions in the wells, (5) store data autonomously without the need for a computer, (6) simple assembly within the reach of non-professional electronicians having good DIY soldering experience, and (7) low cost allowing to build five such readers within a total budget of 1000$.

## MATERIALS AND METHODS

### 1. Hardware description and working principle

The detailed user manual of the MniRead turbidimeter is available in Supplementary Material 1. The device (Figure 1) is build around a widely-used Arduino MEGA256 board (see specifications at https://store.arduino.cc/collections/boards/products/arduino-mega-2560-rev3) which includes a C-programmable ATmega2560 micro-controller with a large number of digital and analog input/output pins. The MiniRead is power supplied by an external regulated power supply (desktop power adapter) 12V DC 5A 60W from RS components (https://fr.rs-online.com). This 12V input directly powers the two heating elements *via* Darlington transistors, while there is an additional 5V regulator in the MniRead to supply the Arduino and its external electronics.

**Figure 1.**
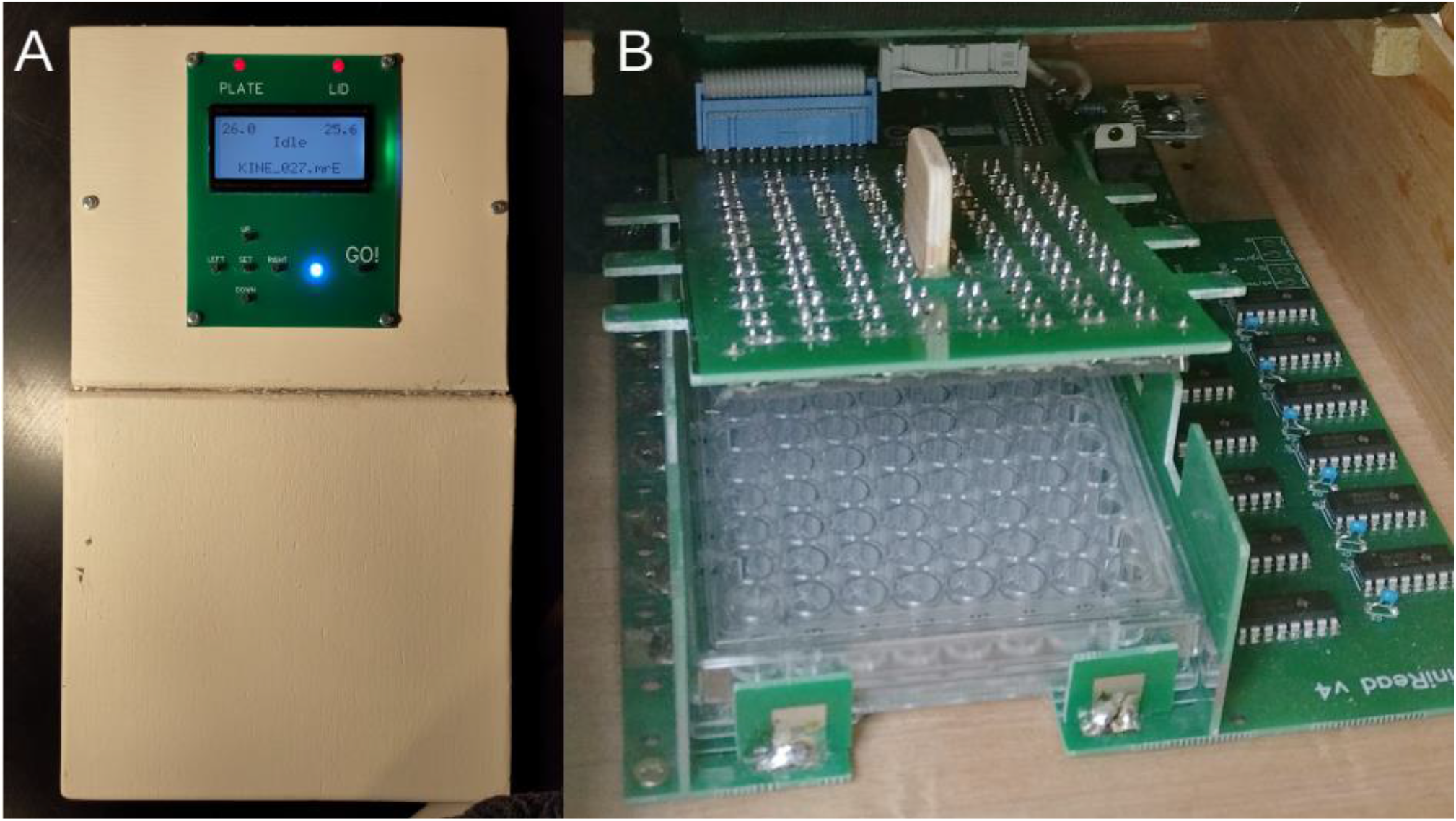
General view of the MiniRead. A: top view with the user interface and the front cover. B: inside view showing the plate holder and the heated lid with the 96 LEDs.

For the light sensors, we used 96 cheap GL5516 light-dependent resistors (LDRs) from Aliexpress (https://fr.aliexpress.com/) but we quality-controlled each LDR by measuring its resistance under a fixed level of illumination. This procedure is recommended, since it allowed us to identify and discard some defective components with outlier resistance values. More details are available on our web site at https://moulon.inrae.fr/materiel_labo/miniread/. We used also cheap 3mm white light emitting diodes (LEDs) from the same provider as sources of illumination. We did only a qualitative quality control of each LED by checking its illumination under a constant 20mA current, and found very few defective LEDs.

Analog signals from the 96 LDRs are multiplexed using twelve CD4051BE analog 8-to-1 multiplexers and their outputs are connected to 12 analog inputs of the Arduino. To avoid any well-to-well light contamination, the 96 wells are lit and read one at a time by the software. Each reading, which takes less than one second, is the average of 500 successive acquisitions to reduce noise. The LED matrix and the multiplexers are addressed by digital outputs of the Arduino. LED intensity can be adjusted from the user interface by using the pulse-width modulation functionalities of Arduino digital outputs.

Temperature measurements are achieved by two NTC thermistances (10kOhm 1%, EPCOS ref B57861S0103F040) purchased from RS components (https://fr.rs-online.com). There is one NTC in contact with the bottom of the culture plate and the other one in contact with the lid. Top and bottom heating elements are made by soldering resistive wire (6 Ohm/m from RS components https://fr.rs-online.com) on a PCB plate in a serpentine design around all wells (see Figure S1 in Supplementary Material 4) to ensure homogeneous heating. For one plate, the length of the wire is about 160 cm. Under 12V power supply, the heating power is thus approximately 15W for each heating plate, and the total maximum consumption of the MiniRead is around 3A (the current drained by the LEDs and the overall electronics is negligible as compared to the heating current).

The user interface of the MiniRead (Figure 1A) contains a 4x16 characters LCD display (MC41605A6W-FPTLW-V2 from RS components; https://fr.rs-online.com), six pushbuttons and a control multicolor LED, connected to digital inputs and outputs of the Arduino. It allows a completely autonomous use of the MiniRead without a computer. Two additional LEDS indicate when heating is activated for the bottom and the lid of the culture plate. For additional details, see the user manual in Supplementary Material 1.

During data acquisition, raw values are written in a micro-SD memory card through a cheap standard card reader module for Arduino from Aliexpress (https://fr.aliexpress.com/). The card (FAT32-formatted 8 Gb) can easily be un/plugged from the side of the device box to transfer files to a computer.

### 2. Printed Circuit Board (PCB)

To easily assemble the device, we designed different pieces of PCB to build the plate holder, with soldering pads allowing for a quick assembly without any gluing or screwing step. Additional pieces of PCB are used for the bottom and top plates with their corresponding heating elements. We used EasyEda editor (https://easyeda.com/) to draw the PCB and ordered 5 pieces from JLCPCB (https://jlcpcb.com/), but other export formats (Gerber fabrication files, SVG and PDF images) are available on https://forgemia.inra.fr/gqe-base/MiniRead so it should be possible to have the PCB fabricated by any other company. We did not use any surface mounted components, so soldering of the components can be done with a conventional soldering iron by anybody having experience in electronics.

### 3. Software

The MiniRead contains an embedded software (firmware) written in the Arduino version of the C language. It controls sequencial LED illumination, light intensity data acquisition and averaging, temperature regulation, writing in the SD card, and user interface (writing information on the LCD display, reading keys,…). This software also includes different programs allowing the user to easily carry out growth kinetics recording, calibration, or single plate acquisition. The firmware source code in Arduino C language is available at https://forgemia.inra.fr/gqe-base/MiniRead. It can be compiled and flashed into the Arduino board *via* an USB cable using the free Arduino IDE software available at https://www.arduino.cc/en/software. Beyond updating the firmware, the USB port can also be used for bidirectional serial communication of the MiniRead with a computer for development or debugging purposes, or to display the reading values in real-time on a computer during the growth kinetics.

Once raw data from the MiniRead have been transferred to a computer *via* the SD card, we propose a dedicated R package to help analyzing them. The main functions provided in this package allow to compute and model calibration curves for each of the 96 wells and to analyze growth kinetics experiments to extract different proxies of growth parameters. The package file is available as Supplementary Material 2 and as a release of the MiniRead project at https://forgemia.inra.fr/gqe-base/MiniRead/-/releases.

### 4. Housing case

MiniRead devices must be insulated from external light during operation. Therefore, we designed a plywood box with hinged front cover allowing for easy placing and removing of the plate, top window fitting the user interface, and side openings to access the SD card and the USB connector of the Arduino board, which is necessary to flash new versions of the firmware or to use serial communication. Dimensions of the plywood sheets required to build the housing are available at https://moulon.inrae.fr/materiel_labo/miniread/.

### 5. Physical calibration

To study the response curves of LDRs in all wells of all MiniRead machines, we measured an empty plate with its cover, after placing under the lid zero to 13 layers of neutral optical filter sheet (Frost Diffusion White Lighting Filter Gel Sheets from https://cableandlight.co.uk/) covering all wells. The optical density (OD) of the filter stack is proportional to the number of filter layers, so we used this method to model the non-linear relationship between the raw MiniRead output values and the corresponding number of filter layers, by using the *Calibrate()* function of the MiniRead R package (see tutorial in Supplementary Material 3) which uses the *smooth*.*spline()* function of R.

### 6. Biological calibration

We tested the MiniRead devices with the yeast species *S. cerevisiae* (diploid SK1 and haploid S288c strains), *Candida glabrata*, and *Nakaseomyces delphensis* species (haploid), and with the bacteria *E. coli* (DH5α strain). Liquid cultures for growth kinetics were carried out in 200 μL standard YPD medium for yeasts and LB medium for bacteria, in flat-bottom 96-wells plates (Sarstedt ref. 833924500). Initial inoculation was done at OD=0.01 and growth kinetics were carried out for 48 hours at constant temperature of 30°C (plate bottom) and 32°C (lid) for yeasts, and at 37°C (plate bottom) and 39°C (lid) for the bacteria. LED light intensity was set to 100 (arbitrary units are between 1 and 255) and the MiniReads were programmed to make one acquisition every five (bacteria) or ten (yeasts) minutes (KINETICS program).

To compute calibration curves, we focused only on yeasts. An initial liquid cell culture (600mL) at the plateau stage was centrifuged to remove the medium and re-suspended in 150mL liquid YPD supplemented with 250mM EDTA (to stop growth). This cell suspension was used as the highest point of a linear range of 5 diluted cell concentration standards; the lowest (zero) point was made with pure YPD+EDTA medium. Cell suspensions of each dilution point were carefully homogenized and uniformly distributed in the 96 wells of a flat-bottom culture plate (200mL/well), then measured with the CALIBRATION program of the MiniRead in four technical replicates.

## RESULTS AND DISCUSSION

### 1. Functionalities of the user interface

The MiniRead displays in real-time the temperatures of the bottom and lid of the culture plate and the turbidity value acquired by the LDR sensors for each well coordinate (Figure S2 in Supplementary Material 5). The display also show the set values for the LEDs brightness, the time interval between two data acquisitions in the KINETICS program, and the current output file name (see details in user manual in Supplementary Material 1). File names are composed of a prefix depending on the program (SING for single plate acquisition, CALI for sensors calibration, and KINE for growth kinetics experiments) followed by a number that is automatically incremented before each new experiment. File name extension can be set to MRA, MRB, MRC, etc… to distinguish data produced by different MiniRead units when several devices are used at the same time.

### 2. Accuracy of temperature control

To ensure stable results over time, the MinRead must be powered on several hours before starting the experiment, ideally the day before. Once powered on, it takes about 10-15 minutes to stabilize plate and lid temperatures to respectively 30 and 32°C. The precision obtained on these temperatures in our five prototypes is better than 0.1°C (Figure 2).

**Figure 2.**
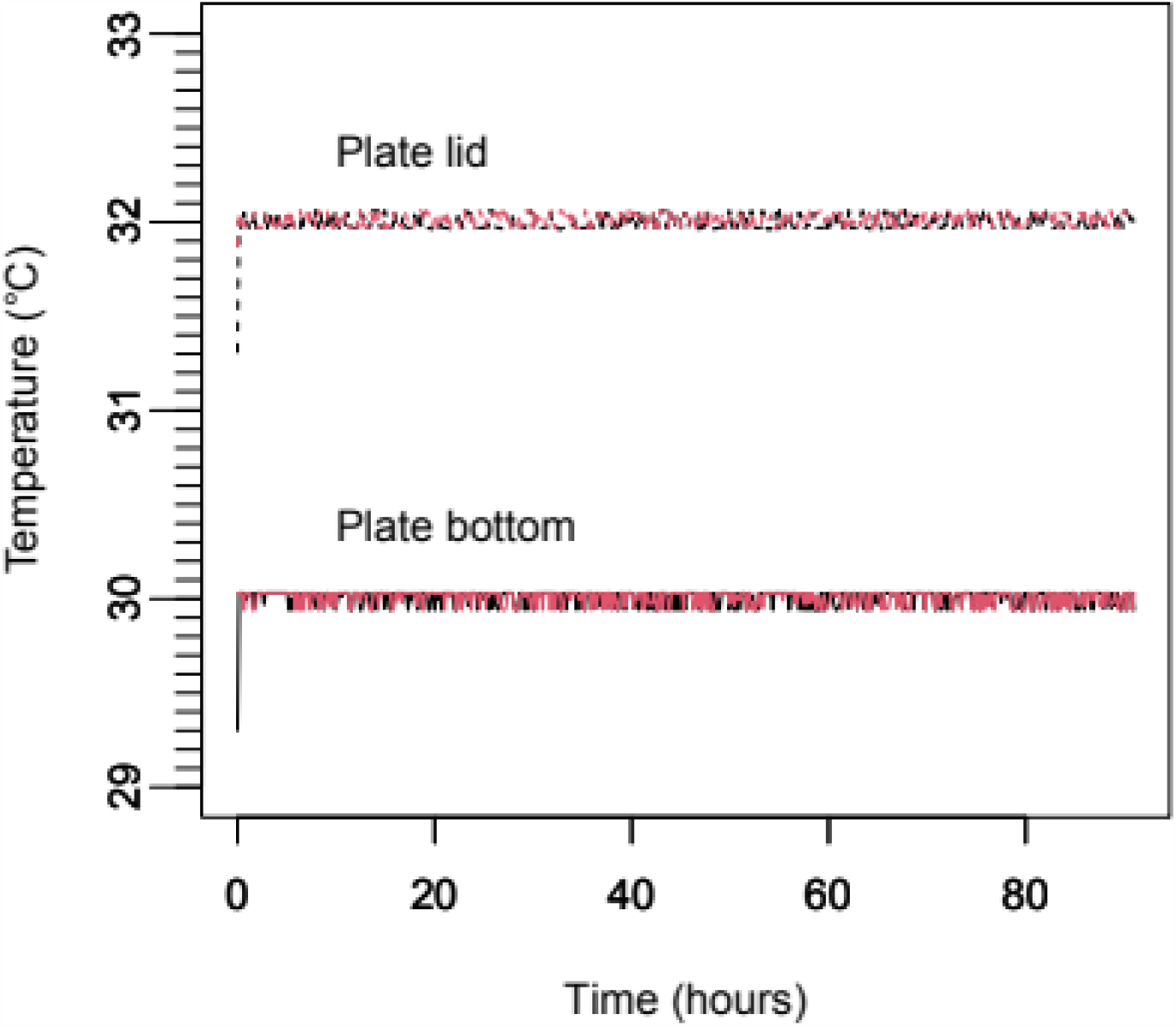
Temperature of the plate bottom (solid lines) and plate lid (dashed lines) during a 4-days growth kinetics. Black (respectively red) curves are temperatures measured just before (respectively after) each data acquisition (every 10^th^ minute).

### 3. Calibration results

The MiniRead raw output values are in arbitrary units between 0 and 1023, which measure the amount of light received by the LDR sensors. However, (1) the response curve of the sensors is not linear and may vary from well to well, and (2) the relationship between measured OD and cell concentration is also non-linear as soon as the concentration becomes high. So we performed two different calibrations: (1) a physical calibration, to model the response curve of each individual LED-LDR pair (see Materials and Methods). The 96 response curves (Figure 3A) were computed by the *Calibrate()* function of the MiniRead R package (see tutorial of the package in Supplementary Material 3). Using these curves, the *ApplyCalibration()* function can transform raw data from any well of any MiniRead unit into a predicted value of the number of filter layers (see Figure 3B) which is proportional to the OD. This value is comparable across all wells, including from different machines. (2) a biological calibration, to model directly the relationship between raw MiniRead output values and cell concentrations, based on standard dilutions of a yeast cell culture (see Materials and Methods). As before, the *Calibrate()* function was used to compute calibration curves (Figure 4A), which can be applied to raw data with the function *ApplyCalibration()*. Figure 4B shows the quality of this biological calibration. As we can see in the X-axis values of Figure 4A, even with a four-times concentrated plateau yeast culture, we are still far below the saturation value of the MiniRead (1023) and the response curves are relatively close to linearity. On the other hand, we see in Figure 3A that when OD is very high (with 13 layers of optical filter), the response curve is clearly non-linear for some wells, but nevertheless, Figure 3B shows that that non-linearity is very well corrected by the *ApplyCalibration()* function of the package, so the MiniRead can be used accurately over its whole dynamic range.

**Figure 3.**
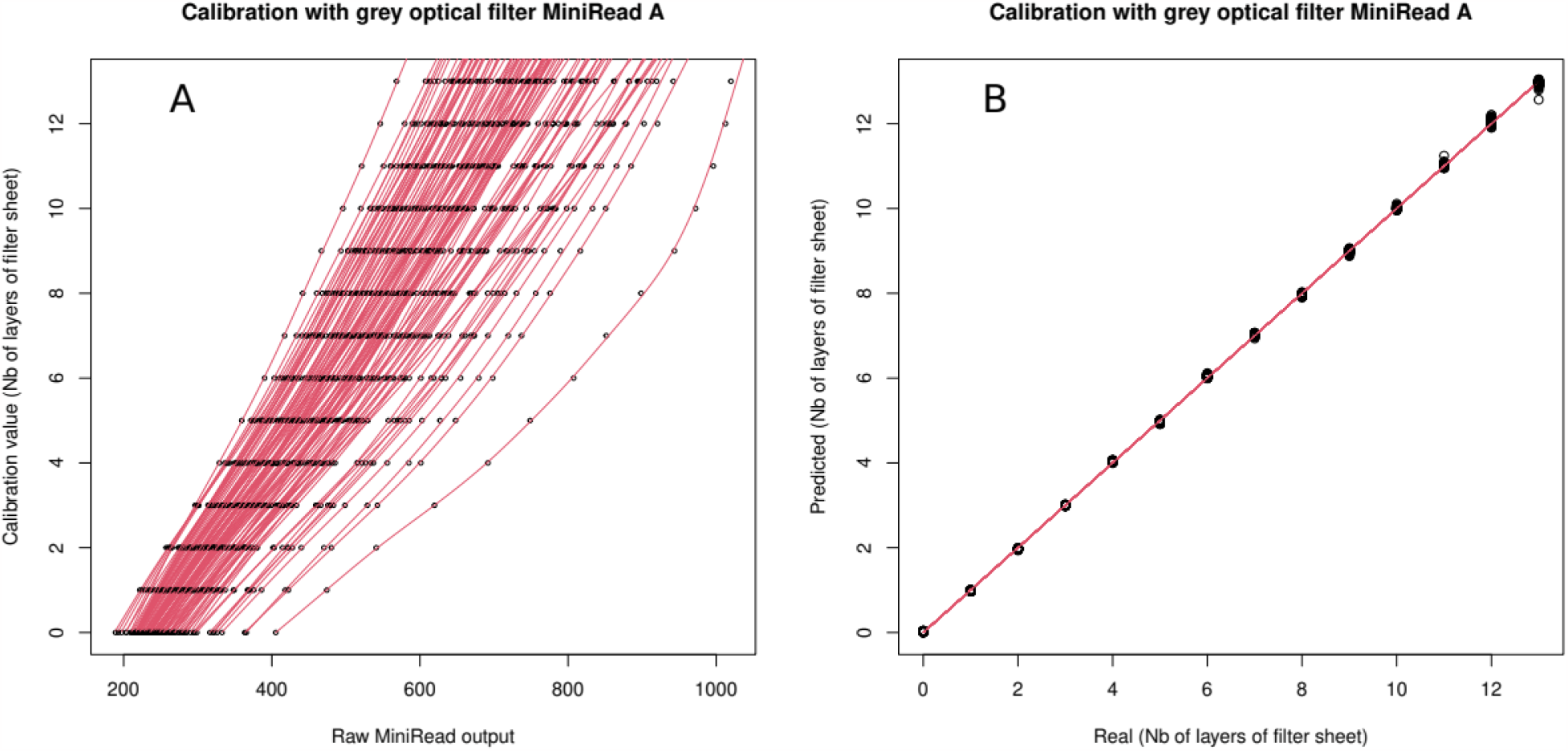
Physical calibration of the 96 LED-LDR pairs using the *Calibrate()* function of the MniRead R package. A: non-linear relationship observed between raw output values produced by the MiniRead (X-axis) when placing increasing numbers of layers of neutral optical filter sheet under the plate lid (Y-axis). Black circles are the real values measured and the red curve is the result of a non-linear fit using splines. B: comparison between real numbers of filter sheets (X-axis) and predicted number (Y-axis) as computed by the *ApplyCalibration()* function from the raw output values, using the non-linear relationship modeled by *Calibrate()*. Black circles are the experimental values and the red line is the result of linear regression.

**Figure 4.**
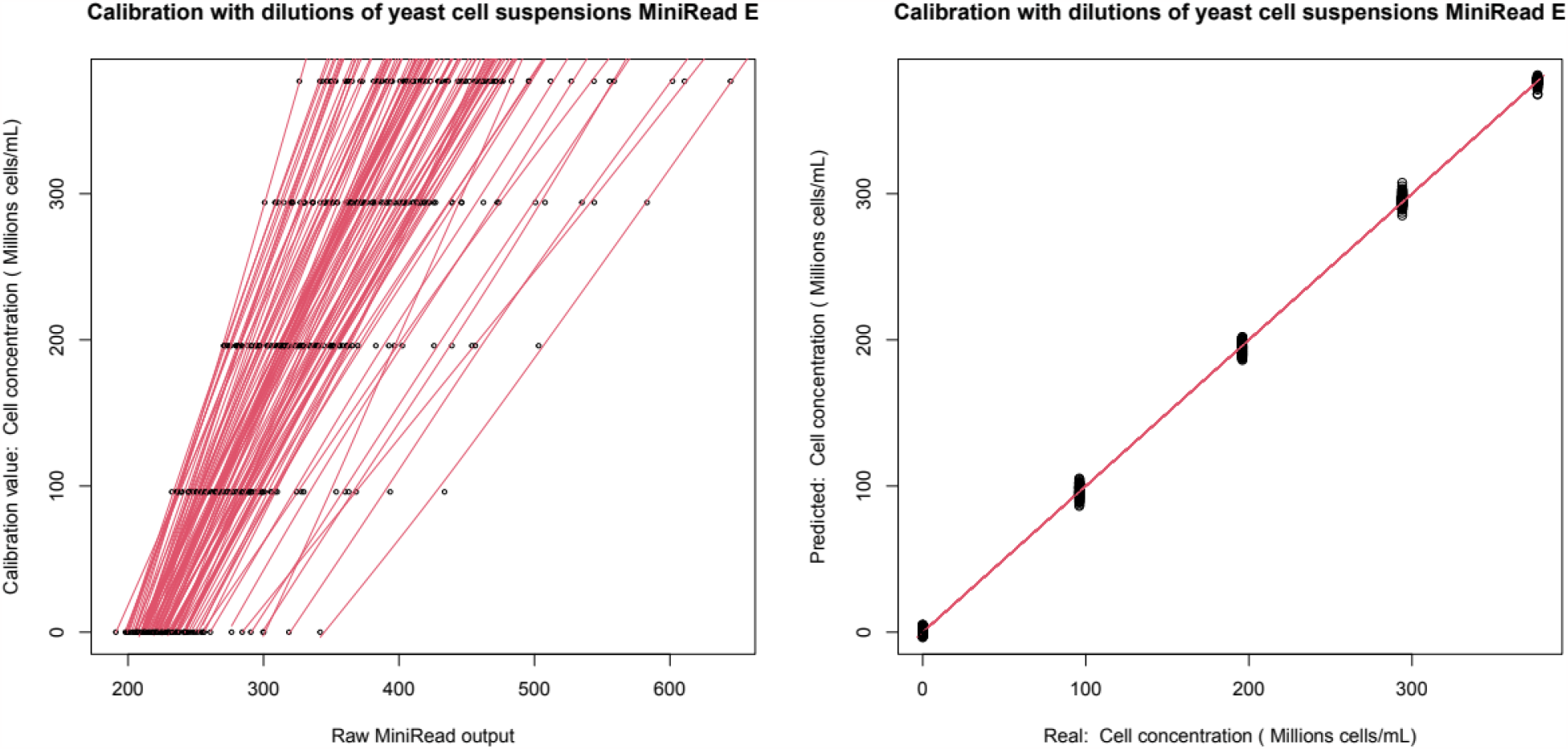
Biological calibration of the 96 wells with the same approach as for Figure 3, except that filter layers were replaced by 96-well culture plates containing a range of yeast cell suspensions of increasing concentrations in YPD medium, from zero to a four-times concentrated culture at the plateau phase (see Materials and Methods).

In case some very dark or very transparent culture media were to be used, it may be useful to change the LED brightness (0 to 255), instead of the default value set to 100, to optimize the dynamic range.

### 4. Growth kinetics analysis

During growth kinetics, the evaporation was not negligible and resulted in a linear increase of the estimated cell concentration over time, even with non-inoculated sterile YPD medium (see Figure S3 in Supplementary Material 6). This was expected since we chose plates designed to ensure sterile gas exchanges allowing aerobic micro-organisms to grow in acceptable conditions. Thus the MiniRead package uses the plateau phase of the cell growth kinetics to model the effect of evaporation, in order to correct this effect when estimating growth parameters. Figure 5 shows a typical growth curve of *S. cerevisiae* over 48 hours, obtained with the MiniRead and analyzed with the *AnalyzeGrowth()* function of the R package using all default parameters (right panel) or with argument flatPlateau=FALSE (left panel). Three different phases of culture growth are automatically identified based on the first and second derivatives of the curve. First, the lag phase from the inoculation until cell concentration reaches a given proportion of the value at the plateau, Then the exponential growth phase with more and more saturation, resulting in an inflection point. Finally the plateau phase where the number of cells is not supposed to change over time. Of course, depending on the organism and conditions used, this does not reflect the complexity of all phenomena controlling the kinetics of cell concentration (e.g. possible switch from fermentation to respiration in yeasts). However, this simple model allowed us to easily extract efficient proxies of three useful parameters of cell growth: the maximum rate of cell division (at the inflection point), the duration of the lag phase, and the cell concentration at the plateau (proxying the carrying capacity). These parameters are automatically estimated from the data before (between parentheses) and after correction for evaporation (Figure 5 right panel). The *AnalyzeGrowth()* function also computes the rate of evaporation (drift) and the signal-to-noise ratio which gives a useful estimation of the physical quality of the results. All parameters computed from the kinetics, without and with correction for evaporation, are also written in a tabulated file to be opened by any spreadsheet software.

**Figure 5.**
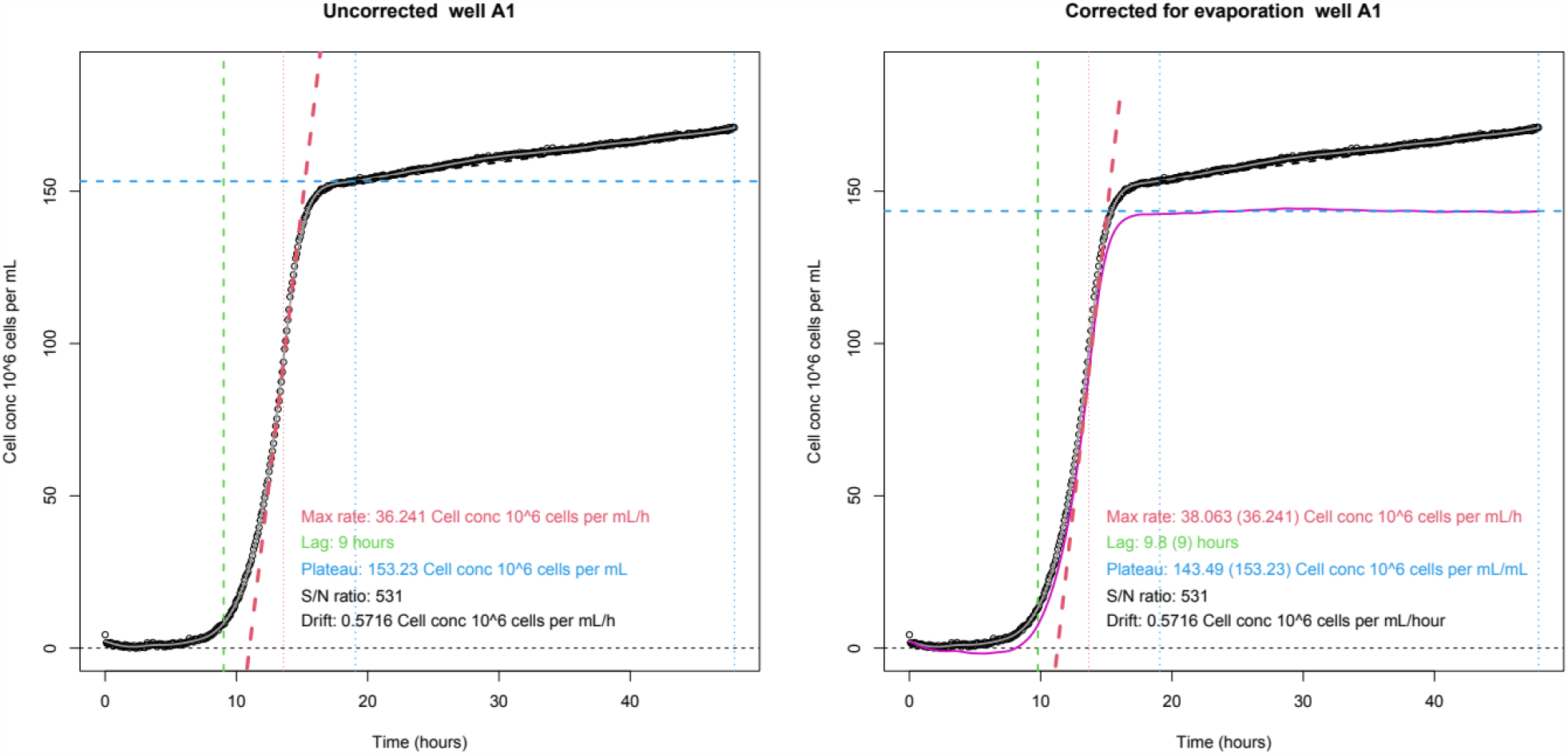
Typical yeast growth kinetics measured with the MiniRead, after automatic analysis of the output file with the function *AnalyzeGrowth()* of the MiniRead R package. Left panel: analysis without correction for evaporation (argument flatPlateau=FALSE). Right panel: with correction for evaporation (default: argument flatPlateau=TRUE). Black circles: experimental measurements transformed into cell concentrations after calibration with titrated cell suspensions. Grey solid line: non-linear fit (splines smoothing) of the experimental points. Black dashed line: linear regression of the plateau phase, used to correct for evaporation. Pink solid line (right panel only): growth curve corrected for evaporation based on the plateau slope. Thick red dashed line: tangent at the inflection point (maximum growth rate) of the uncorrected grey (left panel) or corrected pink (right panel) curve. The other dashed lines indicate: beginning of the plateau (thin blue), value of the plateau (thick blue), end of the lag phase (thick green), time of the inflection point (thin red). Values provided as text in the figures indicate the estimated values of the growth parameters. In the right panel, the values between parentheses refer to the analysis without correction for evaporation.

### 5. Conclusion

The MiniRead provides an efficient and inexpensive solution to measure growth kinetics of yeasts and bacteria in several hundreds of samples in parallel. All required information to home-build these small machines, including associated firmware and a data analysis R package, are freely available under GPL license. So we hope this device can be useful to other research groups, as it is to us.

## Supporting information

User manual

R package

Tutorial for R package

## ACKNOWLEDGEMENTS

Authors contributions: MF developed the hardware, firmware, and R package, built the devices, and wrote the manuscript. AB, FD, MdC, CD performed the wet-lab experiments and helped with the first R scripts. All authors performed yeast growth kinetics experiments and approved the manuscript.

The authors thank Olivier Martin for his help in specifying the temperature regulation algorithm, Franck Gauthier for checking the R package, and Cécile Fairhead, Monique Bolotin-Fukuhara, Youfang Zhou, Killian Métivier, and Xavier Raffoux for helpful discussions and comments on the project. The present work has benefited from funding from the EVOLREC project (ANR-20-CE13-0010). GQE - Le Moulon benefits from the support of the LabEx Saclay Plant Sciences-SPS (ANR-10-LABX-0040-SPS).

## Notes

https://forgemia.inra.fr/gqe-base/MiniRead

https://moulon.inrae.fr/materiel_labo/miniread/

### Competing Interest Statement

The authors have declared no competing interest.

