## Supplementary material for "MiniRead: a simple and inexpensive do-it-yourself device for multiple analyses of micro-organism growth kinetics": User manual

### MiniRead - User manual

#### Simple DIY Microplate Turbidimeter To Measure Growth Parameters of Micro-Organisms

Matthieu Falque

Version 4.1.x

December 20, 2023

**7 How to start with the MiniRead R package** **13**

### 1 Introduction

The MiniRead device is a simple and inexpensive do-it-yourself 96-well turbidimeter designed to measure the interception of white light *via* absorption or side scattering by micro-organism cells in liquid culture medium. Turbidity is automatically recorded in each well at regular time intervals for up to several days or weeks. Output tabulated text files are recorded into a micro-SD memory card to be easily transferred to a computer.
The MiniRead machine is designed to be used together with a R package which allows (1) to compute the non-linear calibration fits required to convert raw readings into cell concentration values, and (2) to analyze growth kinetics output files to automatically estimate growth parameters such as lag time, maximum growth rate, or cell concentration at the plateau.
Detailed information on how to build MiniRead devices is not provided in this manual, but can be found on our web site (`https://moulon.inrae.fr/en/materiel_labo/miniread/`).
All software (firmware and R package) required to use the MiniRead devices are available at ForgeMia (`https://forgemia.inra.fr/gqe-base/MiniRead`).

### 2 General overview

The main features for the MiniRead plate reader are:

- uses flat-bottom 96-wells plates of liquid culture (150 – 200 µL/well) without agitation. The plates are covered with their transparent lid.
- records light interception by the column of cell suspension in each well at regular time intervals,
- accurate control (+/- 0.5°C) of the temperature of the bottom of the culture plate (10 to 45°C),
- independent control of the temperature of the plate lid, which should be kept e.g. two degrees warmer than the bottom to avoid mist condensation – that point is critical for optical readings while keeping sterile conditions in the wells,
- store data autonomously without the need for a computer

The raw turbidity values produced by the MiniRead are not linearly related to optical density or cell concentrations, and each well has a different response curve, due to the heterogeneity and the intrinsic characteristics of the light sensors used. Therefore a step of calibration must be carried out for proper interpretation of the values. All necessary procedures and functions are provided in the embedded firmware as well as in the associated R package, to easily proceed with this calibration (see section 5.1).

#### 2.1 Hardware, software, and versions

The MiniRead hardware requires two distinct pieces of software:

- the internal firmware which needs to be flashed into the Arduino board before the first use, and for every version update
- the MiniRead R package used to analyze the output data from the MiniRead machines (see section 6).

The MiniRead project exists in different versions, involving changes in hardware and/or software. For both, new versions may be released in the future. The version numbering **(x.y.z)** of the MiniRead hardware and software followes the following rules:

- **x** is the version number of the hardware. A change in x means that printed circuit board and/or electronic components have been modified
- **y** is the major software version number. The firmware (C software embedded in the device) and the R package used for data calibration and analysis MUST have the same major version number y to ensure that the files generated by the devices can be analyzed with the R package

- **Z** is the minor software version number. Indicates minor bug fixes or improvements. Changes in values of z do not affect the compatibility between firmware and R package

#### 2.2 Connection of the MiniRead to a computer *via* USB

Using the MiniRead does not require a computer, except in two particular cases:

- to update the internal firmware which is flashed into the Arduino board. To do so, use the Arduino IDE and follow its instructions.
- to display additional informations during the run of a program. This is not useful for normal use of the device but can be very useful for debugging purposes.

#### 2.3 Case of several MiniRead units used in parallel

MiniRead devices are cheap and most companies which fabricate the printed circuit boards will not produce less than five units. Moreover, the time required to build five MiniRead devices in parallel is much less than five times the time to build one. So we assumed that most users will build more than one machine to achieve high-throughput analyses, and so we designed the software to easily handle work with several devices in parallel.
To do so, each device is given a unique name under the form of a single-letter label (e.g. 'A', 'B',...). This label is set in the beginning of the source code which has to be flashed into the Arduino board before being able to use the MiniRead. Each machine will then:

- display its label when booting
- write its label in the column 'device' of the output data files (in all lines)
- end its data file names by the extension '.MRA' if the label is 'A', '.MRB' if the label is 'B', etc...

The MiniRead R pachake is designed as much as possible to handle such multiple devices.

#### 2.4 Switching the MiniRead ON/OFF

There is no main ON/OFF switch. To switch off the device, simply remove the Jack plug of the power supply. The power supply must output a good-quality constant DC 12V and must be able to provide at least 3A of constant current.
Since the temperature of the sensors and the internal circuitry may influence the measurements, it is stronglyh advised to power on the device several hours before using it, or even better the day before, so its temperature gets perfectly stablilized. As soon as the devices is powered, it will start warming-up at the set temperature (see section 4.3).

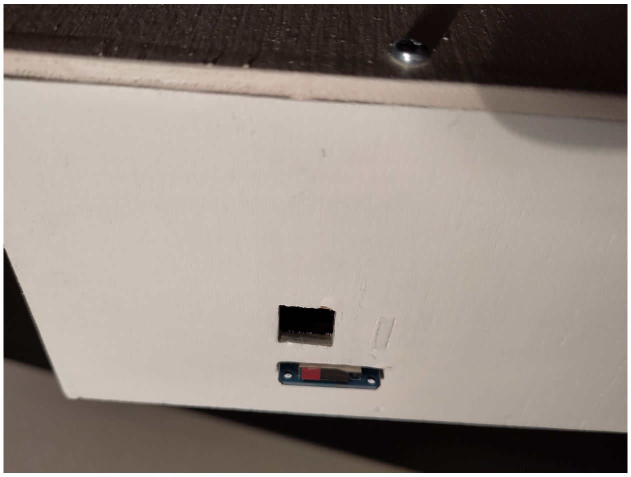

Figure 1: External SD card.

#### 2.5 Placing and removing the micro-SD card

When the MiniRead is doing turbidity acquisitions, the data are saved on the micro-SD card which is plugged on the left side of the device (see Figure 1 page 5). Press on the extremity of the card to insert or release it. Any small-capacity SD card can be used (16Gb is much more than enough !)
**CAUTION** : Never place or remove the SD card while the MiniRead is in action, it may corrupt data or even the card itself. Do it only while the interface displays 'IDLE'.
SD cards must be formatted as FAT32.

#### 2.6 Placing and removing plates

To place a culture plate in the device for reading,

- lift the front cover part up (using both hands...)

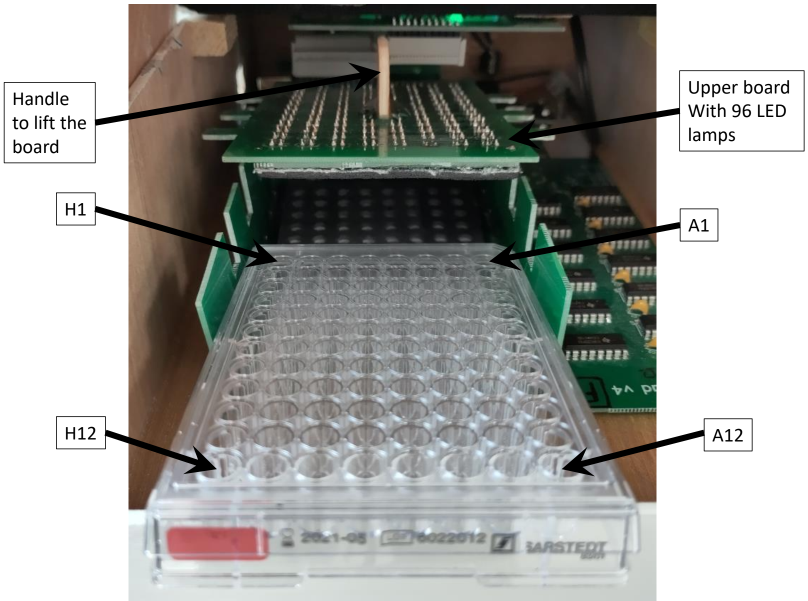

Figure 2: Upper board lifted to introduce the culture plate.

- lift the upper electronic board (with the 96 LED lamps) and push it slightly back until it holds on top of the plate housing, while leaving the space to introduce the plate (see Figure 2 page 5). To do so, hold the lid carefully by the wooden handle, lift it softly upwards without forcing (it would do nothing better than making it even more difficult to lift).
- gently place the plate in its housing, be careful to avoid splashing the medium from the wells. Check carefully that the plate is in direct contact with the bottom of the housing. Check also carefully the correct orientation of the plate, with 'A1-A12' row on your right side and 'H1-H12' on your left side.

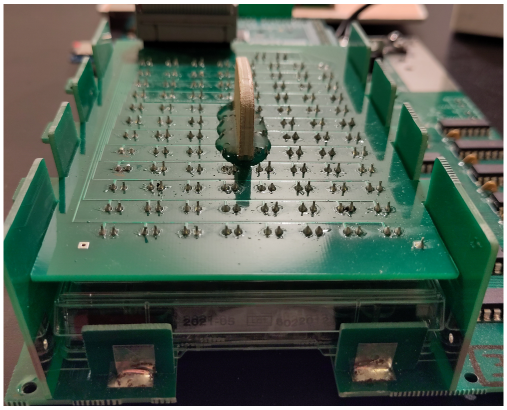

Figure 3: Culture plate correctly in place for reading.

- pull the upper electronic board slightly forward so it can fit again in its lateral slides, and gently press to slide it down. Make sure that the board comes in close contact to the lid of the culture plate (see Figure 3 page 6).
- close the front cover of the device

### 3 Output files

#### 3.1 File names

MiniRead data file names are made of three parts:

- a prefix (KINE, CALI, SING, or PMAP) which indicates the program used to produce the data,
- a three-digit number which is automaticallhy incremented each time a program is run by the MiniRead. This choice was made for traceability reasons, it is strongly advised to NEVER change these file names (although it is possible to do so) to avoid overwriting existing data.
- an extension made with '.MR' and a single last letter which indicates the name (label) of the MiniRead device being used. This is very useful when several MiniRead devices are run at the same time. That label (last letter) is also written by each device in the column 'device' of the data files (see 2.3).

#### 3.2 File format

The data files produced by any of the programs CALI, KINE, and SING of the MiniRead are simple tabulated text files (space-separated) with exactly the same format made of the 105 following columns:

- Col1: 'hours' (float, in hours) time after launching the program,
- Col2: 'bright' (integer between 1 and 255): brightness of the 96 white LED lamps,
*****CAUTION**** MUST be the same value as that used for calibration

- Col3: 'thBottom1' (float, in degrees Celsius): temperature of the bottom of the plate, measured just before reading the plate
- Col4: 'thLid1' (float, in degrees Celsius): temperature of the lid of the plate, measured just before reading the plate
- Col5: 'device' (one letter): label of the MiniRead machine. Is automatically filled by the MiniRead.
- Col6: 'rep' (integer): index of the replicate (used only for the CALIB program). Is not used by the MiniRead device, but can be useful for subsequent analyses with the MiniRead R package.
- Col7: 'calib' (integer): index of the calibration point (used only for the CALIB program), for instance: index of the point of a dilution series of cell suspensions, or numbers of optical filter sheets superimposed. This value is automatically incremented by the MiniRead device before each next sample of a calibration series is read.
- Cols8-103: 'W1' to 'W96': well (in the order: A1..A12, B1..B12,.., H1..H12)
- Col104: 'thBottom2': (float, in degrees Celsius): temperature of the bottom of the plate, measured just after reading the plate
- Col105: 'thLid2': (float, in degrees Celsius): temperature of the lid of the plate, measured just after reading the plate

Such files can be easily viewed with any spreadsheet program or directly used with the different functions of the MiniRead R package.

### 4 User interface

#### 4.1 Idle display

Just after switching the MiniRead on, it turns into Idle mode. In that mode, it displays in real time the temperature of the bottom of the plate (PLATE) and of its top (LID) (see Figure 4 page 8). In Idle mode, the MiniRead alternatively switches between two phases:

- control of plate and lid temperatures. During this step, the multicolor LED flickers rapidly.
- warming-up of the 96 LED-sensor pairs. To do so, while in Idle mode, the MniRead never stops doing turbidity acquisitions at regular time intervals, to maintain the electronic components at stable temperature. During such readings, the multicolor LED makes short flashes as explained in section 4.2.

#### 4.2 Control LEDs

The MiniRead interface has three control lamps (LEDs, see Figure 4 page 8) which provide the following informations:

- Heating LEDs (2 red LEDs on top, named 'PLATE' and 'LID')
  These LEDs turn ON when the plate and lid heating elements are powered.
- Multi-colored control LED
  This LED gives different types of informations to help understanding what the MiniRead is doing:
  - very quick flickering indicates that the MiniRead is measuring and adjusting plate bottom and lid temperatures, and the color indicates if the plate is too cold (rather blue), too hot (rather red), or just at the right temperature (green).

– short flashes at regular time intervals (1-2 per second) indicate that turbidity measurements is being performed. This happens during data acquisition in CALI, KINE, SING, or PMAP programs (then, the color indicates the well being measured, from blue (A1) to red (H12)), but also during the Idle phase, while regularly warming-up the 96 lEDs and sensors (then the color stays white).

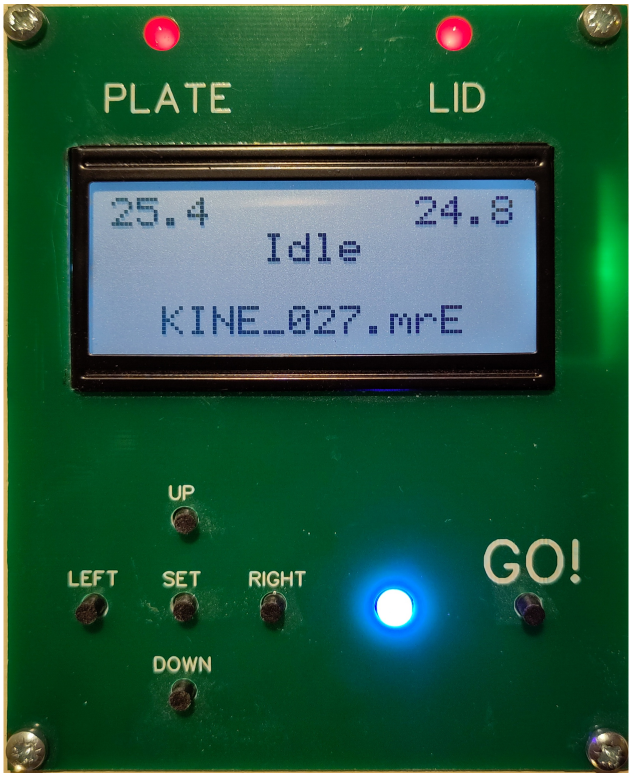

Figure 4: Interface board in Idle mode.

#### 4.3 Parameters setting

The MiniRead stores some parameters in its internal EEPROM memory, which is persistent event when the device is unplugged. To view or change the general settings of the MiniRead, press SET to enter the settings mode (see Figure 5 page 9).
The display shows the current values of the parameters settings. To change any of these values, use the LEFT and RIGHT buttons to navigate from one setting to another, and the TOP and DOWN buttons to increase or decrease the value (each digit separately). Then press SET again to validate the new values and/or come back to the Idle menu. The parameters are listed below (from top to bottom of the screen):

- set temperatures for bottom (PLATE) and top (LID) of the culture plate.
  ****IMPORTANT**** it is strongly advised to set the lid temperature two degrees Celsius higher than the temperature of the plate bottom, to avoid any condensation of water under the lid, which would completely alter the readings.
- name of the current program
- brightness value (from 1 to 255) for the 96 LED lamps used to measure turbidity. The optimal value may depend on the type of culture medium and the type of LEDs used to builf the device. The value 100 can be a good start.

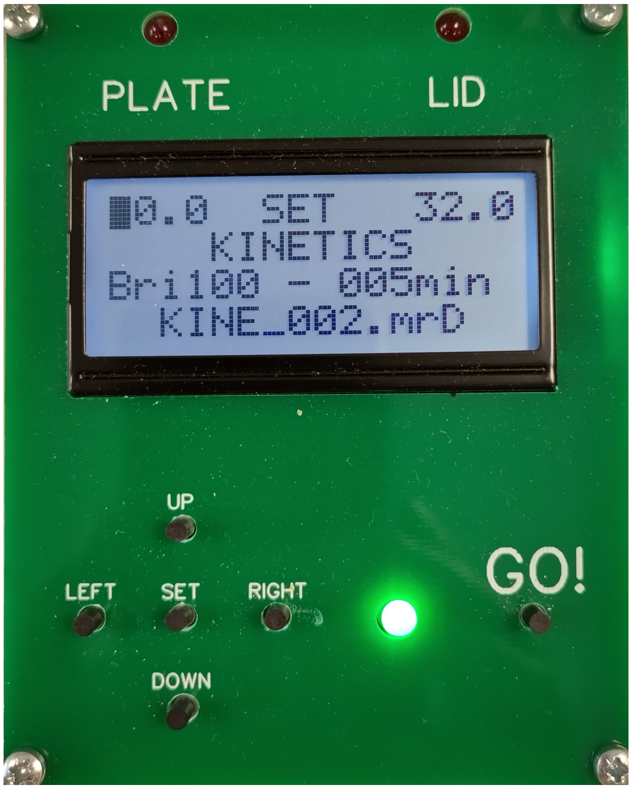

Figure 5: Interface in settings mode.

- time interval (in minutes) between two successive acquisitions during the KINE program (recording growth kinetics)
- name of the current data file (see details on file format and names in section 3.2).

### 5 Programs

The MiniRead has four different programs:

- `CALIBRATION`: Used to produce calibration curves which will be applied to the output values of the MiniRead.
- `KINETICS`: Used to record growth kinetics curves by measuring turbidity every fixed interval of time (minimum 5 minutes, or even less if the temperature control is not critical).
- `SINGLE SCAN`: Does only one measurement of each well of the plate (takes approx. 50 seconds). Used for checks or debugging purposes.
- `PLATE MAP`: Same as `SINGLE SCAN`, but outputs the data under the form of a plate map instead of a tabulated file. Used only for debugging purposes, particularly when the MiniRead is connected to a computer serial terminal in USB mode.

#### 5.1 Running program `CALIBRATION`

Calibration may be done with plates containing for instance:

- a series of dilutions of a well-titrated cell suspension, which will subsequently allow to convert raw output values into cell concentrations with the MiniRead R package. Since the response curve of the sensors is not perfectly linear, it is advised to use at least 4 or 5 points for the dilution series to capture the shape of the curve. For exemple, with *S. cerevisiae* cells, this may be achieved by adding 0.25M EDTA (to stop growth) to a culture concentrated at 4x the plateau,and then making serial dilutions of this suspension. In that case, using the calibration curves obtained, it will be possible to convert the MiniRead output values into cell concentrations as long as the same species/strain/medium is used.

- increasing numbers of layers of neutral optical filter paper, which will subsequently allow to convert raw output values into Optical Density (OD) values with the MiniRead R package.

To carry out calibration, first prepare a series of plates as described above (e.g. 5 plates, one for each point of the calibration curve) with ***increasing*** known values of turbidity. All wells of a given plate must have exactly the same turbidity.
After letting the device warm up (see section 2.4) and setting up the parameters, choose the `CALIBRATION` program as described in section 4.3, then return to the Idle mode with SET, and then:

- press GO! The multicolor LED lights up yellow
- follow the instructions on the screen and introduce the plate as described in section 2.6,
- press GO! again. The multicolor LED makes short blue-to-red flashes as each well is read (see section 4.2).
- when the scanning of the plate is completed, follow the instructions of the screen to remove the plate and replace it by the next one of the calibration series. The sample number on the display will be automatically incremented.
- do the same with all remaining plates until all points of the calibration curve have been measured
- when the last plate has been read, terminate the program by pressing simultaneously SET and GO! until the MiniRead exits the program CALIBRATION and comes back to Idle mode.

During each acquisition step, the screen will display in real time the (raw) values of turbidity measured. The value of the column 'calib' of the output file, which contains the index of the calibration point to be measured, will be automatically incremented after each plate has been read.

#### 5.2 Running program `KINETICS`

After letting the device warm up (see section 2.4) and setting up the parameters, choose the `KINETICS` program as described in section 4.3, then return to the Idle mode with SET, and then:

- press GO!
- place the plate as described in section 2.6,
- press GO! again to start the recording

During each acquisition step, the screen will display in real time the (raw) values of turbidity read. Reading steps (approx 50 seconds) will be followed by waiting steps (duration depending on the parameter set, see section 4.3), during which the screen will display the remaining time before the next data acquisition.
When the desired time of the experiment is elapsed (kinetics are possible over up to weeks if the plates are properly sealed against evaporation), press simultaneously SET and GO! to stop the program until the MiniRead turns back to Idle mode.
You can then remove the SD card from the device and bring it to your computer for data analysis with the MiniRead R package.

#### 5.3 Running program `SINGLE SCAN`

This program is mainly used for debugging purposes. It will carry out a single scan of the plate and write a single-line output file.

##### 5.4 Running program `PLATE MAP`

This program is used to be able to quickly visualize a 2D mapping of the values within a 96-wells plate. It is mainly useful when the MiniRead is connected to a computer *via* the USB port of the Arduino board and using the Arduino serial terminal.

#### 6 MiniRead data analysis using the MiniRead R package

The MiniRead R package contains functions to calibrate the MniRead optical plate reader / turbidimeter and automatically analyze growth kinetics data produced by the device. The two main functions are `Calibrate()` and `AnalyzeGrowth()`.

##### 6.1 Analyzing calibration data

After running calibration plate analyses with the `CALIBRATION` program, as described in section 5.1, the output files (named `CALI_xxx.MRx`) can be used to model the non-linear relationship between the raw turbidity values produced by the MiniRead and real values of cell concentrations or optical density (OD). This is done automatically by the function `Calibrate()` (see all details of use in the R package documentation). You will have to give the different calibration values (e.g. vector of the values of cell concentration of each point of the dilution series) as an argument of the function. The returned object is a list of spline parameters which can subsequently be used to transform raw MiniRead kinetics data into real cell concentration (or OD) values using the function `ApplyCalibration()`. Applying this calibration is also done automatically when using the function `Analyzegrowth()`.

##### 6.2 Analyzing kinetics data

The function `Analyzegrowth()` (see all details of use in the R package documentation) allows to generate kinetics graphs from the raw output files (named `KINE_xxx.MRx`) of the `KINETICS` program, obtained as described in section 5.2. The graphs can be written in a pdf file. In addition, each kinetics curve is automatically analyzed to detect three successive phases: the lag phase before exponential growth, the exponential growth phase, and the plateau phase. Typical parameters are estimated for each of these phases, and these parameters are also written in a tabulated text file.

##### 6.3 Mostly used functions of the R package

The complete liste of functions of the MiniRead R package should be found in the documentation of the package. The main ones to be used to perform usual growth analyses are the following:

- `ReadMiniReadData()`: reads MiniRead output files into a data frame. `WriteMiniReadFile()`: writes files compatible with MiniRead output files from a data frame (allows to re-work the data with other scripts)
- `Calibrate()`: computes calibration curves from data obtained with the `CALIBRATION` program of the MiniRead, as explained in section 5.1.
- `ApplyCalibration()`: applies previously computed calibration curves to a Miniread output file to convert the value into true OD or cell concentration measurements.
- `AnalyzeGrowth()`: analyzes ouput data from the `KINETICS()` program of the MiniRead to compute growth parameters, as explained in section 6.2.
- `WellNames2WellCoords()`: converts well names as written in the MiniRead output files ('W1'... 'W96') into well coordinates of the form 'A1'...'H12'.
- `WellCoords2WellNames()`: converts well coordinates of the form 'A1'...'H12' into well names as written in the MiniRead output files ('W1'... 'W96').

- `HomogeneityPlot()`: draws heat-maps of the values of growth parameters across the different positions within a culture plate.
