## Supplementary material for "MiniRead: a simple and inexpensive do-it-yourself device for multiple analyses of micro-organism growth kinetics": R package: Create_Package_MiniRead.R

```
rm(list=ls())

 pathToPkg <- path.expand("/OCT/ACEP/EVOLREC/42_MiniRead/MiniRead/")
 pathToPkg <- path.expand("/home/mfalque/DATA/GIT_DEV/MiniRead/")

 if (require(MiniRead)) detach(package:MiniRead, unload=TRUE)    # Detach package if previously loaded
if(! require("devtools"))  install.packages(pkgs="devtools")
if(! require("roxygen2"))  install.packages(pkgs="roxygen2")
if(! require("testthat"))  install.packages(pkgs="testthat")

pkgName <- "MiniRead"
pkgFields <- list()
pkgFields[["Package"]] <- pkgName
pkgFields[["Version"]] <- "4.1.3"
pkgFields[["Title"]] <- "MiniRead"
pkgFields[["Description"]] <- "Set of R functions to calibrate the MiniRead turbidimeter and automatically analyze growth kinetics of microorganisms."
pkgFields[["Authors@R"]] <- c(person(given="Matthieu", middle=NULL, family="Falque", email="", role=c("cre","aut"), comment="ORCID=0000-0002-6444-858X"))
pkgFields[["License"]] <- "GPL-3"
stopifnot( dir.exists(pathToPkg))

#  Builds the package
file.remove(paste0(pathToPkg, "/NAMESPACE"))
usethis::create_package(path=pathToPkg, fields=pkgFields, rstudio=TRUE, open=FALSE)       # Create package
devtools::document(pkg=pathToPkg)                               # Format Doc of packages
BuildPkgPath <- devtools::build(pkg=pathToPkg, path=pathToPkg)  # Build package as a tar.gz file

###########  DO NOT FORGET TO ACTIVATE THE FOLLOWING AT LEAST ONCE
  devtools::check(pkg=pathToPkg)                    # Check package
########### 

# Tests installation
detach("package:MiniRead", unload=TRUE)
pkgFileName <- paste0(pkgFields[["Package"]], "_", pkgFields[["Version"]], ".tar.gz")
install.packages(paste0(pathToPkg,"/",pkgFileName), repos=NULL)
library(MiniRead)
help(package="MiniRead")
rm(list=ls())
```
