## Supplementary material for "MiniRead: a simple and inexpensive do-it-yourself device for multiple analyses of micro-organism growth kinetics": R package: MiniRead_V4.ino

```
/***********************************************************************
 * Copyright (c) 2023 Matthieu Falque <>.
 *
 * This file is part of MiniRead.
 *
 *  MiniRead is free software: you can redistribute it and/or modify
 *  it under the terms of the GNU General Public License as published by
 *  the Free Software Foundation, either version 3 of the License, or
 *  (at your option) any later version.
 *  MiniRead is distributed in the hope that it will be useful,
 *  but WITHOUT ANY WARRANTY; without even the implied warranty of
 *  MERCHANTABILITY or FITNESS FOR A PARTICULAR PURPOSE.  See the
 *  GNU General Public License for more details.
 *
 *  You should have received a copy of the GNU General Public License
 *  along with MiniRead. If not, see <http://www.gnu.org/licenses/>.
 *
 ***********************************************************************/
//######################################################################
//#####                         MiniRead                          ######
//######################################################################

#include "LiquidCrystal.h"
#include <SPI.h>
#include <SD.h>
#include <EEPROM.h>
//#include "TM1637Display.h"
//#include <SdFat.h>

#define DEF_VERSION          "v4.1.3"
#define THIS_MINIREAD        'A'
#define THERMISTOR_NOMINAL   10000   // resistance at 25 degrees C 
#define TEMPERATURE_NOMINAL  25      // temp. for nominal resistance (almost always 25 C)
#define NB_CAPTURES_TH       200     // how many samples to take and average, more takes longer
#define NB_CAPTURES_LIGHT    500    // how many samples to take and average, more takes longer
#define B_COEFF_TH           3988    // The beta coefficient of the thermistor (usually 3000-4000)
#define SERIES_RESISTOR_TH   10000   // the value of the 'other' resistor with the thermitors
#define NB_TEMP_STORED       4       // memory of the last steps to estimate the slope
#define     CYCLE_TIME       50    // duration of one cycle (approx X 60milliseconds)
#define        TAU_HEAT      20      // characteristic time (in CYCLES) of the control of heating
#define        TAU_MAX       5       // characteristic time (in CYCLES) of the system (reflects thermal inertia) if close to target
#define CLOSE_TEMP_RANGE     1.0     // temperature range around target beyond which TAU must decrease
#define MAX_PAR_TESTS        4       // number of different values to scan when testing temperature control parameters
#define NB_THS               3       // number of thermistors
#define NB_PARAMS            25      // nb bytes in EEPROM for brightness, kineDelay, setThPlate, setThLid, kineFileIndex, scanFileIndex, caliFileIndex, mapFileIndex
#define NB_PROTOCOLS         4       // nb of different protocols (kinetics, single scan, Mapping...

#define PIN_BTleft      28  
#define PIN_BTright     29  
#define PIN_BTdown      30  
#define PIN_BTup        31  
#define btnNONE          0
#define btnLEFT          1
#define btnRIGHT         2
#define btnDOWN          3
#define btnUP            4
#define btnSET           5
#define btnGO            6
#define btnCANCEL        7

#define PIN_LDRext       A14   // LDR for ambiant light level
#define PIN_LED1R        44   // LED between CANCEL and LEFT
#define PIN_LED2G        45  // LED between LEFT and RIGHT
#define PIN_LED3B        46  // LED between RIGHT and VALID
#define PIN_OUT3         36  // output via transistor BD679, also LEDext
#define PIN_OUT4         37  // output via transistor BD679

/*
NOTES FOR MiniREAD BOX

W <- 180  #  bottom plate width (interior)
D <- 320  #  bottom plate depth (interior)
T <- 3.6    #  thickness of wood
TB <- 10  #  thickness of bottom plate
H <- 80   #  height (interior)
FH <- 30  #  height (interior) of the front side
DF <- 150 #  depth (interior) of the fix part of the top

Bottom <- c(W, D)
Left <- c(D, H+TB)
Right <- Left
Rear <- c(W+2*T, H+TB)
Front <- c(W+2*T, FH+TB)
TopFix <- c(W+2*T, DF+T)
TopCover <- c(W+4*T, (D-DF)+2*T)
LeftCover <- c((D-DF)+T, H+TB)
RightCover <- LeftCover
FrontCover <- c(W+4*T, H+TB)

Bottom
Left
Right
Rear
Front
TopFix
TopCover
LeftCover
RightCover
FrontCover

> Bottom         180 x 320   ctp 10mm
> Left-Right     320 x 181   ctp 3,6mm
> Rear-Front     187 x 181   ctp 3,6mm
> TopFix         187 x 154   ctp 3,6mm
> TopCover       194 x 177   ctp 3,6mm
> LeftRightCover 174 x 181   ctp 3,6mm
> FrontCover     194 x 181   ctp 3,6mm


*/
//*********************************************
//***************  GLOBALS  *******************
//*********************************************
// char VERSION[7]=DEF_VERSION;
uint8_t PINS_TEMP[NB_THS] = {A13, A12, A15};   //  A13=plate bottom,  A12=plate lid,   A15=external (ambiant temp)
uint8_t PINS_HEAT[NB_THS] = {35, 34};

uint8_t PINS_MPX_ADR[3] = {49, 48, 47}; // address bus of 4051 MPXers (for analog signal from LDRs)
uint8_t PINS_LED[8] = {43,42,41,40,39,38,33,32}; // return from LED rows
uint8_t PINS_PWM[12] = {3, 2, 5, 4, 7, 6, 9, 8, 11, 10, 13, 12}; // PWM signal to modulate light intensity of LED columns
uint8_t PINS_LDR[12] = {A11,A10,A9,A8,A7,A6,A5,A4,A3,A2,A1,A0};   // analog input from LDR 8-PLEXes to pins A0-A11 
uint8_t PINS_SD[4] = {50, 51, 52, 53};   // to SD card reader 50=MISO, 51=MOSI, 52=SCL, 53=CS
uint8_t PINS_LCD[6] = {27, 26, 25, 24, 23, 22};
uint8_t COLOR_R[10] = { 0, 255,   0,   0, 220, 255, 255,  100 };
uint8_t COLOR_G[10] = { 0,   0, 255,   0, 255,  50, 160,   50 };
uint8_t COLOR_B[10] = { 0,   0,   0, 255,  80,   0,   0,  100 } ;
//                      N    R    G    B    W    O    Y    P
//                      0    1    2    3    4    5    6    7

//typedef struct {
//  float gain;          // gain value to apply to the light intensity readings
//  float offset;        // offset value to apply to the light intensity readings
//  float brightness;  // LED PWM value to be used for OD measurement
//} S_LDR;
//S_LDR ldr[96];

uint8_t defaults[NB_PARAMS]={3,0,0,  3,2,0,   0,  1,0,0,  0,1,0,  0,0,1,  0,0,1,  0,0,1,  0,0,1};
//                           30.0    32.0     0    100      10        1       1       1       1
//                          Plate     Lid   Prog    Bri   Delay    KINE    SING    CALI     MAP

typedef struct  {
  float setTh=0;                      // temp to be reached
  float tau;                        // weight of slope
  float slope;                      // slope of temperature ~ cycles regression
  float intercept;                  // intercept of temperature ~ cycles regression
//  float offset;                     // value to be added to the calculated temperature to get zero on ice
//  float calib;                      // factor to be multiplied to the calculated temperature to get calibration value
  float heat;                       // duration of heating (relative to one)
  float th;                         // temp of Control dish, in ° Celsius
  float storedTh[NB_TEMP_STORED];   // stores the last NB_TEMP_STORED temperatures measured
  bool  isConnected;                // true if a unit is connected to this channel
} S_THER;
S_THER ths[NB_THS];                 //  th[1] for LID thermistor, th[0] for bottom thermistor, th[2] for external temperature

File dataFile;

uint8_t iLine=0, iCol=0, btn=btnNONE;
float reading=0, bright=0, kinDelay=0;
char odFile[13]="xxxxxxxx.txt";
bool USE_SD_CARD = true;
bool USE_HEATING = true;

LiquidCrystal lcd(PINS_LCD[0], PINS_LCD[1], PINS_LCD[2], PINS_LCD[3], PINS_LCD[4], PINS_LCD[5]);

//***************************************************
//********************  SETUP  **********************
//***************************************************
void setup() {
  Serial.begin(9600); while (!Serial) ;
  delay(200);
  digitalWrite(LED_BUILTIN, LOW);
  Serial.println();Serial.println(); Serial.print("MiniRead "); Serial.println(DEF_VERSION);
  for (uint8_t i=0; i<3; i++) { pinMode(PINS_MPX_ADR[i],OUTPUT); digitalWrite(PINS_MPX_ADR[i],LOW); }
  for (uint8_t i=0; i<8; i++) { pinMode(PINS_LED[i],OUTPUT); digitalWrite(PINS_LED[i],HIGH); }
  for (uint8_t i=0; i<12; i++) { pinMode(PINS_PWM[i],OUTPUT); analogWrite(PINS_PWM[i],0); }
  pinMode(PIN_LED1R,OUTPUT); pinMode(PIN_LED2G,OUTPUT); pinMode(PIN_LED3B,OUTPUT); 
  pinMode(PIN_BTleft, INPUT);  pinMode(PIN_BTright, INPUT); pinMode(PIN_BTdown, INPUT); pinMode(PIN_BTup, INPUT); 
  pinMode(PINS_HEAT[1],OUTPUT); digitalWrite(PINS_HEAT[1],LOW);
  pinMode(PINS_HEAT[0],OUTPUT); digitalWrite(PINS_HEAT[0],LOW);
  lcd.begin(16,4);
  lcd.clear();
 
 // for (uint8_t iPar=0; iPar<NB_PARAMS; iPar++) EEPROM.put(iPar, defaults[iPar]);
//  EEPROM.put(22, 0);  EEPROM.put(23, 0);  EEPROM.put(24, 1);  
 
  lcd.setCursor(0,0); lcd.print("****************");
  lcd.setCursor(0,1); lcd.print("*  Mini Read  *");
  lcd.setCursor(13,1); lcd.print(THIS_MINIREAD);
  lcd.setCursor(14,1); lcd.print(" *");
  lcd.setCursor(0,2); lcd.print("*    ");
  lcd.setCursor(5,2); lcd.print(DEF_VERSION);
  lcd.setCursor(12,2); lcd.print("   *");
  lcd.setCursor(0,3); lcd.print("****************");
  btn=btnNONE;
  while (millis()<10000 & btn==btnNONE) btn=Read_external_buttons();
  lcd.clear();

  if (!USE_HEATING) Serial.println("Use of heating disabled");
  if (USE_SD_CARD) {
    uint8_t ok=false;
    while (!ok) {
      ok=SD.begin(PINS_SD[3]);
      if (!ok) Error_flash(3,1,100);
    }
 //   Sd2Card sdCard;
 //   Sd_test(sdCard);
  }
  ths[0].setTh=0; ths[1].setTh=0; bright=0; kinDelay=0;
  for (uint8_t i=0; i<3; i++) {
    ths[0].setTh += myPow(10, 1-i) * EEPROM.read(0 + i);
    ths[1].setTh += myPow(10, 1-i) * EEPROM.read(3 + i);
    bright += myPow(10, 2-i) * EEPROM.read(7 + i);
    kinDelay += myPow(10, 2-i) * EEPROM.read(10 + i);
  }
}

//***************************************************
//**************  LOOP  ************************
//***************************************************
void loop(void) {
    Serial.print("Pre-heating plate/lid to "); Serial.print(ths[0].setTh,1);
    Serial.print("/"); Serial.print(ths[1].setTh,1); Serial.print(" from "); 
    Serial.print(Get_temp(PINS_TEMP[0]),1);  Serial.print("/"); Serial.println(Get_temp(PINS_TEMP[1]),1);
    if (EEPROM.read(6)==0) sprintf(odFile,"KINE_---.mrX");
    if (EEPROM.read(6)==1) sprintf(odFile,"SING_---.mrX");
    if (EEPROM.read(6)==2) sprintf(odFile,"CALI_---.mrX");
    if (EEPROM.read(6)==3) sprintf(odFile,"PMAP_---.mrX");
    odFile[11] = THIS_MINIREAD;
    *(odFile-1) = Build_FileName(odFile);
    btn=btnNONE;                // btnNONE;
    while (btn==btnNONE) {
      lcd.setCursor(2,3); lcd.print(odFile);
      lcd.setCursor(5,1);  lcd.write(" Idle ");
 //     btn = Cycle_heating (&ths[0], &ths[1], TAU_HEAT, TAU_MAX, CLOSE_TEMP_RANGE);
      btn = LED_LDR_pre_heating(bright, true);
      if (btn==btnGO) {
        if (EEPROM.read(6)==0) {   // GROWTH KINETICS
          Color_led(6,0.2);         
          lcd.setCursor(5,1);  lcd.write("Waiting");
          Increment_file_index(13);
          *(odFile-1) = Build_FileName(odFile);
          Serial.print("\nRecording kinetics to file: "); Serial.println(odFile);
          lcd.setCursor(2,3);  lcd.write(odFile);
          lcd.blink();
          lcd.setCursor(4,0);  lcd.write(" KINE  ");
          lcd.setCursor(0,1);  lcd.write(" Load plate...  ");
          lcd.setCursor(0,2);  lcd.write(" then press GO  "); lcd.setCursor(14,1);
          btn=btnNONE;
          while (btn==btnNONE) btn = Read_buttons();
          lcd.noBlink();
          if (btn==btnGO) btn=Growth_kinetics(bright, odFile, kinDelay, true);
        }
        if (EEPROM.read(6)==1) {   // SINGLE SCAN
          Color_led(4,0.2);
          lcd.setCursor(5,1);  lcd.write("Waiting");
          Increment_file_index(16);
           *(odFile-1) = Build_FileName(odFile);
          Serial.print("\nRecording kinetics to file: "); Serial.println(odFile);
          lcd.setCursor(2,3);  lcd.write(odFile);
          lcd.blink();
          lcd.setCursor(4,0);  lcd.write(" SINGLE"); lcd.setCursor(10,0);
          lcd.setCursor(0,1);  lcd.write(" Load plate...  ");
          lcd.setCursor(0,2);  lcd.write(" then press GO  "); lcd.setCursor(14,1);
          btn=btnNONE;
          while (btn==btnNONE) btn = Read_buttons();
          lcd.noBlink();
          lcd.clear();
          if (btn==btnGO) btn=Single_scan(bright, odFile, 0, 0, true);
        }
        if (EEPROM.read(6)==2) {   // CALIBRATION
          Color_led(1,0.2);
          Increment_file_index(19);
           *(odFile-1) = Build_FileName(odFile);
          Serial.print("\nStarting calibration to file: "); Serial.println(odFile);
          lcd.setCursor(2,3);  lcd.write(odFile);
          lcd.blink();
          lcd.setCursor(4,0);  lcd.write(" CALIB "); lcd.setCursor(10,0);
          lcd.clear();
          btn=Calibration(bright, odFile, false);
          lcd.noBlink();
        }
        if (EEPROM.read(6)==3) {   // PLATE MAPPING
          Color_led(4,0.2);
          lcd.setCursor(5,1);  lcd.write("Waiting");
          Increment_file_index(22);
           *(odFile-1) = Build_FileName(odFile);
          Serial.print("\nRecording plate map to file: "); Serial.println(odFile);
          lcd.setCursor(2,3);  lcd.write(odFile);
          lcd.blink();
          lcd.setCursor(4,0);  lcd.write(" PL MAP "); lcd.setCursor(10,0);
          lcd.setCursor(0,1);  lcd.write(" Load plate...  ");
          lcd.setCursor(0,2);  lcd.write(" then press GO  "); lcd.setCursor(14,1);
          btn=btnNONE;
          while (btn==btnNONE) btn = Read_buttons();
          lcd.noBlink();
          lcd.clear();
          if (btn==btnGO) btn=Plate_mapping(bright, odFile, true);
        }
        btn=btnNONE;
        lcd.clear();
      }
 
      if (btn==btnSET) {
        lcd.clear();
        lcd.blink();       
        set_params();
        lcd.noBlink();
        lcd.clear();
        ths[0].setTh=0;  ths[1].setTh=0, bright=0, kinDelay=0;
        for (uint8_t i=0; i<3; i++) {
          ths[0].setTh += myPow(10, 1-i) * EEPROM.read(0 + i);
          ths[1].setTh += myPow(10, 1-i) * EEPROM.read(3 + i);
          bright += myPow(10, 2-i) * EEPROM.read(7 + i);
          kinDelay += myPow(10, 2-i) * EEPROM.read(10 + i);
        }
      }   
        
      if (btn==btnCANCEL) {
        Color_led(5,0.2);
        btn = Read_external_buttons();
        int cnt=0;
        lcd.setCursor(0,0);  lcd.write("    RESETTING   ");
        lcd.setCursor(0,1);  lcd.write("FILE NUMs IN 20s ");
        while (btn==btnCANCEL) {
          btn = Read_external_buttons();
          delay(50);
          cnt++;
          if (cnt>400) {
            Color_led(1,0.2);    // button pressed for more than 400*50 ms = 20s
            lcd.setCursor(0,1);  lcd.write("FILE NUMs IN 30s ");
            lcd.setCursor(0,1);  lcd.write("    DONE !       ");
          }
        }
        if (cnt>400) {
          for (uint8_t ixCan=13; ixCan<13+3*NB_PROTOCOLS; ixCan++) EEPROM.put(ixCan,0);  // RESETS FILE INDICES
          lcd.setCursor(0,1);    lcd.write("       DONE!   ");    delay(1000);
        } else {
          lcd.setCursor(0,1);    lcd.write("    CANCELLED   ");
        }
      delay(1000);
      lcd.clear();
    }
  }
}


//=================================================
//===================== FUNCTIONS =================
//=================================================

//********************************************************
//***********  UI RELATED FUNCTIONS  ************
//********************************************************

//---------------------------------------------------
void Increment_file_index(uint8_t adr) {
  float value=0;
  for (int i=0; i<3; i++) value += myPow(10, 2-i) * EEPROM.read(adr + i);
  value ++;
  EEPROM.put(adr  ,((int)value % 1000)/100);
  EEPROM.put(adr+1,((int)value % 100)/10);
  EEPROM.put(adr+2,((int)value % 10)/1);
  return(value);
}

//---------------------------------------------------
void inc_eeprom(uint8_t adr) {
    uint8_t val=0;
    EEPROM.get(adr, val);
    val ++;
    if (val == 10) val = 0;
    if (adr==6 &  val == NB_PROTOCOLS) val = 0;
    EEPROM.put(adr, val);
}

//---------------------------------------------------
void dec_eeprom(uint8_t adr) {
    uint8_t val=0;
    EEPROM.get(adr, val);
    val --;
    if (val == 255) {
      val = 9;
      if (adr==6) val = NB_PROTOCOLS-1;
    }
    EEPROM.put(adr, val);
}

//---------------------------------------------------
void display_digit(uint8_t iDi) {     // Displays digit being set
  uint8_t digit=0, row=0, col=0;
  digit = EEPROM.read(iDi);
  if (iDi>12 & iDi<16) digit = EEPROM.read(iDi+3*EEPROM.read(6));
  lcd.setCursor(2, 0); lcd.print(".");
  lcd.setCursor(14, 0); lcd.print(".");
  
  if (iDi<6) {
    row=0;
    lcd.setCursor(6, row); lcd.print("SET");
    col = iDi + (iDi>1)+ (iDi>2)*8 + (iDi>4);
    lcd.setCursor(col, row); lcd.print(digit);
  }
  if (iDi==6) {
    row=1; col=0;
    if (digit==0) { lcd.setCursor(0, row); lcd.print("    KINETICS  "); }
    if (digit==1) { lcd.setCursor(0, row); lcd.print("  SINGLE SCAN "); }
    if (digit==2) { lcd.setCursor(0, row); lcd.print("  CALIBRATION "); }
    if (digit==3) { lcd.setCursor(0, row); lcd.print("   PLATE MAP  "); }
  }
  if (iDi>6 & iDi<10) {
    row=2;
    lcd.setCursor(0, row); lcd.print("Bri");
    col = 3 + iDi - 7;
    lcd.setCursor(col, row); lcd.print(digit);
    lcd.setCursor(7, row); lcd.print("-");
  }
  if (iDi>9 & iDi<13) {
    row=2;
    lcd.setCursor(12, row); lcd.print("min");
    col = 9 +      iDi - 10;
    lcd.setCursor(col, row); lcd.print(digit);
  }
  if (EEPROM.read(6)==0) { lcd.setCursor(0, 3); lcd.print("  KINE_"); }
  if (EEPROM.read(6)==1) { lcd.setCursor(0, 3); lcd.print("  SING_"); }
  if (EEPROM.read(6)==2) { lcd.setCursor(0, 3); lcd.print("  CALI_"); }
  if (EEPROM.read(6)==3) { lcd.setCursor(0, 3); lcd.print("  PMAP_"); }
  lcd.setCursor(10, 3); lcd.print(".mr");  lcd.print(THIS_MINIREAD); 
 // if (iDi>12 & iDi<16) {
  for (uint8_t jdi=13; jdi<16; jdi++) {
    lcd.setCursor(7 + jdi - 13, 3); lcd.print(EEPROM.read(jdi+3*EEPROM.read(6)));
  }
  if (iDi>12 & iDi<16) {
    row=3;
    col=7 + iDi - 13;
    lcd.setCursor(col, row); lcd.print(digit);
  }
 lcd.setCursor(col, row);
}

//---------------------------------------------------
void set_params() {     // enters settings mode
    uint8_t btn=0, iDi=0;
    bool settingsOk = false;
    while (Read_external_buttons() != btnNONE) {}
    for (uint8_t iDi2=0; iDi2<18; iDi2++) display_digit(iDi2);
    lcd.setCursor(0, 0);
    while ( ! settingsOk) {
        btn = Read_buttons();
        if (btn==btnDOWN) {
          if (iDi>12 & iDi<16) dec_eeprom(iDi+3*EEPROM.read(6)); else dec_eeprom(iDi);
        }
        if (btn==btnUP) {
          if (iDi>12 & iDi<16) inc_eeprom(iDi+3*EEPROM.read(6)); else inc_eeprom(iDi);       
        }
        if (btn==btnRIGHT) iDi ++;
        if (iDi == 16) iDi = 0;
        if (btn==btnLEFT) iDi --;
        if (iDi == 255) iDi = 15;
        if (btn==btnSET) settingsOk = true;
        display_digit(iDi);
    }
}

//---------------------------------------------------
float myPow(float x, int n) {     // Returns x^n for n signed integer
  float res = x;
  if (n == 0) return(1);
  int nAbs = n;
  if (n < 0) nAbs = -n;
  for (int i=0; i<nAbs-1; i++) res *= x;
  if (n < 0) return(1.0/res); else return(res);
}


//---------------   Internal function for Reads keys pressed  ----------------
uint8_t Read_external_buttons() {
  uint8_t p0 = digitalRead(PIN_BTleft);
  delay(2);
  uint8_t p1 = digitalRead(PIN_BTright);
  delay(2);
  uint8_t p2 = digitalRead(PIN_BTdown);
  delay(2);
  uint8_t p3 = digitalRead(PIN_BTup);
  delay(2);
  if ((p0==0) & (p1==0) & (p2==0) & (p3==0)) return btnNONE; // no button pressed
  if ((p0==1) & (p1==0) & (p2==0) & (p3==0)) return btnLEFT;
  if ((p0==0) & (p1==1) & (p2==0) & (p3==0)) return btnRIGHT;
  if ((p0==0) & (p1==0) & (p2==1) & (p3==0)) return btnDOWN;
  if ((p0==0) & (p1==0) & (p2==0) & (p3==1)) return btnUP;
  if ((p0==1) & (p1==1) & (p2==1) & (p3==0)) return btnSET;
  if ((p0==0) & (p1==1) & (p2==1) & (p3==1)) return btnGO;
  if ((p0==1) & (p1==1) & (p2==1) & (p3==1)) return btnCANCEL;
 return btnNONE;
}

//---------------   Reads keys pressed  ----------------
uint8_t Read_buttons() {
    uint8_t btn = btnNONE;
    while (Read_external_buttons()!=btnNONE) {}
    while (Read_external_buttons()==btnNONE) {}
    while (btn==btnNONE) {
        btn = Read_external_buttons();   // Reads external keys 
        while (Read_external_buttons()!=btnNONE) {}
    }
    return(btn);
}

//---------------   Displays float with LCD  ----------------
void DisplayFloat (float myFloat) {
    if (myFloat<0) { lcd.print("Neg "); return; }
    if (myFloat>99.9) { lcd.print("100+"); return; }

    myFloat += 0.05;
    if (myFloat < 10.0) lcd.print(" ");
    lcd.print(int(myFloat));
    lcd.print(".");
    lcd.print(int(myFloat*10)-int(myFloat)*10);
}

// -------------  Displays all temperature values on LCD   --------------   
void DisplayTemps(S_THER *myTh) {
  lcd.clear();  
  int col=0, row=0;
  for (int iTh=0; iTh<NB_THS; iTh++) {
    if (iTh == 0) { lcd.setCursor(0, 0); }
    if (iTh == 1) { lcd.setCursor(12, 0); }
    DisplayFloat(myTh[iTh].th);
  }
}

//********************************************************
//***************  RGB LED RELATED FUNCTIONS  ************
//********************************************************

//==============  Flash for error  ============
void Error_flash(uint8_t nbFlashes, uint8_t color, float bright) {
  uint8_t btn = btnNONE;
  Serial.println("SD card error !!!");
  lcd.setCursor(0,1); lcd.print("SD CARD ERROR !!");
  while (btn != btnCANCEL) {
    btn = Read_external_buttons();
    for (byte i=1; i<=nbFlashes; i++) { 
      Color_led(color, bright);        delay(100); 
      Color_led(0, 0);            delay(200); 
    }
    delay(1000);
  }
  lcd.clear();
}

//=========== Controls RGB LED =====================
void Color_led(uint8_t color, float bright) {
    analogWrite(PIN_LED1R, (uint8_t) 1.0*COLOR_R[color]*bright); 
    analogWrite(PIN_LED2G, (uint8_t)1.0*COLOR_G[color]*bright);
    analogWrite(PIN_LED3B, (uint8_t)1.0*COLOR_B[color]*bright); 
}

//=========== Breeze =====================
void breeze(void) {
  float iii=0.5;
  bool aaa=true;
  while(1) {
    Color_led(3,iii);
    if (aaa) iii+=0.001; else iii-=0.001;
    if (iii>0.999 | iii<0.001) aaa = !aaa;
    delay(2);
  }
}

//=========== Scan_colors =====================
void Scan_colors(void) {
  while (1) {
    for (int cc=0; cc<8;cc++) {
      Color_led(cc,1);
      delay(100);
    }
  }
}

//********************************************************
//***************  SD CARD RELATED FUNCTIONS  **************
//********************************************************
          
          //=========== SD_initialize =====================
          void Sd_test(Sd2Card sdCard) {
            SdVolume sdVolume;
            uint32_t sdVolumeSize;
            if (!sdCard.init(SPI_HALF_SPEED, PINS_SD[3])) {
              Serial.println(F("SD card initialization failed. Please check if card is correctly inserted."));
              Error_flash(3,1,100);  // 3 flashes => SD card initialization problem
            } else {
              switch (sdCard.type()) {
                case SD_CARD_TYPE_SD1:
                  Serial.print(F("SD1 card found ")); break;
                case SD_CARD_TYPE_SD2:
                  Serial.print(F("SD2 card found ")); break;
                case SD_CARD_TYPE_SDHC:
                  Serial.print(F("SDHC card found ")); break;
                default:
                  Serial.println(F("Unknown SD card type"));
              Error_flash(3,1,100);  // 3 flashes => SD card initialization problem
              }
            }
            if (!sdVolume.init(sdCard)) {
              Serial.println(F("No FAT16/FAT32 partition found. Is the card correctly formatted?"));
              Error_flash(3,1,100);  // 3 flashes => SD card initialization problem
            } else {
              Serial.print(F("FAT")); Serial.print(sdVolume.fatType(), DEC); Serial.print(F(", ")); 
            }
            sdVolumeSize = sdVolume.clusterCount() * sdVolume.blocksPerCluster() / 2; // blocks size in SD cards is always 512 bytes
            Serial.print(sdVolumeSize/1024/1024.0); Serial.print(F(" GB, "));
            Serial.print(sdVolume.clusterCount()); Serial.print(F(" clusters, ")); 
           // Serial.println(sdVolume.blocksPerCluster()); Serial.print(F("blocs per cluster, ")); 
            Serial.print(sdVolume.blocksPerCluster()*sdVolume.clusterCount()); Serial.println(F(" blocks")); 
          }
          
          //=========== sd_write =====================
          void Sd_write(char* odFile, char* line) {
            if (!SD.begin(PINS_SD[3])) {
              Serial.println("SD card initialization failed!");
              while (1);
            }
            dataFile = SD.open(odFile, FILE_WRITE);
            if (dataFile) {
              Serial.print("Writing to "); Serial.print(odFile);
              dataFile.println(line);
              dataFile.close();
              Serial.println(" done:");
              Serial.println(line);
            } else {
              Serial.println("ERROR opening file");
            }
          }
          
          //=========== sd_read =====================
          void Sd_read(char* odFile) {
            char line[256];
            if (!SD.begin(PINS_SD[3])) {
              Serial.println("SD card initialization failed!");
              while (1);
            }
            dataFile = SD.open(odFile);
            if (dataFile) {
              Serial.print("Reading from file "); Serial.println(odFile);
              while (dataFile.available()) {
                Serial.write(dataFile.read());
              }
              dataFile.close();
            } else {
              Serial.println("ERROR opening file");
            }
          }
          
          //=========== sd_remove_file_if_exists =====================
          void Sd_remove_file_if_exists(char* odFile) {
            if (!SD.begin(PINS_SD[3])) {
              Serial.println(F("SD card initialization failed. Please check if card is correctly inserted."));
              Error_flash(3,1,100);  // 3 flashes => SD card initialization problem
            } else {
              if (SD.exists(odFile)) {
                Serial.print(F("WARNING, file ")); Serial.print(odFile); Serial.println(F(" already exists, will be deleted."));
                if(!SD.remove(odFile)) {
                  Serial.println(F("ERROR File deletion failed..."));
                  Error_flash(3,1,100);
                } // else Serial.println(F("File successfully deleted."));
              } // else Serial.print(F("No file ")); Serial.print(odFile); Serial.println(F(" found in card"));
            }
          }

//=========== Build_FileName =====================
char Build_FileName(char *odFile) {
  char odIndexStr[4]="+++";
  float fileIndexFloat=0;
  int fileIndexByte=0;
  for (int i=0; i<3; i++) fileIndexFloat += myPow(10, 2-i) * EEPROM.read((13 + EEPROM.read(6)*3) + i);
  fileIndexByte = fileIndexFloat;
  sprintf(odIndexStr, "%03d", fileIndexByte);
  for (uint8_t i=0; i<3; i++) odFile[i+5]=odIndexStr[i];
//  if (USE_SD_CARD) Sd_remove_file_if_exists(odFile);
  Serial.print("Current file name: "); Serial.println(odFile);
  return(odFile);
}

//********************************************************
//***************  LIGHT RELATED FUNCTIONS  **************
//********************************************************

//============== test_LEDs ========================
void Test_LEDs(uint8_t brightness) {
  for (byte iLine=0; iLine<8; iLine++) {
    for (byte iCol=0; iCol<12; iCol++) {
      digitalWrite(PINS_LED[iLine],LOW);
      analogWrite(PINS_PWM[iCol], brightness);
      delay(300);
      digitalWrite(PINS_LED[iLine],HIGH);
      analogWrite(PINS_PWM[iCol], 0);
    }
  }
}

//============== Ambient_light ========================
float Ambient_light(void) {
  int iCapt;
  float average;
  analogReference(DEFAULT);
  average = 0;
  for (iCapt=0; iCapt< 100; iCapt++) analogRead(PIN_LDRext);
  for (iCapt=0; iCapt< NB_CAPTURES_LIGHT; iCapt++) {
    average += analogRead(PIN_LDRext);
 //   delay(10);
  }
  average /= 1.0*NB_CAPTURES_LIGHT;
  return (average);
}

//============== read_light ========================
float Read_light(uint8_t iLine, uint8_t iCol, uint8_t brightness) {
  int iCapt;
  uint8_t analogPin = PINS_LDR[iCol];
  float average;
  analogReference(INTERNAL1V1);
  digitalWrite(PINS_LED[iLine],LOW);
  analogWrite(PINS_PWM[iCol], brightness); 
  digitalWrite(PINS_MPX_ADR[0], (iLine & 1));
  digitalWrite(PINS_MPX_ADR[1], (iLine & 2)/2);
  digitalWrite(PINS_MPX_ADR[2], (iLine & 4)/4);
  delay(500);
  average = 0;
  for (iCapt=0; iCapt< 100; iCapt++) analogRead(analogPin);
  for (iCapt=0; iCapt< NB_CAPTURES_LIGHT; iCapt++) {
    average += analogRead(analogPin);
 //   delay(10);
  }
  average /= 1.0*NB_CAPTURES_LIGHT;
  digitalWrite(PINS_LED[iLine],HIGH);
  analogWrite(PINS_PWM[iCol], 0);
  return (average);
}

//********************************************************
//***************  TEMPERATURE RELATED FUNCTIONS  **************
//********************************************************

//============== Adjust_temp ========================
uint8_t Adjust_temp(float precision) {
  long int cnt=0;
  uint8_t btn=btnNONE;
  bool adjustMore=true;
  while (adjustMore) {
    Serial.print(Get_temp(PINS_TEMP[0]),1);  Serial.print("/"); Serial.print(Get_temp(PINS_TEMP[1]),1);  Serial.print("  ");
    btn=Cycle_heating (&ths[0], &ths[1], TAU_HEAT, TAU_MAX, CLOSE_TEMP_RANGE);
    if (btn==btnNONE) {
      cnt++;
      if (cnt % 10 == 0) { cnt=0; Serial.println(); }
//      if (ths[0].th-ths[0].setTh>=0 & ths[0].th-ths[0].setTh<precision & ths[1].th-ths[1].setTh>=0 & ths[1].th-ths[1].setTh<precision) {
//         Serial.println("\nTemperatures ok");
//        cnt=0;
//      } 
    } else adjustMore=false;
  }
  return(btn);
}

//============== get_temp ========================
float Get_temp(uint8_t pinNb) {
  uint8_t i;
  float average, steinhart;

  analogReference(DEFAULT);
  average = 0;
  for (int i=0; i< NB_CAPTURES_TH; i++) {
    average += analogRead(pinNb);
    delay(2);
  }
  average /= NB_CAPTURES_TH;
  average = 1023 / average - 1;
  average = SERIES_RESISTOR_TH / average;
  steinhart = average / THERMISTOR_NOMINAL;         // (R/Ro)
  steinhart = log(steinhart);                     // ln(R/Ro)
  steinhart /= B_COEFF_TH;                         // 1/B * ln(R/Ro)
  steinhart += 1.0 / (TEMPERATURE_NOMINAL + 273.15);  // + (1/To)
  steinhart = 1.0 / steinhart;                    // Invert
  steinhart -= 273.15;                            // convert to C
  return steinhart;
}

//---------------   Cycle heating  ----------------
float Cycle_heating (S_THER *thisTh0, S_THER *thisTh1, float tauH, float tauM, float closeRange) {
  float meanX0=0, meanY0=0, varX0=0, covXY0=0;
  float meanX1=0, meanY1=0, varX1=0, covXY1=0, diff, red, blue;
  bool quitNow=false;
  thisTh0->th = Get_temp(PINS_TEMP[0]);
  thisTh1->th = Get_temp(PINS_TEMP[1]);
  lcd.setCursor(0,0); lcd.print(thisTh0->th,1);
  lcd.setCursor(12,0); lcd.print(thisTh1->th,1);
  
  for (uint8_t iSt=1; iSt<NB_TEMP_STORED; iSt++) thisTh0->storedTh[iSt-1] = thisTh0->storedTh[iSt];
  thisTh0->storedTh[NB_TEMP_STORED-1] = thisTh0->th;
  for (uint8_t iSt=1; iSt<NB_TEMP_STORED; iSt++) thisTh1->storedTh[iSt-1] = thisTh1->storedTh[iSt];
  thisTh1->storedTh[NB_TEMP_STORED-1] = thisTh1->th;

  for (uint8_t i=1; i<=NB_TEMP_STORED; i++) { meanY0 += thisTh0->storedTh[i-1]; meanX0 += 1.0*i; }
  for (uint8_t i=1; i<=NB_TEMP_STORED; i++) { meanY1 += thisTh1->storedTh[i-1]; meanX1 += 1.0*i; }
  meanY0 /= 1.0*NB_TEMP_STORED;  meanX0 /= 1.0*NB_TEMP_STORED;
  meanY1 /= 1.0*NB_TEMP_STORED;  meanX1 /= 1.0*NB_TEMP_STORED;
  for (uint8_t i=1; i<=NB_TEMP_STORED; i++) {
    varX0 += (1.0*i - meanX0) * (1.0*i - meanX0);
    covXY0 += (1.0*i - meanX0) * (thisTh0->storedTh[i-1] - meanY0);
    varX1 += (1.0*i - meanX1) * (1.0*i - meanX1);
    covXY1 += (1.0*i - meanX1) * (thisTh1->storedTh[i-1] - meanY1);
  }
  thisTh0->slope = covXY0 / varX0;
  thisTh1->slope = covXY1 / varX1;
  thisTh0->intercept = meanY0 - thisTh0->slope * meanX0;
  thisTh1->intercept = meanY1 - thisTh1->slope * meanX1;
  
  thisTh0->tau = 1.0*tauM*exp(-(thisTh0->th-thisTh0->setTh)*(thisTh0->th-thisTh0->setTh) / (closeRange*closeRange));
  thisTh1->tau = 1.0*tauM*exp(-(thisTh1->th-thisTh1->setTh)*(thisTh1->th-thisTh1->setTh) / (closeRange*closeRange));
  thisTh0->heat = thisTh0->heat-1.0/tauH*(thisTh0->slope*thisTh0->tau+thisTh0->th-thisTh0->setTh);
  thisTh1->heat = thisTh1->heat-1.0/tauH*(thisTh1->slope*thisTh1->tau+thisTh1->th-thisTh1->setTh);
  if (thisTh0->heat < 0 | thisTh0->th > thisTh0->setTh + 1 ) thisTh0->heat = 0;
  if (thisTh1->heat < 0 | thisTh1->th > thisTh1->setTh + 1 ) thisTh1->heat = 0;
  if (thisTh0->heat > 1 | thisTh0->th < thisTh0->setTh - 1 ) thisTh0->heat = 1;
  if (thisTh1->heat > 1 | thisTh1->th < thisTh1->setTh - 1 ) thisTh1->heat = 1;
  
  long int cnt=0;
  if (thisTh0->heat>0 & USE_HEATING) digitalWrite(PINS_HEAT[0], HIGH);    //  heater turned ON
  if (thisTh1->heat>0 & USE_HEATING) digitalWrite(PINS_HEAT[1], HIGH);    //  heater turned ON
  cnt = 0;
  while (!quitNow & cnt < CYCLE_TIME) {
    if (cnt > thisTh0->heat * CYCLE_TIME) digitalWrite(PINS_HEAT[0], LOW);  // heater OFF if time elapsed
    if (cnt > thisTh1->heat * CYCLE_TIME) digitalWrite(PINS_HEAT[1], LOW);  // heater OFF if time elapsed
    cnt++;
    btn = Read_external_buttons();
    if (btn!=btnNONE) quitNow=true;
  
    lcd.setCursor(4,0); lcd.print("     ");
    if (fabs(thisTh0->th - thisTh0->setTh) < 0.1) {
      lcd.setCursor(4,0); lcd.print(" ok");
    }     
    lcd.setCursor(9,0); lcd.print("   ");
    if (fabs(thisTh1->th - thisTh1->setTh) < 0.1) {
      lcd.setCursor(9,0); lcd.print("ok");     
    }
    if (fabs(thisTh0->th - thisTh0->setTh) < 0.1 & fabs(thisTh1->th - thisTh1->setTh) <0.2) {
      Color_led(0,0); delay(10); Color_led(2,0.1); delay(50); 
    } else {
      Color_led(0,0); delay(10);
      diff=1.0*(thisTh0->th - thisTh0->setTh);
      red=1.0*((diff)*15)+10;  if (red<0) red=0;
      blue=1.0*((-diff)*15)+10;  if (blue<0) blue=0;   
      analogWrite(PIN_LED3B, blue*1.2);
      analogWrite(PIN_LED1R, red); 
      analogWrite(PIN_LED2G, 0); 
      delay(50); 
    }
  }
//  Serial.print(diff); Serial.print("  "); Serial.print(red); Serial.print("  "); Serial.println(blue);
  digitalWrite(PINS_HEAT[0], LOW);
  digitalWrite(PINS_HEAT[1], LOW);
  return(btn);
}

//---------------   Scan heating regulation parameters  ----------------
void Scan_heating_params (S_THER *myThs) {
  int cnt=0, fileIndex=0;
  char odFile[13]="Scan----.txt";
  char odIndexStr[5]="++++";

  uint8_t                 nbNTS = 1;
  uint8_t t_nbTempStored[nbNTS] = {4};
  uint8_t                  nbTH = 1;
  float   t_tauH[nbTH]          = {20};
  uint8_t                  nbTM = 1;
  float   t_tauM[nbTM]          = {5};
  uint8_t                  nbCR = 1;
  float   t_closeRange[nbCR]    = {1.0};
  uint8_t                 nbTAR = 4;
  float   t_targets[nbTAR]      = {30, 28, 31, 29};
  int                  nbCycles = 5;

  EEPROM.get(sizeof(uint8_t), fileIndex);
  fileIndex ++;
  EEPROM.put(sizeof(uint8_t), fileIndex);
  sprintf(odIndexStr, "%04d", fileIndex);
  for (uint8_t i=0; i<4; i++) odFile[i+4]=odIndexStr[i];
  Sd_remove_file_if_exists(odFile); Serial.println();

  Serial.print("SCANNING PARAMETERS in file: ");          Serial.println(odFile);
  Serial.print("Waiting for temperature to drop below "); Serial.println(t_targets[0] - 1.5);
  while (Get_temp(PINS_TEMP[1])>t_targets[0] - 1  ) {   // +9999 
    Serial.print(Get_temp(PINS_TEMP[1]),1);  Serial.print("  ");
    delay(10000);  cnt++;
    if (cnt % 10 == 0) { cnt=0; Serial.println(); }
  }
  Serial.println(); Serial.println();
  for (uint8_t iTauH=0; iTauH<nbTH; iTauH++) {
    for (uint8_t iTauM=0; iTauM<nbTM; iTauM++) {       
      for (uint8_t iCloseRange=0; iCloseRange<nbCR; iCloseRange++) {
        for (uint8_t iTar=0; iTar<nbTAR; iTar++) {
          myThs->setTh = t_targets[iTar];
          for (uint8_t i=0; i<NB_TEMP_STORED; i++) myThs->storedTh[i] = myThs->setTh;
          analogWrite(PIN_LED2G,255);
          dataFile = SD.open(odFile, FILE_WRITE);
          if (dataFile) Error_flash(2,1,100);
          dataFile.print(myThs->setTh);                    dataFile.print("  ");
          dataFile.print(t_tauH[iTauH]);                   dataFile.print("  ");
          dataFile.print(t_tauM[iTauM]);                   dataFile.print("  ");
          dataFile.print(t_closeRange[iCloseRange]);       dataFile.print("  ");
          dataFile.print(11111);                           dataFile.println();
          dataFile.close();
          analogWrite(PIN_LED2G,0);
          Serial.print(myThs->setTh);                    Serial.print("  ");
          Serial.print(t_tauH[iTauH]);                   Serial.print("  ");
          Serial.print(t_tauM[iTauM]);                   Serial.print("  ");
          Serial.print(t_closeRange[iCloseRange]);       Serial.print("  ");
          Serial.print(11111);                           Serial.println();
   
          for (uint8_t iCycle=0; iCycle<nbCycles; iCycle++) {
            
    //        Cycle_heating (myThs, t_tauH[iTauH], t_tauM[iTauM], t_closeRange[iCloseRange]);
            
            Serial.print(iCycle);          Serial.print(" ");
            Serial.print(myThs->th,4);     Serial.print(" ");
            Serial.print(myThs->tau, 1);   Serial.print(" ");
            Serial.print(myThs->slope, 3); Serial.print(" ");
            Serial.print(myThs->heat,3);   Serial.print("     ");
          analogWrite(PIN_LED2G,255);
            dataFile = SD.open(odFile, FILE_WRITE);
            if (dataFile) Error_flash(2,1,100);
            dataFile.print(iCycle);          dataFile.print(" ");
            dataFile.print(myThs->th,4);     dataFile.print(" ");
            dataFile.print(myThs->tau, 1);   dataFile.print(" ");
            dataFile.print(myThs->slope, 3); dataFile.print(" ");
            dataFile.print(myThs->heat,3);   dataFile.print("     ");
            dataFile.close();
            Serial.println();
            dataFile = SD.open(odFile, FILE_WRITE);
            if (dataFile) dataFile.println();
            dataFile.close();
          analogWrite(PIN_LED2G,0);
          }
        }
      }
    }
  }
  Serial.println("Parameters scan completed");
  Serial.println();
}


//---------   Reads one line of wells to warm-up LDRs and LEDs. Does not record readings  ---------
uint8_t LED_LDR_one_line_warmup(uint8_t line, uint8_t bright, bool heat) {
  uint8_t nbFail=0;
  
  lcd.setCursor(5,0);  lcd.write("WarmUp");
  for (iCol=0; iCol<12; iCol+=1){ //12
    Color_led(0,0);
    btn = Read_external_buttons();
    if (btn!=btnNONE) return(btn);
    reading = Read_light(iLine, iCol, bright);
    Color_led(4,0.2);         
    btn = Read_external_buttons();
    if (btn!=btnNONE) {
      lcd.setCursor(5,0); lcd.write("WarmUp");
      return(btn);
    }
  }
  return(btn);
}


//---------   Reads complete plate to pre-heat LDRs and LEDs. Does not record readings  ---------
uint8_t LED_LDR_pre_heating(uint8_t bright, bool heat) {
  uint8_t nbFail=0;
  
  for (iLine=0; iLine<8; iLine+=1){  //8
    btn = Read_external_buttons();
    if (btn!=btnNONE) return(btn);
    if (heat) btn = Cycle_heating (&ths[0], &ths[1], TAU_HEAT, TAU_MAX, CLOSE_TEMP_RANGE);
    lcd.setCursor(5,0);  lcd.write("WarmUp");
    btn = Read_external_buttons();
    if (btn!=btnNONE) return(btn);
    btn=LED_LDR_one_line_warmup(iLine, bright, heat);
    btn = Read_external_buttons();
    if (btn!=btnNONE) return(btn);
  }
  return(btn);
}


//---------------   Reads plate and sends to serial according to plate plan  ----------------
uint8_t Plate_mapping(uint8_t bright, char *odFile, bool heat) {
  uint8_t nbFail=0, btn=btnNONE;

  while ( USE_SD_CARD & (! dataFile) & nbFail < 5) {
    dataFile = SD.open(odFile, FILE_WRITE);
    nbFail++; delay(50);
  }
  if (nbFail>=5) Error_flash(3,1,100);
  ths[1].th = Get_temp(PINS_TEMP[1]);
  ths[0].th = Get_temp(PINS_TEMP[0]);
  lcd.setCursor(0,0); lcd.print(ths[0].th,1);
  lcd.setCursor(12,0); lcd.print(ths[1].th,1);
  lcd.setCursor(0,2); lcd.print("Bri "); 
  lcd.setCursor(4,2); lcd.print(bright); 
  lcd.setCursor(0,3); lcd.print("                "); 
  lcd.setCursor(2,3); lcd.print(odFile);
  Serial.println();
  Serial.print("Brightness: "); Serial.print(bright);
  Serial.print("   Bottom temp. "); Serial.print(ths[0].th); Serial.print("   Lid temp. "); Serial.print(ths[1].th); Serial.println(); Serial.println();
  if (USE_SD_CARD) {
    dataFile.print("Brightness "); dataFile.print(bright);
    dataFile.print("    Bottom temp. "); dataFile.print(ths[0].th); dataFile.print("    Lid temp. "); dataFile.print(ths[1].th); dataFile.println(); dataFile.println();
  }
  if (USE_SD_CARD) 
  for (iLine=0; iLine<8; iLine+=1){
    for (iCol=0; iCol<12; iCol+=1){
      Color_led(0,0);
      reading = Read_light(iLine, iCol, bright);
      lcd.setCursor(0,1); lcd.print("Well            ");     
      lcd.setCursor(5,1); lcd.print((char)(iLine+65));    
      lcd.setCursor(6,1); lcd.print("  "); lcd.setCursor(6,1); lcd.print(iCol+1);    
      lcd.setCursor(8,1); lcd.print(":");
      lcd.setCursor(10,1); lcd.print("     "); lcd.setCursor(10,1); lcd.print(reading,1);       
      analogWrite(PIN_LED3B, 1.0*(iCol+1)/12*(8-(iLine+1))/8*90 );
      analogWrite(PIN_LED1R, 1.0*(iCol+1)/12*(  (iLine+1))/8*90);
      if (USE_SD_CARD) dataFile.print(reading); dataFile.print(" ");
      Serial.print(reading); Serial.print("  ");
      btn = Read_external_buttons();
      if (btn==btnCANCEL) {
        if (USE_SD_CARD) dataFile.close();
        return;
      }
    }
    if (USE_SD_CARD) dataFile.println();
    Serial.println();
  }
  if (USE_SD_CARD) dataFile.close();
  return(btn);
}


//---------------   Reads plate and sends to serial in a singel line  ----------------
uint8_t Single_scan(uint8_t bright, char* odFile, float initTime, uint8_t nbPaper, bool heat) {
  float elapTime=0, minutesBeforeScan=0;
  uint8_t btn=btnNONE;

  elapTime = (1.0*millis()-1.0*initTime)/1000/3600;    // Elapsed time in hours
  ths[1].th = Get_temp(PINS_TEMP[1]);
  ths[0].th = Get_temp(PINS_TEMP[0]);
  lcd.setCursor(0,0); lcd.print(ths[0].th,1);
  lcd.setCursor(12,0); lcd.print(ths[1].th,1);
  lcd.setCursor(0,3); lcd.print("                "); 
  lcd.setCursor(2,3); lcd.print(odFile);
  dataFile = SD.open(odFile, FILE_WRITE);
  if (!dataFile) {
    Serial.println(); Serial.print("Error opening file "); Serial.print(odFile); Serial.println("in 'Single_scan'");
    Error_flash(3,1,100);
  }
  Serial.print(elapTime,5);     Serial.print(" "); // column 'hours'
  Serial.print(bright);         Serial.print(" "); // column 'bright'
  Serial.print(ths[0].th);      Serial.print(" "); // column 'thBottom1'
  Serial.print(ths[1].th);      Serial.print(" "); // column 'thLid1'
  Serial.print(THIS_MINIREAD);  Serial.print(" "); // column 'device'
  Serial.print(0);              Serial.print(" "); // column 'rep'
  Serial.print(nbPaper);        Serial.print(" "); // column 'calib'
  if (USE_SD_CARD) { 
    dataFile.print(elapTime,5);     dataFile.print(" "); // column 'hours'
    dataFile.print(bright);         dataFile.print(" "); // column 'bright'
    dataFile.print(ths[0].th);      dataFile.print(" "); // column 'thBottom1'
    dataFile.print(ths[1].th);      dataFile.print(" "); // column 'thLid1'
    dataFile.print(THIS_MINIREAD);  dataFile.print(" "); // column 'device'
    dataFile.print(0);              dataFile.print(" "); // column 'rep'
    dataFile.print(nbPaper);        dataFile.print(" "); // column 'calib'
  }
  for (iLine=0; iLine<8; iLine+=1){  //8
    if (heat) {
      btn = Cycle_heating (&ths[0], &ths[1], TAU_HEAT, TAU_MAX, CLOSE_TEMP_RANGE);
    } else delay(600);
    lcd.setCursor(5,0);  lcd.write("READ..");
    lcd.setCursor(0,1);  lcd.write("               ");
    for (iCol=0; iCol<12; iCol+=1){ //12
      Color_led(0,0);
      reading = Read_light(iLine, iCol, bright);
      lcd.setCursor(0,1); lcd.print("Well            ");     
      lcd.setCursor(5,1); lcd.print((char)(iLine+65));    
      lcd.setCursor(6,1); lcd.print("  "); lcd.setCursor(6,1); lcd.print(iCol+1);    
      lcd.setCursor(8,1); lcd.print(":");
      lcd.setCursor(10,1); lcd.print("     "); lcd.setCursor(10,1); lcd.print(reading,1);       
      analogWrite(PIN_LED3B, 1.0*(iCol+1)/12*(8-(iLine+1))/8*90 );
      analogWrite(PIN_LED1R, 1.0*(iCol+1)/12*(  (iLine+1))/8*90);
      Serial.print(reading);      Serial.print(" ");
      if (USE_SD_CARD) {
        dataFile.print(reading);  dataFile.print(" ");
      }
      btn = Read_external_buttons();
      if (btn==btnCANCEL) {
        if (USE_SD_CARD) dataFile.close();
        lcd.setCursor(0,1); lcd.print("                ");     
        return(btn);
      }
    }
  }
  ths[0].th = Get_temp(PINS_TEMP[0]);  
  ths[1].th = Get_temp(PINS_TEMP[1]);
  Serial.print(ths[0].th);      Serial.print(" ");
  Serial.print(ths[1].th);      Serial.println();
  if (USE_SD_CARD) {
    dataFile.print(ths[0].th);  dataFile.print(" "); 
    dataFile.print(ths[1].th);  dataFile.println();
    dataFile.close();
  }
  lcd.setCursor(0,1); lcd.print("                ");     
  return(btn);
}

//---------------   Growth_kinetics  ----------------
uint8_t Growth_kinetics(uint8_t bright, char* odFile, int minutesBetweenReads, bool heat) {
  float cycleTime=0, elapTime=0, initTime=0;
  uint8_t btn=btnNONE, iLine=0, ix=0;
  Serial.print("Brightness: "); Serial.print(bright);
  Serial.print("  Delay between scans:"); Serial.print(minutesBetweenReads); Serial.println(" min");
  lcd.setCursor(5,0); lcd.print("KINE  "); 
  lcd.setCursor(0,1); lcd.print("                "); 
  lcd.setCursor(0,2); lcd.print("                "); 
  lcd.setCursor(2,3); lcd.print(odFile);
  initTime=1.0*millis();
  while (1) {
    Color_led(2,0.5);
    cycleTime = millis();
    lcd.setCursor(9,2); lcd.print("Bri "); lcd.setCursor(13,2); lcd.print(bright);
    btn=Single_scan(bright, odFile, initTime, 0, heat);
    lcd.setCursor(0,1); lcd.print("                "); 
    if (btn==btnCANCEL) return(btn);
    iLine=0;
    while (((float)millis()-1.0*cycleTime)/(float)1000 < minutesBetweenReads*60) {
      lcd.setCursor(1,1); lcd.print(((float)millis()-1.0*cycleTime)/(float)60000 , 1); 
      lcd.setCursor(6,1); lcd.print("/"); 
      lcd.setCursor(8,1); lcd.print(minutesBetweenReads); 
      lcd.setCursor(11,1); lcd.print("min "); 
      if (USE_HEATING & heat) {
        btn = Cycle_heating (&ths[0], &ths[1], TAU_HEAT, TAU_MAX, CLOSE_TEMP_RANGE);
        if (btn==btnCANCEL) return(btn);
        btn = LED_LDR_one_line_warmup(iLine, bright, heat);
      } else delay(200);
      if (btn==btnCANCEL) return(btn);
      elapTime = (1.0*millis()-1.0*initTime)/1000/3600;    // Elapsed time in hours
      if (elapTime>=10 & elapTime<100) ix=1;
      if (elapTime>=100 & elapTime<1000) ix=2;
      if (elapTime>=1000 & elapTime<10000) ix=3;
      lcd.setCursor(0,2); lcd.print(elapTime); lcd.setCursor(4+ix,2); lcd.print(" h ");
      iLine ++;
      if (iLine>=7) iLine=0;
    }
  }
  return(btn);
}

//---------------   Calibration  ----------------
uint8_t Calibration(uint8_t bright, char* odFile, bool heat) {
  uint8_t btn=btnNONE;
  int sample=0;  // 1;
  Serial.print("Brightness: "); Serial.print(bright);
  Serial.println("Starting calibration"); 
  lcd.setCursor(0,0); lcd.print(" CALIBRATION "); 
  lcd.setCursor(0,2); lcd.print("Bri "); 
  lcd.setCursor(4,2); lcd.print(bright); 
  lcd.setCursor(7,2); lcd.print(" Samp "); 
  lcd.setCursor(2,3); lcd.print(odFile);
  iLine=0;
  while (1) {
    lcd.setCursor(13,2); lcd.print(sample); 
    lcd.setCursor(0,1); lcd.print("Press GO to scan"); 
    btn = btnNONE;
    while (btn != btnGO) {
      btn = Read_external_buttons();
      if (btn==btnCANCEL) return(btn);
      btn = Cycle_heating (&ths[0], &ths[1], TAU_HEAT, TAU_MAX, CLOSE_TEMP_RANGE);
      if (btn==btnCANCEL) return(btn);
      btn = LED_LDR_one_line_warmup(iLine, bright, heat);
      if (btn==btnCANCEL) return(btn);
      iLine ++;
      if (iLine>=7) iLine=0;
    }
    lcd.setCursor(0,1); lcd.print("                "); 
    Color_led(2,0.5);
    btn=Single_scan(bright, odFile, 0, sample, heat);
    if (btn==btnCANCEL) return(btn);
    sample ++;
  }
  lcd.setCursor(0,0); lcd.print("             "); 
  return(btn);
}
```
