## Supplementary material for "MiniRead: a simple and inexpensive do-it-yourself device for multiple analyses of micro-organism growth kinetics": R package: MiniRead.R

```
#***********************************************************************
# Copyright (c) 2023 Matthieu Falque <>.
#
# This file is part of MiniRead.
#
#  MiniRead is free software: you can redistribute it and/or modify
#  it under the terms of the GNU General Public License as published by
#  the Free Software Foundation, either version 3 of the License, or
#  (at your option) any later version.
#  MiniRead is distributed in the hope that it will be useful,
#  but WITHOUT ANY WARRANTY; without even the implied warranty of
#  MERCHANTABILITY or FITNESS FOR A PARTICULAR PURPOSE.  See the
#  GNU General Public License for more details.
#
#  You should have received a copy of the GNU General Public License
#  along with MiniRead. If not, see <http://www.gnu.org/licenses/>.
#
#***********************************************************************

#' @description The MiniRead R package contains functions to calibrate the 
#' MniRead optical plate reader / turbidimeter and automatically analyze 
#' growth kinetics data produced by this device.\cr
#' The version numbering (x.y.z) of the MiniRead hardware and software has 
#' the following meaning:
#' - x is the version number of the hardware. A change in x means that printed
#' circuit board and/or electronic components have been modified
#' - y is the major software version number. The firmware (C software embedded
#' in the device) and the R package used for data calibration and analysis 
#' MUST have the same major version number y to ensure that the files 
#' generated by the devices can be analyzed with the R package
#' - z is the minor software version number. Indicates minor bug fixes 
#' or improvements. Changes in values of z do not affect the compatibility 
#' between firmware and R package
"_PACKAGE"


#' @importFrom grDevices pdf dev.off rainbow
#' @importFrom graphics abline image legend par points text
#' @importFrom stats aggregate lm predict smooth.spline
#' @importFrom utils read.table write.table


#' @name GetCalibFilePath
#' @title Gives path to MiniRead calibration file
#' @description Returns path to file 'MiniRead_CALIBRATION.rds' containing the 
#' calibration curves obtained by \code{\link{Calibrate}()} for all sensors
#' of the MiniRead device(s).
#' @details The file 'MiniRead_CALIBRATION.rds' will be stored in the folder
#' 'MiniRead_Calibration' in your home directory. If this folder doesn't exist, 
#' it wil be created, and if it does not contain a file named 
#' 'MiniRead_CALIBRATION.rds', the example version of this file present in
#' the package will be copied in 'MiniRead_Calibration'. But in that case, 
#' you will have to remplace it by your own file once you've performed
#' proper calibration of your device(s).
#' @return Character string. Path to the file containing the calibration curves 
#' of the MniRead device(s).
GetCalibFilePath <- function() {

  userCalibFilePath <- normalizePath(file.path(Sys.getenv("HOME"),
                                              "MiniRead_Calibration",
                                              "MiniRead_CALIBRATION.rds"),
                                    winslash='/')
  if( !file.exists(userCalibFilePath) ){
    dn = dirname(userCalibFilePath)
    if( !dir.exists(dn) ) {
      success = dir.create(dn, recursive=TRUE, mode="0755")
      if( !success ){
        stop( paste("Could not create directory:", dn, sep='\n'), domain=NA )
      }
    }
    installedCalibFilePath = system.file("extdata/MiniRead_CALIBRATION.rds"
                                         , package = "MiniRead")
    cat(paste0("WARNING: the example file for calibration curves was copied to "
               , userCalibFilePath,"\n"))
    cat("These calibration curves are only to serve as an example.\n")
    cat("You must do the calibration on your own MiniRead machines\n")
    cat("to replace this example file by yours.\n")
    success = file.copy(installedCalibFilePath, userCalibFilePath)
    if( !success ){
      stop( paste("could not copy file:", installedCalibFilePath, "to ", dn
                  , sep='\n'), domain=NA )
    }
  }
  return(userCalibFilePath)
}


#' @name ExampleFile
#' @title Returns path to example files
#' @description Gives path to MiniRead example output files
#' @param type Character string. May be one of the following:
#' - "KINE_B" outpout file of the KINE program of the MiniRead device B
#' used to measure cell concentration at regular time intervals during growth
#' - "CALI_A" output file of the CALI program of the MiniRead device "A"
#' - "CALI_B" output file of the CALI program of the MiniRead device "B"
#' - "CALI_C" output file of the CALI program of the MiniRead device "C"
#' - "CALI_D" output file of the CALI program of the MiniRead device "D"
#' - "CALI_E" output file of the CALI program of the MiniRead device "E"
#' - NULL then the list of files contained in folder "extdata" of the package
#' is returned as a vector
#' @details The KINETICS program of the MiniRead records growth kinetics data
#' while the CALIBRATION program is used to calibrate the machine with samples
#' of known cell concentration or OD.
#' @return 
#' - Character string: name with complete path of different types of 
#' example MiniRead output files
#' - or vector of Character strings containing the list of files contained
#' in the folder "(/inst/)extdata"
#' @examples
#' ExampleFile("KINE_B")
#' @export
ExampleFile <- function(type=NULL) {
  if (is.null(type)) {
    return(system.file("extdata", package="MiniRead") |> list.files())
  } else if (type=="KINE_B") {
    return(system.file("extdata", "KINE_026.MRB", package = "MiniRead") )
  } else if (type=="CALI_A") {
    return(system.file("extdata", "CALI_053.MRA", package = "MiniRead") )
  } else if (type=="CALI_B") {
    return(system.file("extdata", "CALI_025.MRB", package = "MiniRead") )
  } else if (type=="CALI_C") {
    return(system.file("extdata", "CALI_024.MRC", package = "MiniRead") )
  } else if (type=="CALI_D") {
    return(system.file("extdata", "CALI_026.MRD", package = "MiniRead") )
  } else if (type=="CALI_E") {
    return(system.file("extdata", "CALI_042.MRE", package = "MiniRead") )
  } else if (type=="CALI_REPS") {
    return(system.file("extdata", "CALI_REPS.txt", package = "MiniRead") )
  } else if (type=="HOMOG") {
    return(system.file("extdata", "HOMOGENEITY.txt", package = "MiniRead") )
  } else {
    stop(" in ExampleFile(): argument 'type' must be 'KINE_B' or 'CALI_X'")
  }
}


#' @name ConvertVx12ToVx13
#' @title Converts data from MiniRead x.1.2 to x.1.3 format
#' @details Reads file, converts, and overwrites the file with updated data if
#' writeFile==TRUE
#' @param mrFileName Character string. Name (with path) of file containing the 
#' MiniRead data to be read by the function
#' @param verbose Logical. If TRUE, the function will display explanatory 
#' messages during its execution
#' @param writeFile Logical. If TRUE, the function will write its results in a 
#' file
#' @returns Data frame. 
#
ConvertVx12ToVx13 <- function(mrFileName, verbose=TRUE, writeFile=TRUE) {
  
  if (verbose) cat("Conversion from v0.1.2 to v0.1.3 format...  ")
  if( ! file.exists(mrFileName)) stop(paste0("in 'ReadMiniReadData()': File ",
                                         mrFileName," not found"))
  mrData <- read.table(mrFileName, fill=TRUE)
  if ( ! (is.data.frame(mrData) && dim(mrData)[2]>=103)) {
    stop(paste0("Invalid data file ",mrFileName,
                " (MiniReadVx.1.2-compatible tables have at least 103 columns)"))
  }
  colnames(mrData)[(1:5)] <- c("hours","bright","thBottom1","thLid1",
                               "calib")
  colnames(mrData)[(6:101)] <- paste0("W",1:96)
  colnames(mrData)[(102:103)] <- c("thBottom2","thLid2")
  
  toRem <- c()
  for (iLine in 1:dim(mrData)[1]) {
    if (sum(is.na(mrData[iLine,]))>0) toRem <- c(toRem, iLine)
  }
  if (length(toRem)>0) {
    mrData <- mrData[-toRem,]
    if (verbose) print(paste0("Incomplete line: ", toRem, 
                              " removed in file ", mrFileName))
  }
  len <- length(unlist(strsplit(mrFileName,"")))
  device <- substr(mrFileName,len,len)
  device[device=="B"] <- rep("A", sum(device=="B"))
  device[device=="C"] <- rep("B", sum(device=="C"))
  device[device=="D"] <- rep("C", sum(device=="D"))
  device[device=="E"] <- rep("D", sum(device=="E"))
  device[device=="F"] <- rep("E", sum(device=="F"))
  mrData <- cbind.data.frame(mrData[,1:4], 
                             device=rep(device, nrow(mrData)),
                             rep=rep(0, nrow(mrData)),
                             mrData[, 5:ncol(mrData)]
  )
  if (verbose) cat("done\n")
  if (writeFile) {
    WriteMiniReadFile(mrFileName=mrFileName, mrData=mrData)
  }
  return(mrData)
}


#' @name IsMiniReadData
#' @title Checks if a data frame is compatible with the MiniRead data format
#' @description Stops if not all mandatory columns of a MiniRead data file 
#' are present. Does allow additional columns.
#' @param mrData Data frame.
#' Must have AT LEAST the following 105 columns:  
#' - Col1: 'hours' (float, in hours) time after starting program,  
#' - Col2: 'bright' (integer between 1 and 255): brightness of the 96 white 
#' LED lamps, **CAUTION** MUST be the same value as that used for calibration
#' - Col3:. 'thBottom1' (float, in degrees Celsius): temperature of the bottom 
#' of the plate, measured just before reading the plate
#' - Col4: 'thLid1' (float, in degrees Celsius): temperature of the lid of the 
#' plate, measured just before reading the plate
#' - Col5: 'device' (one letter): label of the MiniRead machine. Is 
#' automatically filled by the MiniRead.
#' - Col6: 'rep' (integer): index of the replicate. Is not used by the 
#' MiniRead device, but can be useful for subsequent analyses with the 
#' MiniRead R package.
#' - Col7: 'calib' (integer): index of the calibration point (used only for 
#' the CALIBRATION program), for instance: index of the point of a dilution series
#' of cell suspensions, or numbers of optical filter sheets superimposed. 
#' This value is automatically incremented by the MiniRead device before each 
#' next sample of a calibration series is read.
#' - Cols8-103: 'W1' to 'W96': well (in the order: A1..A12, B1..B12,.., H1..H12)
#' - Col104: 'thBottom2': (float, in degrees Celsius): temperature of the bottom of 
#' the plate, measured just after reading the plate
#' - Col105: 'thLid2': (float, in degrees Celsius): temperature of the lid of the 
#' plate, measured just after reading the plate
#' @return Logical. TRUE if the argument data frame is compatible with the 
#' MinRead format, else FALSE
#' @examples
#' kineFile <- ExampleFile("KINE_B")
#' dat <- ReadMiniReadData(kineFile)
#' IsMiniReadData(mrData=dat)
#' @export
#
IsMiniReadData <- function(mrData) {
  ok <- FALSE
  if (is.data.frame(mrData)) {
    if (dim(mrData)[2]>=105) {
      if (dim(mrData)[1]>0) {
        if (sum(colnames(mrData)[(1:7)] != c("hours","bright","thBottom1","thLid1",
                                             "device","rep","calib"))==0) {
          if (sum(colnames(mrData)[(8:103)] != paste0("W",1:96))==0) {
            if (sum(colnames(mrData)[(104:105)] != c("thBottom2","thLid2"))==0){
              ok <- TRUE
            } else stop("Invalid MiniRead data frame: wrong column names after wells")
          } else stop("Invalid MiniRead data frame: wrong column names for wells")
        } else stop("Invalid MiniRead data frame: wrong column names before wells")
      } else stop("Invalid MiniRead data frame: empty data frame")
    } else stop("Invalid MiniRead data frame: number of columns must be >= 105")
  } else stop("Invalid MiniRead data frame: not a data frame")
  return(ok)
}


#' @name ReadMiniReadData
#' @title Reads MiniRead output data, either from a file or a data frame
#' @description Returns MiniRead data frame from either a file or a data frame.  
#' Checks the validity of the format
#' @param verbose Logical. If TRUE, the function will display explanatory 
#' messages during its execution
#' @param mrDataOrFile Character string or Data frame.
#' - If character string: name (with path) of output file from the MiniRead
#' - If data frame: must contain at least the columns of a MiniRead output 
#' file as described for the parameter mrData of \code{\link{IsMiniReadData}()}
#' @return Data frame. Same format as for the parameter mrData 
#' of \code{\link{IsMiniReadData}()}
#' @examples
#' kineFile <- ExampleFile("KINE_B")
#' kineDat <- ReadMiniReadData(kineFile)
#' head(kineDat, 3)
#' @export
#
ReadMiniReadData <- function(mrDataOrFile, verbose=TRUE) {
  if ( ! is.data.frame(mrDataOrFile) && ! is.character(mrDataOrFile)) {
    stop(" in ReadMiniReadData(), 'mrDataOrFile' must be either a file name or 
         a data frame")
  }
  if (is.data.frame(mrDataOrFile)) {
    if (IsMiniReadData(mrData=mrDataOrFile)) {
      if (verbose) {
        cat("Checking format of data frame 'mrDataOrFile'\n")
      }
      mrData <- mrDataOrFile
    } else stop(" in ReadMiniReadData(), 'mrDataOrFile' data frame is not valid. 
                It must be compatible with MiniRead data as checked in 
                IsMiniReadData()")
  } else if (is.character(mrDataOrFile)) {
    if (verbose) cat(paste0("\nReading MiniRead output file:\n "
                            ,mrDataOrFile,"\n"))
    if( ! file.exists(mrDataOrFile)) stop(paste0("in 'ReadMiniReadData()': File ",
                                           mrDataOrFile," not found"))
    mrData <- read.table(file=mrDataOrFile, fill=TRUE)
    if (dim(mrData)[2]==103) {
      mrData <- ConvertVx12ToVx13(mrFileName=mrDataOrFile, writeFile=FALSE)
    }
    colnames(mrData)[1:7] <- c("hours","bright","thBottom1","thLid1"
                              , "device","rep","calib")
    colnames(mrData)[8:103] <- paste0("W",1:96)
    colnames(mrData)[104:105] <- c("thBottom2","thLid2")
    if (! IsMiniReadData(mrData)) stop()
    toRem <- c()
    for (iLine in 1:dim(mrData)[1]) {
      if (sum(is.na(mrData[iLine,]))>0) toRem <- c(toRem, iLine)
    }
    if (length(toRem)>0) {
      mrData <- mrData[-toRem,]
      if (verbose) cat(paste0("\nIncomplete line: "
                   , toRem, " removed in file\n",mrDataOrFile,"\n"))
    }
  } else stop(" Argument 'mrDataOrFile is not a string or a data frame")
  return(mrData)
} 


#' @name WriteMiniReadFile
#' @title Writes text files from MiniRead data table
#' @description Writes files similar to those produced by the MiniRead device,
#' from a data frame
#' @inheritParams ReadMiniReadData
#' @inheritParams IsMiniReadData
#' @param mrFileName Character String. Name of the file to be written
#' @examples
#' kineFile <- ExampleFile("KINE_B")
#' kineDat <- ReadMiniReadData(kineFile)
#' WriteMiniReadFile(mrFileName="MiniRead_exported_file.MRB", mrData=kineDat)
#' @export
#
WriteMiniReadFile <- function(mrFileName, mrData, verbose=TRUE) {
  if ( ! is.character(mrFileName)) stop(paste0("in 'WriteMiniReadFile(), the 
                          argument 'mrFileName' must be a file name"))
  if (! IsMiniReadData(mrData)) stop(paste0("Invalid data file"))
  if (verbose) cat(paste0("\nWriting data to ",getwd(),"/",mrFileName,"\n"))
  write.table(mrData, col.names=F, row.names=F, quote=F, file=mrFileName)
}


#' @name CheckWellsList
#' @title Checks validity of well list entry
#' @description Checks if the format of the well list provided is compatible 
#' with the other functions of the MiniRead package
#' @param wells Vector of characters. Must have one of the following forms:
#' - "all" Then all wells of the plate will be considered
#' - a vector of strings of the form 'Wxx' where xx is in 1-96 and represents 
#' the number of the well, ordered as 1(A1)-12(A12), 13(B1)-24(B12),...
#' 84(H1)-96(H12)
#' - a vector of plate well coordinates of the form c('A1', 'B2',...)
#' @return Vector of well names of the form "W1".."W96" following the order
#' 1(A1)-12(A12), 13(B1)-24(B12),...84(H1)-96(H12)
#' @examples
#' wells <- c("A1", "B2", "H12")
#' CheckWellsList(wells=wells)
#' wells <- c("W1", "W2", "W3", "W96")
#' CheckWellsList(wells=wells)
#' wells <- "all"
#' CheckWellsList(wells=wells)
#' @export
#
CheckWellsList <- function(wells) {
  validLetters <- c("A","B","C","D","E","F","G","H")
  if (! is.character(wells)) stop(" in CheckWellsList(): 'wells' must be a 
                                  character string")
  if (length(wells)==1 && wells=="all") {
    wells <- paste0("W", 1:96)
    return(wells)
  }
  if (sum(substr(wells,1,1) != "W")==0) { # all are Wxx
    if (sum(! substr(wells,2,3) %in% 1:96)==0) {
      return(wells)
    }
  }
  if (sum(! substr(wells,1,1) %in% validLetters)==0) { # none is Wxx
    if (sum(! as.numeric(substr(wells,2,3)))==0) {
      if (sum(! as.numeric(substr(wells,2,3)) %in% 1:12)==0) {
        wells <- WellCoords2WellNames(wellCoords=wells)
        return(wells)
      }
    }
  }
  stop(" in CheckWellsList(): invalid entry for 'wells' parameter")
}


#' @name GetCalibrationCurves
#' @title Reads calibration curves produced by \code{\link{Calibrate}()}
#' @description Accepts either lists of \code{\link{smooth.spline}()} objects 
#' or the name of a file containing these lists. Checks the structure of the 
#' data in each case
#' @inheritParams ReadMiniReadData
#' @param calibrationCurves Character string or List
#' - If list: lists of \code{\link{smooth.spline}()} output objects modeling
#' the calibration curves, as described for the output value 
#' of \code{\link{Calibrate}()}
#' - If character string: name of a compressed file generated with 
#' \code{\link{saveRDS}()} containing the list detailed above, from which
#' the calibration curves will be loaded
#' - If calibrationCurves="auto" (default value): as above, but the file 
#' 'MiniRead_CALIBRATION.rds' will be automatically loaded from the folder 
#' 'MiniRead_Calibration' in your home directory
#' @return list of lists of \code{\link{smooth.spline}()} output objects as 
#' explained above
#' @examples
#' calibFile <- ExampleFile("CALI_B")
#' cellConcentrations <- c(cal0=0.00E+00, cal1=9.60E+01, cal2=1.96E+02, 
#'                         cal3=2.94E+02, cal4=3.77E+02)
#' calibCurves <- Calibrate(mrDataOrFile=calibFile,
#'                          calibValues=cellConcentrations,
#'                          pdfName=NULL)
#' calibCurves <- GetCalibrationCurves()
#' @export
#
GetCalibrationCurves <- function(calibrationCurves="auto", verbose=TRUE) {

  if ( ! is.list(calibrationCurves)) {
    if (is.character(calibrationCurves)) {
      if ((calibrationCurves=="auto")[1L]) {
        internalCalibFile <- GetCalibFilePath()
        if (verbose) {
          cat("\nReading calibration curves from folder:\n")
          cat(paste0(internalCalibFile,"\n"))
        }
        calibrationCurves <- readRDS(file=internalCalibFile)
      } else {
        if (verbose) {
          cat(paste0("\nReading calibration curves from file ",
                     getwd(), calibrationCurves,"\n"))
        }
        if ( ! file.exists(calibrationCurves)) {
          stop(paste0(" in GetCalibrationCurves(), file ",calibrationCurves,
                      "not found"))
        }
        calibrationCurves <- readRDS(file=calibrationCurves)
      }
    } else stop(" in GetCalibrationCurves(): invalid argument calibrationCurves")
  } else { # is a list
    nbDevs <- length(calibrationCurves)
    iDev <- 1
    for (iDev in 1:nbDevs) {
      if (length(calibrationCurves[[iDev]]) != 96) {
        stop(" in GetCalibrationCurves(), invalid calibrationCurves list. Must be be NA or a 
               list[[1-x]] of list[[1-96]] of smooth.spline() output objects")
      }
      iWell <- 4
      for (iWell in 1:96) {
        ca <- calibrationCurves[[iDev]][[iWell]]
        if (length(ca)==21) {
          if (attributes(calibrationCurves[[iDev]][[iWell]])$class!="smooth.spline") {
            stop(paste0(" in GetCalibrationCurves(), 
                      calibrationCurves[[",iDev,"]][[",iWell,"]] is not NA or a 
                      smooth.spline() output object"))
          }
        } else {
          if (length(ca) != 1 || ! is.na(ca)) {
            stop(paste0(" in AnalyzeGrowth(), 
                  calibrationCurves[[",iDev,"]][[",iWell,"]] is a scalar but not NA"))
          }
        }
      }
    }
  }
  if (verbose) {
    cat(paste0("Calibration date ",attributes(calibrationCurves)$calibDate, "\n"))
    cat(paste0("Calibration data type: ",attributes(calibrationCurves)$calibType, "\n"))
    cat(paste0("Calibration units: ",attributes(calibrationCurves)$calibUnit, "\n"))
    cat(paste0("File used for calibration: ",attributes(calibrationCurves)$calibFile, "\n"))
    cat(paste0("Devices used for calibration: "
               , paste(attributes(calibrationCurves)$calibDevices, collapse=" "), "\n"))
    if (length(attributes(calibrationCurves)$calibWells)!=96) {
      cat(paste0("List of wells calibrated: "
                 , WellNames2WellCoords(
                   attributes(calibrationCurves)$calibWells), "\n"))
    } else {
      cat(paste0("Number of wells calibrated: 96\n"))
    }
  }
  return(calibrationCurves)
}


#' @name RemoveSaturated
#' @title Removes saturated data from MiniRead data
#' @description For each value >1023, either that value is turned into NA or
#' the intire line is removed
#' @inheritParams IsMiniReadData
#' @inheritParams ReadMiniReadData
#' @inheritParams CheckWellsList
#' @param removeNextLines Logical.
#' - If TRUE, the first line containing saturated (>1023) values and all following
#' lines till the end of the data frame are removed from the data file 
#' (e.g. to use for calibration data)
#' - If FALSE, values >1023 are turned into NA but no line is removed
#' @return Data frame. Similar to the input one
#' @examples
#' kineFile <- ExampleFile("KINE_B")
#' kineDat <- ReadMiniReadData(kineFile)
#' nonSatData <- RemoveSaturated(mrData=kineDat, removeNextLines=TRUE)
#' @export
#
RemoveSaturated <- function(mrData, wells="all", removeNextLines=FALSE
                            , verbose=TRUE) {
  if (verbose) cat("Looking for saturated values")
  wells <- CheckWellsList(wells)
  if (sum(mrData[,wells] >= 1023)>0) {  # MiniRead dynamics is 0-1023
    coordsSaturatedWells <- which(mrData[,wells] >= 1023, arr.ind=TRUE)
    ixSaturatedWells <- which(mrData[,wells] >= 1023, arr.ind=FALSE)
    satRows <- coordsSaturatedWells[,"row"]
    satCols <- coordsSaturatedWells[,"col"]
    i <- 1
    for (i in 1:length(satRows)) {
      col <- (satCols[i]-1) %% 12 + 1
      line <- c("A","B","C","D","E","F","G","H")[floor((satCols[i]-1)/12) + 1]
      if (verbose) cat(paste0("\nWARNING, saturated read removed for read #"
                              ,satRows[i], " in well ", line,col))
    }
    lastNonSaturated <- min(satRows)-1
    if (removeNextLines) {
      mrData <- mrData[order(mrData$calib),]
      mrData <- mrData[1:(min(lastNonSaturated)),]
      if (verbose) cat("\nAll corresponding lines will be removed, as well 
                       as all next ones")
    } else {
      mrData[,wells][mrData[,wells]>=1023] <- NA
      if (verbose) cat("\nThese values have been turned into NA")
    }
  }
  return(mrData)
}


#' @name Calibrate
#' @title Computes calibration curves of light sensors
#' @description Analyzes calibration data produced using the CALI program of 
#' the MiniRead with samples of known OD or cell concentration values (these 
#' values must be put in the 'calib' column of the data table). 
#' Uses splines to compute a non-linear calibration function for each of 
#' the 96 wells.
#' @inheritParams ReadMiniReadData
#' @inheritParams GetCalibrationCurves
#' @inheritParams CheckWellsList
#' @param calibValues Vector of numerics. Real values of e.g. cell concentration
#' for each point of the series of calibration samples that are being used to
#' calibrate the MiniRead.
#' @param smoothing Numeric. Value of the 'spar' parameter to be passed to the 
#' function \code{\link{smooth.spline}()} for non-linear modeling of the 
#' calibration curves
#' @param graphose Logical. If TRUE graphs will be generated
#' @param graphose Logical. If TRUE graphs are produced as pdf files
#' @param graphTitle Character string. Title of the graph to be produced
#' @param pdfName Character string or NULL
#' - If Character string: name of the graph file to be generated
#' - If NULL, no pdf file will be written
#' @param calibType Character string. Type of value to be displayed in the 
#' graphs as calibration values (e.g. OD, or cell concentration)
#' @param calibUnit Character string. Unit of the calibration 
#' value (e.g. millions cells/mL)
#' @param calibrationStorage Character string. 
#' - If calibrationStorage = "auto", the calibration data will be written in 
#' the file 'MiniRead_CALIBRATION.rds' in the folder /MiniRead_Calibration 
#' of your home directory.   
#' This ensures that this calibration file will be automatically found by 
#' the function \code{\link{GetCalibrationCurves}()}.
#' - Else, name (with path and without extension) of the 
#' compressed data file (.rds file) containing the calibration curves. 
#' @param printWellName Logical. If TRUE, well numbers will be displayed on 
#' the graph
#' @return \code{list[[1-nb_devices]] of list[[1-96 wells]]} of 
#' \code{\link{smooth.spline}()} output objects, each of them reflecting 
#' the calibration curve of one well of one MiniRead device.  
#' These objects contain all parameters needed to model the non-linear relationship 
#' between the raw values produced by the MiniRead, and the cell concentration 
#' (or OD or whatever type of turbidity reference used for calibrating the 
#' MiniReads).  
#' The returned list will be given four additional attributes:
#' - calibDate: Character string. Date when  \code{\link{Calibrate}()} was run 
#' to produce the 'calibrationCurves' data
#' - calibType: Character string. Type of values used for calibration 
#' (e.g. OD or cell concentration)
#' - calibUnits: Character string. Unit of the calibration values (e.g. M cells/mL)
#' - calibFile: Character string. Name of the file used for calibration, if
#' calibration is done from a single file (only one device and no replicates)
#' - calibDevices: Vector of characters. Lists devices named by their 
#' one-letter label) for which calibration has been carried out.
#' - calibWells: Vector of character strings. Lists wells for which calibration 
#' has been carried out. This attribute is a vector of well identifiers as 
#' described in \code{\link{CheckWellsList}()}
#' @details This function produces a graph showing for each sensor of each 
#' device the calibration curve with its non-linear fit, and a graph 
#' showing the precision of the correction applied
#' @examples 
#' # First with only one device:
#' calibFile <- ExampleFile("CALI_B")
#' cellConcentrations <- c(cal0=0.00E+00, cal1=9.60E+01, cal2=1.96E+02, 
#'                         cal3=2.94E+02, cal4=3.77E+02)
#' calibCurves <- Calibrate(mrDataOrFile=calibFile,
#'                          calibValues=cellConcentrations,
#'                          pdfName=NULL)
#' attributes(calibCurves)
#'  
#' # Now with five different devices:
#' calibFiles <- c(ExampleFile("CALI_A"), ExampleFile("CALI_B"), 
#'                 ExampleFile("CALI_C"), ExampleFile("CALI_D"), 
#'                 ExampleFile("CALI_E")
#'                )
#' allCalis <- NULL
#' for (file in calibFiles) {
#'   thisCali <- ReadMiniReadData(file)
#'   allCalis <- rbind.data.frame(allCalis, thisCali)
#' }
#' cellConcentrations <- c(cal0=0.00E+00, cal1=9.60E+01, cal2=1.96E+02, 
#'                         cal3=2.94E+02, cal4=3.77E+02)
#' calibCurves <- Calibrate(mrDataOrFile=allCalis,
#'                         calibValues=cellConcentrations,
#'                         pdfName="Calibration_Multiple_Devices")
#' attributes(calibCurves)
#' @export
#
Calibrate <- function(mrDataOrFile, 
                      calibValues, 
                      wells="all", 
                      smoothing=0.5, 
                      graphose=TRUE,
                      verbose=TRUE, 
                      graphTitle="", 
                      pdfName="Calibration_MiniRead",
                      calibType="Cell concentration",
                      calibUnit="10^6 cells/mL",
                      calibrationStorage="auto",
                      printWellName=TRUE) {
  
  wells <- CheckWellsList(wells)
  if (is.character(mrDataOrFile)) {
    len <- length(unlist(strsplit(mrDataOrFile,"")))
    calibFile <- substr(mrDataOrFile, len-11,len)
  } else calibFile <- "NA"
  dat <- ReadMiniReadData(mrDataOrFile=mrDataOrFile)
  dat <- dat[order(as.numeric(dat$calib)),]
  dat <- RemoveSaturated(mrData=dat, removeNextLines=TRUE)
  r2cWellsDevs <- list()
  deviceLabs <- unique(dat$device)
  
  vals <- unique(dat$calib)
  if ( ! is.vector(calibValues) || length(vals) != length(calibValues)) {
    stop(paste0(" in Calibrate(): different numbers of values in the calibration data and in 'calibValues'"))
  }
  
  if (graphose) {
    if ( ! is.null(pdfName)) pdf(paste0(pdfName,".pdf"), 14, 7)
    par(mfrow=c(1,2))
  }
  iDev <- 1
  for (iDev in 1:length(deviceLabs)) {
    datDev <- dat[dat$device==deviceLabs[iDev],]  
    y <- calibValues[unique(datDev$calib)+1]
    r2cWells <- list()
    iw <- 1
    ix <- 1
    for (ix in 1:96) { # final list MUST have 96 elements
      if (paste0("W",ix) %in% wells) {
        wellCoord <- WellNames2WellCoords(wellNames=wells[iw])$coord
        x <- datDev[, wells[iw]]
        if (length(x) != length(y)) {
          stop(paste0(" in Calibrate(): there are ",length(x)," data but "
                      ,length(y)," points of 'calibration'"))
        }
        read2calibSpline <- smooth.spline(x, y, spar=smoothing)
        iw <- iw + 1
      } else {
        read2calibSpline <- NA
      }
      r2cWells <- c(r2cWells, list(read2calibSpline))
    }
    r2cWellsDevs <- c(r2cWellsDevs, list(r2cWells))
  }
  if (graphose) {
    iDev <- 1
    for (iDev in 1:length(deviceLabs)) {
      datDev <- dat[dat$device==deviceLabs[iDev],]  
      y <- calibValues[unique(datDev$calib)+1]
      main <- paste0(graphTitle," MiniRead ",deviceLabs[iDev])
      ## first graph
      yLab <- paste0("Calibration value: ",calibType)
      if (calibUnit!="") yLab <- paste0(yLab," (",calibUnit,")")
      plot(NULL, ylim=range(y), xlim=range(datDev[, wells])
           , main=main, xlab="Raw MiniRead output", ylab=yLab)
      ix <- 1
      for (ix in 1:length(wells)) {
        thisWell <- wells[ix]
        iWell <- as.numeric(substr(thisWell,2,3))
        wellCoord <- WellNames2WellCoords(wellNames=thisWell)$coord
        x <- datDev[, thisWell]
        points(x, y, cex=0.5)
        x <- range(x)[1]:(range(x)[2]*1.3)
        points(x, predict(r2cWellsDevs[[iDev]][[iWell]], x=x)$y, type="l", col=2)
        if (printWellName) text(max(x)*0.8, max(y)-0.5, adj=2, wellCoord)
      }
      x <- calibValues[unique(datDev$calib)+1]
      ## second graph
      yLab <- calibType
      if (calibUnit!="") yLab <- paste0(yLab," (",calibUnit,")")
      plot(NULL, ylim=range(y), xlim=range(x), main=main
           , xlab=paste0("Real: ",yLab)
           , ylab=paste0("Predicted: ",yLab))
      ix <- 1
      for (ix in 1:length(wells)) {
        iWell <- as.numeric(substr(wells[ix],2,3))
        r <- datDev[, paste0("W",iWell)]
        y <- predict(r2cWellsDevs[[iDev]][[iWell]], x=r)$y
        points(x, y, type='p', col=1)
        points(y, y, type='l', col=2)
      }
    }
    if ( ! is.null(pdfName)) {
      dev.off()
      if (verbose) cat(paste0("Graphics written in file:\n",
                              getwd(),"/",pdfName,".pdf"))
    }
  }
  calibSplines <- r2cWellsDevs
  myDate <- format(Sys.time(), "%Y-%m-%d_%Hh%Mm%S")
  if ( calibrationStorage!="auto" && file.exists(calibrationStorage)) {
    cat(paste0("\n WARNING previous file ",calibrationStorage
               ," was overwritten !\n"))
  }
  attr(x=calibSplines, "calibDate") <- myDate
  attr(x=calibSplines, "calibUnit") <- calibUnit
  attr(x=calibSplines, "calibType") <- calibType
  attr(x=calibSplines, "calibFile") <- calibFile
  attr(x=calibSplines, "calibDevices") <- deviceLabs
  attr(x=calibSplines, "calibWells") <- wells
  if ( calibrationStorage!="auto") {
    saveRDS(calibSplines, file=calibrationStorage)
  }
  if (calibrationStorage == "auto") {
    internalCalibFile <- GetCalibFilePath()
    saveRDS(calibSplines, file=internalCalibFile)
    cat(paste0("\nCalibration curves saved successfully in:\n"))
   cat(paste0(internalCalibFile, "\n"))
  }
  calibSplines <- GetCalibrationCurves(calibrationCurves=calibSplines
                                         , verbose=verbose)
  if (verbose) {
    cat("\nCalibration completed\n")
  }
  par(mfrow=c(1,1))
  return(calibSplines)
}


#' @name ApplyCalibration
#' @title Applies calibration curve to raw values produced by the MiniRead
#' @description Uses sensor calibration curves previously obtained using
#' \code{\link{Calibrate}()} to transform MiniRead raw data provided as
#' argument 'mrDataOrFile' into optical density or cell concentration value
#' @inheritParams Calibrate
#' @inheritParams GetCalibrationCurves
#' @param removeSat Logical. If TRUE, saturated values will be replaced by NAs
#' @return Data frame with the same columns as the entry data (as returned 
#' by \code{\link{ReadMiniReadData}()} ) but with the raw turbidity 
#' values replaced by the corrected values after applying calibration curves.
#' The unit of the new values are the same as that of the calibration values.
#' The returned data frame will be given the four additional attributes that
#' are described for the returned output value of \code{\link{Calibrate}()}
#' @details Reads the attributes attached to the calibrationCurves list when it
#' was generated, and attach them to the outputted data frame. See description
#' of these attributes as for the returned value of \code{\link{Calibrate}()}. 
#' @examples
#' kineFile <- ExampleFile("KINE_B")
#' kineDat <- ReadMiniReadData(kineFile)
#' calibFile <- ExampleFile("CALI_B")
#' cellConcentrations <- c(cal0=0.00E+00, cal1=9.60E+01, cal2=1.96E+02, 
#'                         cal3=2.94E+02, cal4=3.77E+02)
#' calibCurves <- Calibrate(mrDataOrFile=calibFile,
#'                          calibValues=cellConcentrations,
#'                          pdfName=NULL)
#' calibratedData <- ApplyCalibration(mrDataOrFile=kineFile)
#' @export
#
ApplyCalibration <- function(mrDataOrFile
                             , calibrationCurves="auto"
                             , removeSat=FALSE
                             , verbose=TRUE) {
  kineData <- mrDataOrFile
  dat <- ReadMiniReadData(mrDataOrFile=kineData)
  calib <- GetCalibrationCurves(calibrationCurves=calibrationCurves)
  if (verbose) cat("\nApplying calibration to KINETICS file\n")
  if (removeSat) dat <- RemoveSaturated(mrData=dat, removeNextLines=FALSE) 
  newDat <- dat
  devsDat <- unique(dat$device)
  devsCal <- attributes(calib)$calibDevices
  if (sum( ! devsDat %in% devsCal)>0) {
    print(paste0("Device used for the data: ",devsDat))
    print(paste0("Devices present in the calibration data: ",devsCal))
    stop(" in ApplyCalibration(): no calibration data available for the 
         device used to produce the data to analyze")
  }
  wells <- paste0("W",1:96)
  nbDevs <- length(calib)
  iDev <- 1
  for (iDev in 1:nbDevs) {
    iWell <- 1
    for (iWell in 1:length(wells)) {
      reads <- dat[, wells[iWell]]
      readsWoNa <- reads[!is.na(reads)]
      spl <- calib[[iDev]][[iWell]]
      if (length(spl)==21) {
        predWoNa <- predict(object=spl, x=readsWoNa)$y
      } else {
        predWoNa <- NA
      }
      nas <- which(is.na(reads))
      res <- c()
      shift <- 0
      for (i in 1:length(reads)) {
        if ( ! i %in% nas) res <- c(res, predWoNa[i-shift])
        else {
          res <- c(res, NA)
          shift <- shift + 1
        }
      }
      newDat[,wells[iWell]] <- res
    }
  }
  attr(newDat, "calibDate") <- attributes(calib)$calibDate
  attr(newDat, "calibType") <- attributes(calib)$calibType
  attr(newDat, "calibUnit") <- attributes(calib)$calibUnit
  attr(newDat, "calibDevice") <- devsDat
  attr(newDat, "calibWells") <- attributes(calib)$calibWells
  attr(newDat, "calibFile") <- attributes(calib)$calibFile
  return(newDat)
}


#' @name AnalyzeGrowth
#' @title Complete growth analysis from raw MiniRead output
#' @description First applies calibration curve (optional), then analyzes growth 
#' kinetics to estimate maximum rate, lag time, and plateau value for each well
#' @inheritParams Calibrate
#' @inheritParams GetCalibrationCurves
#' @inheritParams CheckWellsList 
#' @param useCalibration Logical. If TRUE, will apply calibration curves to the 
#' raw MiniRead data to convert them into e.g. cell concentrations. If FALSE,
#' the raw turbidity values produced by the MiniRead will be directly used. 
#' Be careful, those raw values are not linearly related with cell
#' concentration, that is why calibration is necessary.
#' @param yUnit Character string. Text to be used in graphs to describe the
#' unit of the turbidity values (e.g. 'Millions Cells/mL')
#' @param yValue Character string. Text to be used in graphs to describe the 
#' type of turbidity values (e.g. 'cell concentration')
#' @param thrLag Numeric. Threshold used to determine the lag time. 
#' The lag time will be set as the time until turbidity reaches the
#' fraction 'thrLag' of its value at the plateau
#' @param maxTime Numeric. Maximum kinetic time (in hours) after which all data
#' will be ignored. If maxTime==NULL all data will be considered
#' @param flatPlateau Logical. If TRUE, the rate of evaporation of culture 
#' medium in the wells will be estimated from the plateau phase, and used to
#' correct the values to estimate kinetic parameters
#' @param yRange Vector (size 2) of numeric. Range of values to be used for the Y
#' scale in the graphs. If yRange==NULL, the range of the data will be used
#' @param mainTitle Character string. Text to be displayed as main title of 
#' the graphs
#' @param grOpt List. Contains the following graphical parameters:
#' - grOpt$text: Logical. If TRUE, displays text with parameter values in the graphs
#' - grOpt$deriv: Logical. If TRUE, plots the derivatives of the curves
#' - grOpt$lines: Logical. If TRUE, draws horizontal or vertical lines to show the 
#' values of estimated growth parameters
#' @param pdfName Character string or NULL. Name of the pdf file to be 
#' generated, containing the graphs
#' - if NULL, the graphs will be sent to the current graphic device
#' - if "auto", The pdf file name will be "Growth_" followed by the inputted
#' MiniRead output file name (if any), or "Growth_kinetics_analysis.pdf" if
#' input data in 'mrDataOrFile' is a data frame
#' @param writose Logical. If TRUE, writes the output data frame into a 
#' tabulated text file
#' @details The values displayed on the graphs and used to compute growth 
#' parameters can be for instance: 
#' - raw MiniRead output (when using no calibration curves; arbitrary turbidity
#' values not linearly correlated with OD), or
#' - OD values (when using calibration curves obtained with e.g. serial 
#' dilutions  of a colored chemical, or different numbers of sheets of 
#' optical filter paper superimposed), or
#' - cell concentrations (when using calibration curves obtained with serial 
#' dilutions of cell suspensions)
#' @return Data frame. Contains one line for each well and the following 
#' columns:
#' - 'device': label of the MiniRead machine ("A", "B",...), as automatically 
#' put by the MiniRead firmware in the column 'device' of its output files
#' - 'line': row coordinate on the 96 well plate, (from A to H)
#' - 'col': column coordinate on the 96-well plate, (from 1 to 12)
#' - 'maxRateRaw': maximum growth rate computed without correction for 
#' evaporation, (estimated from the maximum slope of the growth phase)
#' - 'lagRaw': lag time in hours computed without correction for evaporation. 
#' Lag time is defined as the time for the cell concentration to reach the
#' fraction 'thrLag' of the plateau value (default=5%)
#' - 'plateauRaw': cell concentration at the plateau,  computed without 
#' correction for evaporation
#' - 'snr': signal/noise ratio computed as the ratio between the range of cell
#' concentrations during the whole kinetics and the average of absolute values
#' of all difference between raw and smoothed values
#' - 'drift': rate of linear increase of concentration during the plateau 
#' phase due to the evaporation of the liquid medium
#' - 'maxRateCorr': like 'maxRateRaw', but after correcting for evaporation
#' - 'lagCorr': like 'lagRaw', but after correcting for evaporation
#' - 'plateauCorr': like 'plateauRaw', but after correcting for evaporation
#' @examples
#' calibFile <- ExampleFile("CALI_B")
#' cellConcentrations <- c(cal0=0.00E+00, cal1=9.60E+01, cal2=1.96E+02, 
#'                         cal3=2.94E+02, cal4=3.77E+02)
#' calibCurves <- Calibrate(mrDataOrFile=calibFile,
#'                          calibValues=cellConcentrations,
#'                          graphose=FALSE,
#'                          pdfName=NULL)
#' kineFile <- ExampleFile("KINE_B")
#' result <- AnalyzeGrowth(mrDataOrFile=kineFile,
#'                         wells="A1",
#'                         pdfName=NULL)
#' @export
#
AnalyzeGrowth <- function(  mrDataOrFile
                          , calibrationCurves="auto"
                          , useCalibration=TRUE
                          , yUnit="auto"
                          , yValue="auto"
                          , thrLag=0.05, maxTime=NULL
                          , flatPlateau=TRUE, yRange=NULL
                          , graphose=TRUE
                          , verbose=TRUE
                          , writose=TRUE
                          , pdfName="auto"
                          , wells="all"
                          , mainTitle="Growth kinetics"
                          , grOpt=list(text=T, deriv=F, lines=T)  ) {
  
  kineData <- ReadMiniReadData(mrDataOrFile=mrDataOrFile)
  if (!is.null(pdfName) && pdfName=="auto") {
    if (is.character(mrDataOrFile)) {
      len <- length(unlist(strsplit(mrDataOrFile,"")))
      if (substr(mrDataOrFile,len-3,len-1)%in%c(".mr",".MR")) {
        name <- substr(mrDataOrFile, len-11, len-4)
        name <- paste0("_",name,"_",substr(mrDataOrFile, len-2, len))
      } else name <- ""
    } else name <- ""
    pdfName <- resName <- paste0("Growth_kinetics_analysis", name)
  }
  if (!is.null(pdfName)) {
    len2 <- length(unlist(strsplit(pdfName,"")))
    if (substr(pdfName,len2-3,len2)%in%c(".pdf",".PDF")) {
      pdfName <- resName <- substr(pdfName,1,len2-4)
    } else resName <- pdfName
  } else resName <- "Growth_kinetics_analysis"
  wells <- CheckWellsList(wells)
  if ( ! is.logical(useCalibration)) {
    stop(" in AnalyzeGrowth: argument 'useCalibration' must be TRUE/FALSE")
  }
  if (useCalibration) {
    dat <- ApplyCalibration(mrDataOrFile=kineData, calibrationCurves=calibrationCurves)
    if (yValue=="auto") yValue <- attributes(dat)$calibType
    if (yUnit=="auto") yUnit <- attributes(dat)$calibUnit
    calibWells <- attributes(dat)$calibWells
  } else {
    dat <- ReadMiniReadData(mrDataOrFile=kineData, verbose=TRUE)
    if (yValue=="auto") yValue <- "MiniRead raw output values"
    if (yUnit=="auto") yUnit <- "arbitrary units"
    calibWells <- "all"
  }
  calibWells <- CheckWellsList(calibWells)
  wells <- intersect(wells, calibWells)
  if (length(unique(dat$device))>1) {
    cat("\n WARNING: the data contain several MiniRead labels")
  }
  if (graphose) G <- TRUE else G <- FALSE
  if ( ! is.null(yRange) && length(yRange)!=2) {
    stop("ERROR: grOpt$yRange must be either NULL or a vector of two 
         numeric values")
  }
  if (verbose) cat("Preparing plots and analyzing kinetics parameters\n")
	if (is.null(maxTime)) maxTime <- max(dat$hours)
	dat <- dat[dat$hours<=maxTime,]
	iMrx <- unlist(gregexpr(".MR", kineData))-1
	if (G && !is.null(pdfName)) {
	  pdf(file=paste0(pdfName, ".pdf"), 8, 8)
	  if (verbose) cat(paste0("\nGraphs saved as file ",paste0(pdfName, ".pdf")))
	}
	allRes <- NULL
	if (flatPlateau) {
		res <- data.frame(device=NA, line=NA, col=NA, maxRateRaw=NA, 
		                  lagRaw=NA, plateauRaw=NA, snr=NA,
		                  drift=NA, maxRateCorr=NA, lagCorr=NA, 
		                  plateauCorr=NA)
	} else {
		res <- data.frame(device=NA, line=NA, col=NA, maxRate=NA, lag=NA, 
		                  plateau=NA, snr=NA)
	}
	x <- dat[,"hours"]
	xLim <- c(0,max(x))
	iw <- 1
	ix <- 1
	for (ix in 1:96) { # final list MUST have 96 elements
	  if (ix %in% as.numeric(substr(wells,2,3))) {
	    thisWell <- wells[iw]
	    iWell <- as.numeric(substr(thisWell,2,3))
	    iw <- iw + 1
  	  well <- WellNames2WellCoords(wellNames=thisWell)
  		y1 <- dat[, thisWell]
  		y1 <- y1 - min(y1)
  		yLim1 <- c(0, max(y1))*1.1  #  c(0, max(dat[,wells]))
  		if ( ! is.null(yRange)) yLim1 <- yRange
  		main <- paste0(mainTitle, "  well ",well$line, well$col)
  		xLab <- "Time (hours)"
  		yLab <- paste0(yValue, " (",yUnit,")")
  		if (G) plot(x, y1, xlim=xLim, ylim=yLim1, main=main, xlab=xLab, ylab=yLab, 
  		     cex=0.8, col=1, lwd=1)
  		if (G) abline(h=0, lty=2, lwd=1) # bottom base line
  		spl1 <- smooth.spline(x=x, y=y1, spar=0.5)
  		yPred1 <- predict(spl1, x=x, deriv=0)$y
  		if (G) points(x, yPred1, type="l", col=8, lwd=2)  # grey solid curve
  		der1_1 <- predict(spl1, x=x, deriv=1)$y
  		der2_1 <- predict(spl1, x=x, deriv=2)$y
  		if (diff(range(y1))==0) next  #  If all data saturated
  		snr1 <- diff(range(y1)) / mean(abs(y1-yPred1))  # signal/noise ratio
  		iMaxRate1 <- which.max(der1_1)
  		if (G && grOpt$lines && ! flatPlateau) {
  		  abline(v=x[iMaxRate1], lty=3, col=2) # red thin dashed line
  		}
  		maxRate1 <- der1_1[iMaxRate1]
  		iBeforePlateau <- which(abs(der1_1)< 0.2 *max(abs(der1_1)) & 
  		                            x>x[iMaxRate1])
  
  		if (length(iBeforePlateau)>0) {
  			begPlat1 <- min(iBeforePlateau)+0.1*(length(x) - min(iBeforePlateau))
  			if (G && grOpt$lines) {
  			  abline(v=x[begPlat1], lty=3, col=4, lwd=1.5) # blue dashed: begin plateau
  			}
  			plateau1 <- y1[begPlat1]
  			iAfterPlateau <- which(abs(der1_1)>0.2*max(abs(der1_1)) & 
  			                           x>x[begPlat1])
  			if (length(iAfterPlateau)>0) {
  				endPlat1 <- max(iAfterPlateau)-0.2*(max(iAfterPlateau)-
  				                                        begPlat1)
  			} else endPlat1 <- length(x)
  			if (G && grOpt$lines) {
  			  abline(v=x[endPlat1], lty=3, col=4, lwd=1.5) # blue dashed: end plateau
  			}
  
  			if (length(which(der1_1>0 & y1>plateau1 * thrLag))>0) {
  				iLag1 <- min(which(der1_1>0 & y1>plateau1 * thrLag ))
  				lag1 <- x[iLag1]
  			} else {
  			  lag1 <- NA
  			  iLag1 <- -1
  			}
  			if (length(begPlat1:endPlat1)>1) {
  				regPlat1 <- lm(y1[begPlat1:endPlat1] ~ x[begPlat1:endPlat1])
  				platSlope1 <- regPlat1$coefficients[2]
  				platLine1 <- (x-x[begPlat1])*platSlope1+y1[begPlat1]
  				if (G && grOpt$lines) { # straight black dashed plateau line
  				  points(x[begPlat1:endPlat1], 
  				                        platLine1[begPlat1:endPlat1], 
  				                        type='l', lty=2, lwd=2)
  				}
  				y2 <- y1 - x * platSlope1
  			} else y2 <- y1
  
  			if (flatPlateau) {
  				spl2 <- smooth.spline(x=x, y=y2, spar=0.4)
  				yPred2 <- predict(spl2, x=x, deriv=0)$y
  				if (G) points(x, yPred2, type="l", col=6, lwd=2) # pink curve
  				der1_2 <- predict(spl2, x=x, deriv=1)$y
  				der2_2 <- predict(spl2, x=x, deriv=2)$y
  				if (G && grOpt$deriv) {
  				    points(x, der1_2+max(y1)*0.4, type="l", col=2) # thin red curve
  				    abline(h=max(y1)*0.4)
  				    points(x, der2_2+max(y1)*0.4, type="l", col=3) # thin green curve
  				    abline(h=max(y1)*0.4)
  				}
  				iMaxRate2 <- which.max(der1_2)
  				maxRate2 <- der1_2[iMaxRate2]
  				begPlat2 <- begPlat1 ; endPlat2 <- endPlat1
  				platSlope2 <- 0 # regPlat2$coefficients[2]
  				plateau2 <- mean(y2[begPlat2:endPlat2])
  				if (length(which(der1_2>0 & y2>plateau2 * thrLag))>0) {
  					iLag2 <- min(which(der1_2>0 & y2>plateau2 * thrLag))
  					lag2 <- x[iLag2]
  				} else {
  				  lag2 <- NA
  				  iLag2 <- -1
  				}
  				if (grOpt$lines) {
  			    ySlope2 <- (x-x[iMaxRate2])*maxRate2+y2[iMaxRate2]
  			    segSlope2 <- (max(0,iMaxRate2 - 
  			                          (iMaxRate2-iLag2)*0.65)):
  			        max(0,(iMaxRate2 + (iMaxRate2-iLag2)*0.65))
  			    if (G) points(x[segSlope2], ySlope2[segSlope2], type='l', 
  			           col=2, lty=2, lwd=3.5) # red dashed slope line (corrected)
  			    if (G) abline(v=x[iMaxRate2], lty=3, col=2) # red thin dashed line
  			    if (G) abline(h=plateau2, lt=2, lwd=2
  			             , col=4) # thick blue dashed plateau line
  			    if (G) abline(v=x[iLag2], lty=2, col=3, lwd=2) # thick green dashed line
  				}
  			} else {  # No flatPlateau
  				lag2 <- lag1
  				iLag2 <- iLag1
  				plateau2 <- NA
  				maxRate2 <- NA
  				iMaxRate2 <- iMaxRate1
  				y2 <- y1
  				if (grOpt$lines) {
  					ySlope <- (x-x[iMaxRate1])*maxRate1+y2[iMaxRate1]
  					segSlope <- (max(0,iMaxRate1 - 
  					                     (iMaxRate1-iLag1)*0.65)):
  					max(0,(iMaxRate1 + (iMaxRate1-iLag1)*0.65))
  					if (G) points(x[segSlope], ySlope[segSlope], type='l', 
  					       col=2, lty=2, lwd=3.5) # red dashed slope line (uncorrected)
  					if (G) abline(h=plateau1, lt=2, lwd=2, col=4)
  					if (G) abline(v=x[iLag1], lty=2, col=3, lwd=2)
  				}
  				if (G && grOpt$deriv) {
  				  points(x, der1_1+max(y1)*0.5, type="l", col=2, lwd=2) # red curve
  				  abline(h=max(y1)*0.5)
  				  points(x, 2*der2_1+max(y1)*0.5, type="l", col=3, lwd=2) # green curve
  				  abline(h=max(y1)*0.5)
  				}
  			}
  		} else {   # length(iBeforePlateau) == 0
  			plateau1 <- plateau2 <- NA
  			platSlope1 <- NA
  			lag1 <- lag2 <- NA
  			iLag2 <- iLag1 <- NA
  			maxRate2 <- maxRate1
  			iMaxRate2 <- NA
  			y2 <- y1
  		}
  		xText <- x[iMaxRate2]*1.1
  		if (is.null(yRange)) {
  		  yText <- max(y1)
  		} else yText <- max(yRange)
  		if (grOpt$text && flatPlateau) {
  			if (G && !is.na(maxRate1) && !is.na(maxRate2)) 
  			  text(xText, yText*0.25, paste0("Max rate: "
  			               , round(maxRate2,3), " ("
  			               , round(maxRate1,3), ") ",yUnit,"/h"), 
  			         adj=0, col=2)
  			if (G && !is.na(lag1) && !is.na(lag2)) {
  			  text(xText, yText*0.2, paste0("Lag: "
  			               , round(lag2,1)," ("
  			               , round(lag1,1),") hours"),  adj=0, col=3)
  			}
  			if (G && !is.na(plateau1) && !is.na(plateau2)) {
  			  text(xText, yText*0.15, paste0("Plateau: "
  			              , round(plateau2,2), " ("
  			              , round(plateau1,2),") ",yUnit,"/mL"), adj=0, col=4)
  			}
  		  if (G && ! is.na(snr1)) {
  		    text(xText, yText*0.1, paste0("S/N ratio: "
  		                , round(snr1,0),""), adj=0, col=1)
  		  }
  			if (G && !is.na(platSlope1)) {
  			  text(xText, yText*0.05, 
  			         paste0("Drift: "
  			             , round(platSlope1,4)," ",yUnit,"/hour"), adj=0)
  			}
  		}
  	  if (grOpt$text && ! flatPlateau) {
  	    if (G && !is.na(maxRate1)) {
  	      text(xText, yText*0.25, paste0("Max rate: "
  	                 , round(maxRate1,3), " ",yUnit,"/h"), adj=0, col=2)
  	    }
  	    if (G && !is.na(lag1)) {
  	      text(xText, yText*0.2, paste0("Lag: "
  	                 , round(lag1,1)," hours"), adj=0, col=3)
  	    }
  	    if (G && !is.na(plateau1)) {
  	      text(xText, yText*0.15, paste0("Plateau: "
  	                 , round(plateau1,2), " ",yUnit), adj=0, col=4)
  	    }
  	    if (G && ! is.na(snr1)) {
  	      text(xText, yText*0.1, paste0("S/N ratio: "
  	                 , round(snr1,0),"")
  			     , adj=0, col=1)
  	    }
  			if (G && ! is.na(platSlope1)) {
  			  text(xText, yText*0.05, paste0("Drift: "
  			             , round(platSlope1,4), " ",yUnit,"/h"), adj=0)
  			}
  	  }
  		res[1,"device"] <- unique(dat$device)[1]
  		res[1,"line"] <- paste0(well$line)
  		res[1,"col"] <- paste0(well$col)
  		res[1,"snr"] <- round(snr1,0)
  		if (flatPlateau) {
  		  res[1,"maxRateRaw"] <- round(maxRate1,3)
  		  res[1,"lagRaw"] <- round(lag1,1)
  		  res[1,"plateauRaw"] <- round(plateau1,2)
  		  res[1,"drift"] <- round(platSlope1,4)
  		  res[1,"maxRateCorr"] <- round(maxRate2,3)
  			res[1,"lagCorr"] <- round(lag2,1)
  		  res[1,"plateauCorr"] <- round(plateau2,2)
  		} else {
  		  res[1,"maxRate"] <- round(maxRate1,3)
  		  res[1,"lag"] <- round(lag1,1)
  		  res[1,"plateau"] <- round(plateau1,2)
  		}
  	  allRes <- rbind.data.frame(allRes, res)
	  }  #  end if ix in wells
	}  #  end loop on wells
	if (writose) {
	  write.table(allRes, paste0(resName,".txt"), row.names=FALSE)
	  if (verbose) cat(paste0("\nGrowth parameters saved as: "
	                        ,paste0(getwd(),"/",resName, ".txt\n")))
	}
	if (G && !is.null(pdfName)) {
	  dev.off()
	  if (verbose) cat(paste0("Graphics written in file:\n",
	                          getwd(),"/",pdfName,".pdf"))
	}
	return(allRes)
}


#' @name PlotAllRepsWellsDevices
#' @title Plots a global picture of 96 MiniRead output curves
#' @description Each curve can be either a kinetics curve (time plotted as 
#' X values) or a calibration curve (calibration values plotted as X values).\cr
#' If the column 'rep' contains several different values, the different
#' replicates will be plotted on the same graph, together with their average 
#' curve.\cr
#' If the column 'device' contains several different values (e.g. "A,"B",...)
#' then the output file will contain several pages, one for each device
#' @details The graphs can be written as a PDF file or on the current graphic 
#' device
#' @inheritParams ReadMiniReadData
#' @param linReg Logical. If TRUE, linear regression will be applied on the 
#' (average) curves
#' @param pdfName Character string. Name of the PDF file generated with the
#' graphs. If 'pdfName' is NULL, the plot will be sent to the current graphical 
#' device
#' @param xValue Character string. 
#' - If 'xValue'=="hours", the column 'hours' of the input table will be 
#' used as X values for the plots
#' - If 'xValue'=="calib", the column 'calib' of the input table will be 
#' used as X values for the plots
#' @param xLabel Character string. Text to be displayed as X-axis legend in
#' the graphs
#' @param yLabel Character string. Text to be displayed as Y-axis legend in
#' the graphs
#' @return Data frame (if linReg is TRUE) or NULL (else). The returned data
#' frame has one line per graph and the following columns:
#' - device: Character. Label of the MiniRead device,
#' - i_well: Numeric. Index of the well
#' - well: Character string. Well coordinate (e.g. "A1")
#' - slope: Numeric. Slope of the linear regression
#' - intercept: Numeric. Intercept of the linear regression
#' - R2: Numeric. R-squared of the linear regression
#' - p_value: Numeric. P-value of the linear regression
#' @examples
#' caliReps <- ReadMiniReadData(ExampleFile("CALI_REPS"))
#' reg <- PlotAllRepsWellsDevices(mrDataOrFile=caliReps, 
#'                                xValue="calib")
#' @export
#
PlotAllRepsWellsDevices <- function(mrDataOrFile, 
                              xValue=NULL, 
                              linReg=TRUE, 
                              verbose=TRUE,
                              pdfName="Raw_Values_Multiple_Devices_Wells", 
                              xLabel="Calibration value",
                              yLabel="MiniRead output") { 
  data <- ReadMiniReadData(mrDataOrFile=mrDataOrFile)
  if ( ! is.null(xValue) && ! xValue %in% c("calib","hours")) {
    stop(" in 'PlotAllRepsWellsDevices()': invalid argument 'xValue'. Must be 'calib' or 'hours'")
  }
  if (is.null(data$device)) data <- cbind.data.frame(data, device="MiniRead")
  wells <- paste0("W",1:96)
  if ( ! is.null(pdfName)) {
    pdf(paste0(pdfName,".pdf"), 36, 24)
  }
  par(mfrow=c(8,12))
  dev <- unique(data$device)[1]
  regPar <- NULL
  for (dev in unique(data$device)) {
    datDev <- data[data$device==dev,]
    agg <- suppressWarnings(aggregate(x=datDev, by=list(datDev[,xValue]), FUN=mean))
    x <- unique(datDev[,xValue])
    xLim <- c(0,max(x))
    yLim <- range(datDev[,wells])
    iWell <- 1
    for (iWell in 1:96) {
      col <- (iWell-1) %% 12 + 1
      line <- c("A","B","C","D","E","F","G","H")[floor((iWell-1)/12) + 1]
      xLab <- xLabel
      yLab <- yLabel
      main <- paste0(line, col, "   MiniRead ",dev)
      iRep <- 1
      colReps <- c(2,3,8,4,7)
      for (iRep in sort(unique(datDev$rep))) {
        devRep <- datDev[datDev$rep==iRep,]
        x <- devRep[,xValue]
        y <- devRep[,wells[iWell]]
        if (iRep==sort(unique(datDev$rep))[1]) {
          plot(x, y, type="p", col=iRep+1, xlim=xLim, ylim=yLim
               , main=main, xlab=xLab, ylab=yLab)
        } else points(x, y, type="p", col=colReps[iRep])
      }
      if (length(unique(datDev$rep))>1) { # several reps
        legend(x=0, y=yLim[2], 
               legend=paste("Rep", sort(unique(datDev$rep))),
               col=colReps, pch=1)
      } 
      y2 <- agg[, wells[iWell]]
      points(x, y2, type="l", lty=2, col=1)
      if (linReg) {
        reg <- lm(y2~x)
        abline(reg, col=2, lty=1)
        slope <- reg$coefficients[2]
        inter <- reg$coefficients[1]
        r2 <- summary(reg)$r.squared
        pval <- summary(reg)$coefficients[2,4]
        text(x=max(x)*0.6, y=min(y), 
             labels=paste0("Y = ", round(slope,3), " * X + ", 
                           round(inter,3), adj=c(1,1)))
        df <- data.frame(device=dev, i_well=iWell, well=paste0(line,col), 
                         slope=slope, intercept=inter,
                         R2=r2, p_value=pval)
        regPar <- rbind.data.frame(regPar, df)
      }
    }
  }
  row.names(regPar) <- NULL
  if ( ! is.null(pdfName)) {
    dev.off()
    if (verbose) cat(paste0("Graphics written in file:\n",getwd()
                            ,"/",pdfName,".pdf"))
  }
  if (linReg) return(regPar)
}


#' @name WellNames2WellCoords
#' @title Converts  well names ("W1".."W96") into well coordinates (A1-H12)
#' @description Can handle vectors
#' @param wellNames Vector of Character strings. Must be of the forme 'Wxx'
#' where xx is the number of the well from 1 to 96. Numbers follow the 
#' order: 1(A1)-12(A12), 13(B1)-24(B12),...84(H1)-96(H12)
#' @return List. Elements are:
#' - coord: Vector of Character strings. Coordinate of the plate well
#' - line: Vector of Character strings. Row (A to H) of the plate well
#' - col: Vector of Character strings. Column (1 to 12) of the plate well
#' @examples
#' WellNames2WellCoords(wellNames=c("W1","W2","W96"))
#' @export
#
WellNames2WellCoords <- function(wellNames="W1") {
  wellNames <- CheckWellsList(wells=wellNames) # to check validity
  i <- 1
  wellCoords <- lines <- cols <- rep(NA, length(wellNames))
  for (i in 1:length(wellNames)) {
    iWell <- as.numeric(substr(wellNames[i],2,3))
    col <- (iWell-1) %% 12 + 1
    line <- c("A","B","C","D","E","F","G","H")[floor((iWell-1)/12) + 1]
    wellCoords[i] <- paste0(line, col)
    lines[i] <- line
    cols[i] <- col
  }
  return(list(coord=wellCoords, line=lines, col=cols))
}


#' @name WellCoords2WellNames
#' @title Converts well coordinates (A1-H12) into well names ("W1".."W96")
#' @description Can handle vectors
#' @param wellCoords List or Vector of Character strings. 
#' - if list, must be as returned by \code{\link{WellNames2WellCoords}()}
#' - if vector of character strings, vector of well coordinates 
#' e.g. c("A1","H12")
#' @return Well number of the form 'Wxx' as for parameter 'wellNames' of
#' \code{\link{WellNames2WellCoords}()}
#' @examples
#' WellCoords2WellNames(wellCoords=c("A1","B2","H12"))
#' @export
#
WellCoords2WellNames <- function(wellCoords) {
  well <- wellCoords
  if (is.list(well)) {
    lines <- well$line
    cols <- well$col
    if (is.null(lines) || is.null(cols)) stop("If well is a list, it must 
                                              be list(line=.., col=..)")
    if (length(lines) != length(cols)) stop("well$line and well$col have 
                                            different lengths")
    nb <- length(lines)
  } else if (is.character(well)) {
    cols <- as.numeric(substr(well,2,3))
    lines <- substr(well,1,1)
    valLines <- c("A","B","C","D","E","F","G","H")
    if (sum(! cols%in%1:12)>0 || sum(! lines%in%valLines)>0) stop(paste0("lines 
        must be named by a letter (A to H) and columns by an integer 1-12"))
    nb <- length(well)
  } else stop(paste0("Wells must be named either by a vector of strings ('A1, 
                     'B2',...) or a list(line=  , col=  )"))
  iWell <- rep(NA, nb)
  i <- 1
  for (i in 1:nb) {
   iWell[i] <- paste0("W",12*(which(valLines==lines[i])-1) + cols[i])
  }
  return(iWell)
}


#' @name AggregateReplicates
#' @title Aggregates replicates of MiniRead data with a given function
#' @description Uses the standard aggregate() function 
#' @inheritParams IsMiniReadData
#' @param FUN Function. Function to apply to aggregate the data, as required 
#' for the base function \code{\link{aggregate}()}
#' @return Aggregated data with only one line per value in 
#' fields 'hours' and/or 'calib'
#' @examples
#' caliReps <- ReadMiniReadData(ExampleFile("CALI_REPS"))
#' dim(caliReps)
#' caliReps$rep
#' caliReps[1:20 , 1:7]
#' meanRep <- AggregateReplicates(mrData=caliReps, FUN=mean)
#' dim(meanRep)
#' meanRep[1:20 , 1:7]
#' @export
AggregateReplicates <- function(mrData, FUN=mean) {
  agg <- aggregate(
    x=as.matrix(mrData[,c(1:4, 6:105)])~mrData$calib+mrData$device,
    FUN=FUN)
  agg <- cbind.data.frame(agg[,3:6], device=agg[,2], agg[7:106])
  return(agg)
}


#' @name HomogeneityPlot
#' @title Analyze homogeneity of growth parameters over the wells of plates
#' @description Draws heat maps showing the values of growth parameters 
#' in each well of each MiniRead instrument
#' @details The aim of this plot is to detect possible environmental
#' gradients of growth conditions within a plate, possibly due to heterogeneity
#' of temperature or evaporation e.g. between the borders and the center of
#' the plate
#' @param growthData Data frame. Lines and columns must be the same as for the 
#' output value of \code{\link{AnalyzeGrowth}()}
#' @param suffix Character string. Text added at the end of the graph titles
#' @param values Vector of character strings. Lists the column names from the 
#' 'growthData' data frame that will be taken as growth parameters to 
#' represent in the heat maps
#' @param valLabs Vector of character strings. Lists the text that will be 
#' displayed in the header of each graph to describe which growth parameter
#' is plotted. length(valLabs) must be equal to length(values)
#' @param pdfName Character string. Name of the pdf file that will be generated
#' with the heat maps. If NULL then graphs will be sent to the current graphic
#' device
#' @return NULL
#' @examples
#' homog <- read.table(ExampleFile("HOMOG"), header=TRUE)
#' HomogeneityPlot(growthData=homog)
#' @export
HomogeneityPlot <- function(growthData, suffix=""
                    , values=c("maxRateRaw","lagRaw","plateauRaw","snr","drift")
                    , valLabs=c("Max growth rate","Lag time","Plateau"
                               ,"S/N ratio","Drift")
                    , pdfName="Growth_homogeneity.pdf") {
  gd <- growthData
  if (length(valLabs) != length(values)) {
    stop(" in HomogeneityPlot(): 'values' and 'valLabs' must have the same size")
  }
  devices <- unique(gd$device)
  if ( ! is.null(pdfName)) {
    pdf( pdfName, 7*length(values), 3+3*length(devices))
  }
  iDev <- 1
  cc <- rev(rainbow(256, start=0, end=0.85))
  rowLab <- rep(c("A","B","C","D","E","F","G","H"),length(devices))
  devLab <- sort(rep(devices, 8))
  colLab <- 1:12
  par(mfrow=c(1,length(values)), mar=c(12,7,7,7))
  charSize <- 2
  iVal <- 1
  for (iVal in 1:length(values)) {
    val <- values[iVal]
    rangeVal <- range(gd[,val], na.rm=T)
    norm <- (gd[,val]-rangeVal[1]) / diff(rangeVal)
    mat <- matrix(norm, nrow=8*length(devices), ncol=12, byrow=TRUE)
    ttmat <- t(mat)[ , rev(1:dim(t(mat))[2])]
    image( x=1:12, y=1:(8*length(devices)), z=ttmat, xlab="", ylab=""
           , col=cc, axes=FALSE)
    text(x=6.5
         , y=8*length(devices)+1.5
         , labels=paste0(valLabs[iVal], "  ", suffix)
         , xpd=TRUE
         , adj=c(0.5, 0.5)
         , srt=0, cex=charSize*2)
    text(x=(1:12)+0.25
         , y=0 # 8*length(devices)+1
         , labels=colLab
         , xpd=TRUE
         , adj=c(1, 0.5)
         , srt=0, cex=charSize)
    text(x=6.5
         , y=-0.6
         , labels="Plate column"
         , xpd=TRUE
         , adj=c(0.5, 0.5)
         , srt=0, cex=charSize)
    text(x=0
         , y=1:(8*length(devices))
         , labels=rev(rowLab)
         , xpd=TRUE
         , adj=c(1, 0.5)
         , srt=0, cex=charSize)
    text(x=-1
         , y=4.5+8*(0:(length(devices)-1))
         #         , labels=rep("Plate Row",length(devices))
         , labels="Plate Row"
         , xpd=TRUE
         , adj=c(0.5, 0.5)
         , srt=90, cex=charSize)
    text(x=13
         , y=1:(8*length(devices))
         , labels=rev(devLab)
         , xpd=TRUE
         , adj=c(0, 0.5)
         , srt=0, cex=charSize)
    text(x=14
         , y=4.5+8*(0:(length(devices)-1))
         , labels="Device ID"
         , xpd=TRUE
         , adj=c(0.5, 0.5)
         , srt=90, cex=charSize)
    text(x=1
         , y=-2
         , labels=paste0("Range: ",round(rangeVal[1],2)," - ",round(rangeVal[2],2))
         , xpd=TRUE
         , adj=c(0, 0.5)
         , srt=0, cex=charSize*2)
    abline(h=(0:length(devices))*8+0.5, lwd=3.5)
    abline(v=c(0.5, 12.5), lwd=3.5)
  }
  if ( ! is.null(pdfName)) {
    dev.off()
    cat(paste0("Graphics written in file:\n",getwd(),"/",pdfName,".pdf"))
  }
}
```
