## Supplementary material for "MiniRead: a simple and inexpensive do-it-yourself device for multiple analyses of micro-organism growth kinetics": R package: MiniRead_Tutorial.Rmd

````
---
title: "Tutorial of the MiniRead R Package"
output:
  pdf_document: default
  html_document: default
date: "2023-12-15"
---

```{r setup, include=TRUE, echo=FALSE}
# check if MiniRead is installed
if(!requireNamespace("MiniRead", quietly = TRUE)){
    stop('Please, install MiniRead package first e.g.:\ninstall.packages("MiniRead_4.1.3.tar.gz", type="source", repos=NULL)')
}
knitr::opts_chunk$set(echo = TRUE)
```

This tutorial does not show all possibilities of the MiniRead package. 
It only gives the most useful examples of analyses that the MiniRead users may 
need to do to be able to visualize raw data, perform calibration, and analyze 
growth parameters.   

For a complete list of functions and details about their possible 
arguments, rather use the package documentation and type '? function_name'
   
   
## 1. Visualize raw turbidity data (without calibration)

Load example kinetics data file path from the package
```{r}
library(MiniRead)
kineFile <- ExampleFile("KINE_B")
print(kineFile)
```
Read MiniRead output file into a data frame
```{r}
kineFile <- ExampleFile("KINE_B")
kineDat <- ReadMiniReadData(kineFile)
```
Here is what the output data frame looks like:
```{r}
head(kineDat, 2)
```

<br>
**Overview of 96 raw kinetics curves**
```{r echo=TRUE, fig.height=24, fig.width=36}
kineFile <- ExampleFile("KINE_B")
PlotAllRepsWellsDevices(mrDataOrFile=kineFile, 
                        xValue="hours", 
                        linReg=FALSE, 
                        pdfName=NULL,   # or =name_of_pdf_file
                        xLabel="Calibration value",
                        yLabel="MiniRead output")
```

<br>
**Plot raw data of two wells**   
* without growth parameter analysis   
* without correction for evaporation   
* without applying calibration curves   

```{r, fig.width=14, fig.height=7}
par(mfrow=c(1,2))
res <- AnalyzeGrowth(mrDataOrFile=kineFile, 
                     maxTime=48,
                     wells=c("A1","B2"),
                     useCalibration=FALSE,
                     pdfName=NULL,   # or =name_of_pdf_file
                     flatPlateau=FALSE,
                     writose=FALSE,
                     grOpt=list(text=FALSE, deriv=FALSE, lines=FALSE))
```

<br>
**Plot raw data of two wells**   
* *with growth parameter analysis*   
* without correction for evaporation   
* without applying calibration curves   

```{r, fig.width=14, fig.height=7}
par(mfrow=c(1,2))
res <- AnalyzeGrowth(mrDataOrFile=kineFile,
                     maxTime=48,
                     wells=c("A1","B2"),
                     useCalibration=FALSE,
                     pdfName=NULL,   # or =name_of_pdf_file
                     flatPlateau=FALSE,
                     writose=FALSE,
                     grOpt=list(text=TRUE, deriv=FALSE, lines=TRUE))
head(res)
```
<br>
**Plot raw data of two wells**   
* *with growth parameter analysis*   
* *with correction for evaporation*   
* without applying calibration curves   

```{r, fig.width=14, fig.height=7}
par(mfrow=c(1,2))
res <- AnalyzeGrowth(mrDataOrFile=kineFile, 
                     maxTime=48,
                     wells=c("A1","B2"),
                     useCalibration=FALSE,
                     pdfName=NULL,   # or =name_of_pdf_file
                     flatPlateau=TRUE,
                     writose=FALSE,
                     grOpt=list(text=TRUE, deriv=FALSE, lines=TRUE))
head(res)
```

<br>
**Writes in the current directory a pdf file with graphs for all 96 wells**   
* *with growth parameter analysis*   
* *with correction for evaporation*   
* without applying calibration curves   

```{r, fig.width=14, fig.height=7}
res <- AnalyzeGrowth(mrDataOrFile=kineFile, 
                     maxTime=48,
                     wells="all",
                     useCalibration=FALSE,
                     pdfName="Example_output_graphics",
                     flatPlateau=TRUE,
                     writose=TRUE,
                     grOpt=list(text=TRUE, deriv=FALSE, lines=TRUE))
```


## 2. Analyze calibration data to compute calibration curves (example with only one device)   

Load example calibration data file path from the package
```{r}
library(MiniRead)
calibFile <- ExampleFile("CALI_A")
```

Visualize all raw calibration values
```{r, fig.width=36, fig.height=24}
PlotAllRepsWellsDevices(mrDataOrFile=calibFile, 
                        xValue="calib", 
                        linReg=FALSE, 
                        pdfName=NULL,  # "Raw_Calibration_values_5_devices" 
                        xLabel="Calibration value",
                        yLabel="MiniRead output")
```

Real cell concentrations used for each point of the calibration series
```{r}
cellConcentrations <- c(cal0=0.00E+00, cal1=9.60E+01, cal2=1.96E+02, 
                        cal3=2.94E+02, cal4=3.77E+02)
```

Computes non-linear modelling of calibration curves
```{r, fig.width=14, fig.height=7}
calibCurves <- Calibrate(mrDataOrFile=calibFile,
                         calibValues=cellConcentrations,
                         pdfName=NULL)    # or =name_of_pdf_file
```
The function Calibrate() writes the computed calibration parameters 
(calibCurves, which contains the non-linear models of the calibration curves) 
in the folder 'MiniRead_Calibration' of your home directory.

Some information on the calibration data (including calibration date, 
labels of the MiniReads devices used for calibration, etc...) is attached
as attributes of the 'calibCurves' object:
```{r}
attributes(calibCurves)
```


## 3. Apply calibration curves to kinetics data (example with only one device)

Load example kinetics data (device "B")
```{r}
library(MiniRead)
kineFile <- ExampleFile("KINE_B")
nonCalibratedData <- ReadMiniReadData(kineFile)
```
Raw turbidity values (arbitrary units) recorded around the middle of expo phase:
```{r}
nonCalibratedData[150:160, c(1,8:13)]
```


Load calibration data for the same device ("B")
```{r}
calibFile <- ExampleFile("CALI_B")
```

Real cell concentrations (in Million Cells/mL) used for each point of the 
calibration series
```{r, fig.width=14, fig.height=7}
cellConcentrations <- c(cal0=0.00E+00, cal1=9.60E+01, cal2=1.96E+02, 
                        cal3=2.94E+02, cal4=3.77E+02)
```
Now computing calibration curves
```{r, fig.width=14, fig.height=7}
calibCurves <- Calibrate(mrDataOrFile=calibFile,
                         calibValues=cellConcentrations,
                         pdfName=NULL)    # or =name_of_pdf_file
```
And applying this calibration to the kinetics results:
```{r}
calibratedData <- ApplyCalibration(mrDataOrFile=kineFile, 
                                   calibrationCurves=calibCurves)
```

After conversion to Millions cells/mL by applying calibration curves,
the turbidity values (same points around the middle of expo phase) are:   
```{r}
calibratedData[150:160, c(1,8:13)]
```


## 4. Apply calibration curves to kinetics data (example with multiple devices)

Here, the same calibration plates (containing a five-points dilution
series of cell suspension) have been used to calibrate 5 different
MiniRead devices (named 'A' to 'E')

Load example kinetics data file path from the package
```{r}
library(MiniRead)
kineFile <- ExampleFile("KINE_B")
```

Load example calibration data of devices "A" to "E"
```{r}
calibFiles <- c(ExampleFile("CALI_A"), ExampleFile("CALI_B"), 
               ExampleFile("CALI_C"), ExampleFile("CALI_D"), 
               ExampleFile("CALI_E")
              )
```

Concatenate all data in a single data frame
```{r}
allCalis <- NULL
for (file in calibFiles) {
 thisCali <- ReadMiniReadData(file)
 allCalis <- rbind.data.frame(allCalis, thisCali)
}
```

Real cell concentrations used for each point of the calibration series
```{r}
cellConcentrations <- c(cal0=0.00E+00, cal1=9.60E+01, cal2=1.96E+02, 
                        cal3=2.94E+02, cal4=3.77E+02)
```

Compute and model calibration curves 
```{r}
calibCurves <- Calibrate(mrDataOrFile=allCalis,
                         calibValues=cellConcentrations)
```

The list of devices appears in the attributes:
```{r}
attributes(calibCurves)$calibDevices
```


## 5. Complete automatic analysis
This requires that calibration has already been carried out for the device(s) 
used, and that the calibCurves files have been written in the folder package.


**Plot well 'A1' of the plate**   
* *with growth parameter analysis*   
* *with correction for evaporation*   
* *with applying calibration curves*   

```{r, fig.width=7, fig.height=7}
res <- AnalyzeGrowth(mrDataOrFile=kineFile, 
                     maxTime=48,
                     wells="A1",
                     useCalibration=TRUE,
                     pdfName=NULL, 
                     flatPlateau=TRUE,
                     writose=FALSE,
                     grOpt=list(text=TRUE, deriv=FALSE, lines=TRUE))
head(res)
```
````
