## Supplementary material for "MiniRead: a simple and inexpensive do-it-yourself device for multiple analyses of micro-organism growth kinetics": R package: MiniRead_user-manual.pdf

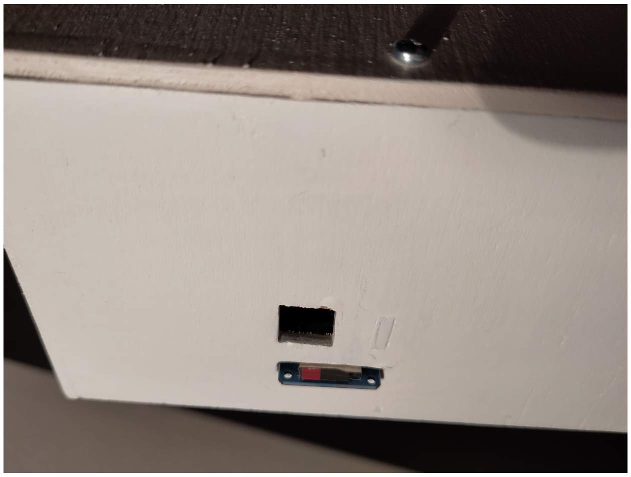

Figure 1: External SD card.

#### 2.5 Placing and removing the micro-SD card

When the MiniRead is doing turbidity acquisitions, the data are saved on the micro-SD card which is plugged on the left side of the device (see Figure 1 page 5). Press on the extremity of the card to insert or release it. Any small-capacity SD card can be used (16Gb is much more than enough !)
**CAUTION** : Never place or remove the SD card while the MiniRead is in action, it may corrupt data or even the card itself. Do it only while the interface displays 'IDLE'.
SD cards must be formatted as FAT32.

#### 2.6 Placing and removing plates

To place a culture plate in the device for reading,

- lift the front cover part up (using both hands...)

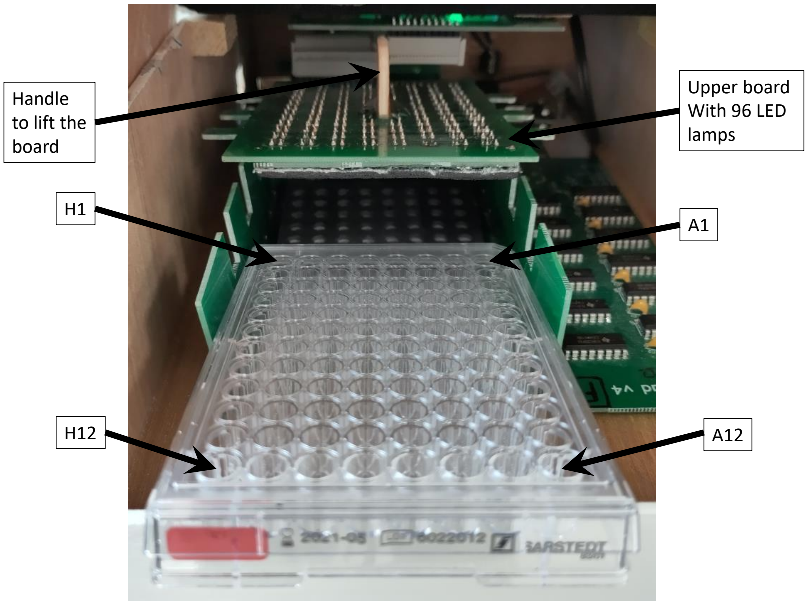

Figure 2: Upper board lifted to introduce the culture plate.

- lift the upper electronic board (with the 96 LED lamps) and push it slightly back until it holds on top of the plate housing, while leaving the space to introduce the plate (see Figure 2 page 5). To do so, hold the lid carefully by the wooden handle, lift it softly upwards without forcing (it would do nothing better than making it even more difficult to lift).

- gently place the plate in its housing, be careful to avoid splashing the medium from the wells. Check carefully that the plate is in direct contact with the bottom of the housing. Check also carefully the correct orientation of the plate, with 'A1-A12' row on your right side and 'H1-H12' on your left side.

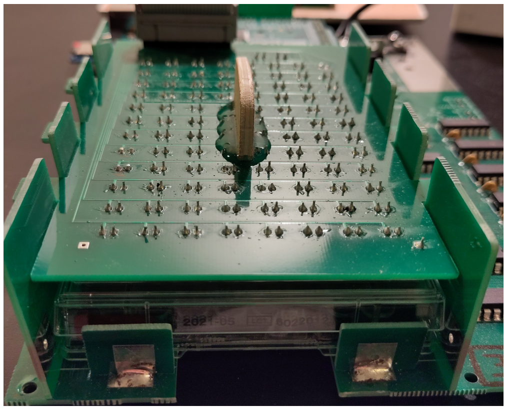

Figure 3: Culture plate correctly in place for reading.

- pull the upper electronic board slightly forward so it can fit again in its lateral slides, and gently press to slide it down. Make sure that the board comes in close contact to the lid of the culture plate (see Figure 3 page 6).

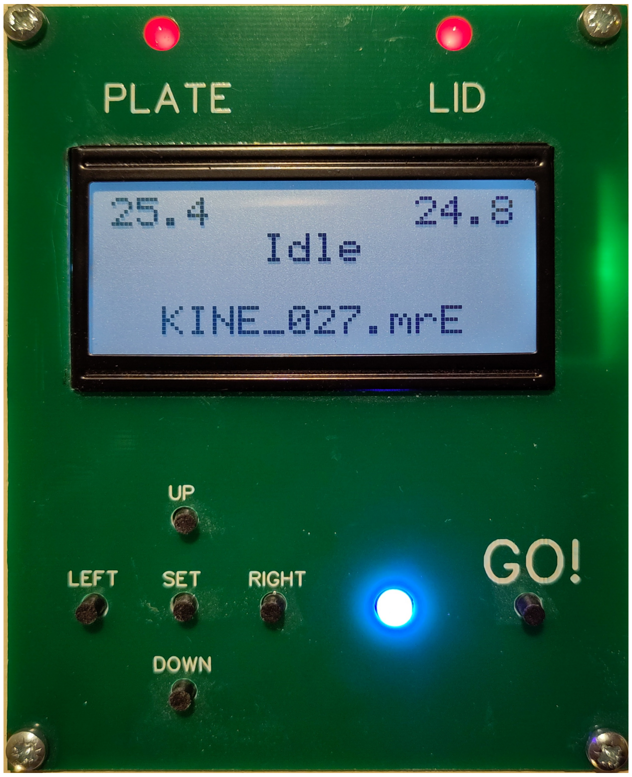

Figure 4: Interface board in Idle mode.

#### 4.3 Parameters setting

The MiniRead stores some parameters in its internal EEPROM memory, which is persistent event when the device is unplugged. To view or change the general settings of the MiniRead, press SET to enter the settings mode (see Figure 5 page 9).
The display shows the current values of the parameters settings. To change any of these values, use the LEFT and RIGHT buttons to navigate from one setting to another, and the TOP and DOWN buttons to increase or decrease the value (each digit separately). Then press SET again to validate the new values and/or come back to the Idle menu. The parameters are listed below (from top to bottom of the screen):

- set temperatures for bottom (PLATE) and top (LID) of the culture plate.
  ****IMPORTANT**** it is strongly advised to set the lid temperature two degrees Celsius higher than the temperature of the plate bottom, to avoid any condensation of water under the lid, which would completely alter the readings.
- name of the current program
- brightness value (from 1 to 255) for the 96 LED lamps used to measure turbidity. The optimal value may depend on the type of culture medium and the type of LEDs used to builf the device. The value 100 can be a good start.

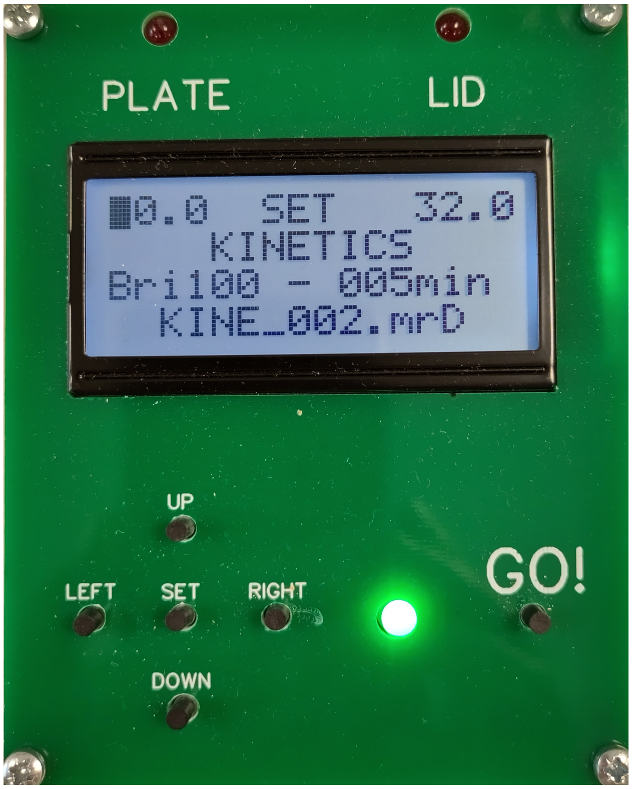

Figure 5: Interface in settings mode.

- time interval (in minutes) between two successive acquisitions during the KINE program (recording growth kinetics)
- name of the current data file (see details on file format and names in section 3.2).

- `HomogeneityPlot()`: draws heat-maps of the values of growth parameters across the different positions within a culture plate.

### 7 How to start with the MiniRead R package

****CAUTION****: trying to copy-paste code directly from a pdf file to a R console may insert many blank spaces in the commands and prevent them to work. A text-file version of this tutorial is available in the MiniRead project at ForgeMia (`https://forgemia.inra.fr/gqe-base/MiniRead`).

```

##############################################################
##                                                          ##
##              TUTORIAL of the MiniRead R PACKAGE          ##
##                                                          ##
##############################################################
# These are simple examples of basic use of the main functions
# For details and complete list of arguments, see User Manual
# or use '? function_name'
#
# Each section starting with library(MiniRead) may be run
# independently from the others

################################################################
# 1. Visualize raw turbidity data (without calibration)
################################################################
library(MiniRead)

# Load example kinetics data file path from the package
kineFile <- ExampleFile("KINE_B")

# Read MiniRead output file into a data frame
kineDat <- ReadMiniReadData(kineFile)
head(kineDat)

# Writes pdf file with overview of 96 raw kinetics curves
PlotAllRepsWellsDevices(mrDataOrFile=kineFile,
xValue="hours",
linReg=FALSE,
pdfName="Raw_Values_Multiple_Devices_Wells",
xLabel="Calibration value",
yLabel="MiniRead output")

# Plot of raw data without growth parameter analysis
par(mfrow=c(1,2))
res <- AnalyzeGrowth(mrDataOrFile=kineFile,
maxTime=48,
wells=c("A1","B2"),
useCalibration=FALSE,
pdfName=NULL,
flatPlateau=FALSE,
grOpt=list(text=FALSE, deriv=FALSE, lines=FALSE))

# Plot of raw data with growth parameters analysis
par(mfrow=c(1,2))
res <- AnalyzeGrowth(mrDataOrFile=kineFile,
maxTime=48,
wells=c("A1","B2"),
```

```
useCalibration=FALSE,
pdfName=NULL,
flatPlateau=FALSE,
grOpt=list(text=TRUE, deriv=FALSE, lines=TRUE))
head(res)
# and writes that in file Growth_kinetics_analysis_KINE_026_MRB.txt

# Plot with correction for medium evaporation
par(mfrow=c(1,2))
res <- AnalyzeGrowth(mrDataOrFile=kineFile,
maxTime=48,
wells=c("A1","B2"),
useCalibration=FALSE,
pdfName=NULL,
flatPlateau=TRUE,
grOpt=list(text=TRUE, deriv=FALSE, lines=TRUE))

# Writes a pdf file with graphs for all 96 wells
res <- AnalyzeGrowth(mrDataOrFile=kineFile,
maxTime=48,
wells="all",
useCalibration=FALSE,
pdfName="Example_output_graphics",
flatPlateau=TRUE,
grOpt=list(text=TRUE, deriv=FALSE, lines=TRUE))

###############################################################
# 2. Analyze calibration data to compute calibration curves
###############################################################
library(MiniRead)

# Load example calibration data file path from the package
calibFile <- ExampleFile("CALI_A")

# Real cell concentrations used for each point of the calibration series
cellConcentrations <- c(cal0=0.00E+00, cal1=9.60E+01, cal2=1.96E+02,
cal3=2.94E+02, cal4=3.77E+02)

# Computes non-linear modelling of calibration curves
# Writes file 'Calibration_MiniRead.pdf' with the calibration curves
calibCurves <- Calibrate(mrDataOrFile=calibFile,
calibValues=cellConcentrations)
# writes also the computed calibration parameters (calibCurves)
# (non-linear model of the calibration curves) in the package folder
# located in the path given in 'calibFile'

# Detailed information of the calibration data (including labels of the
# MiniReadis devices used for calibration) is attached as attributes
# of the 'calibCurves' object.
attributes(calibCurves)

###############################################################
```

```
# 3. Apply calibration curves to kinetics data (1 device)
##############################################################
library(MiniRead)

# Load example kinetics data (device "B")
kineFile <- ExampleFile("KINE_B")
nonCalibratedData <- ReadMiniReadData(kineFile)
# Raw turbidity values (arbitrary units) recorded (around middle of expo phase)
nonCalibratedData[150:160, 1:11]

# Load calibration data for the same device ("B")
calibFile <- ExampleFile("CALI_B")

# Real cell concentrations (in Million Cells/mL) used for each point of the
# calibration series
cellConcentrations <- c(cal0=0.00E+00, cal1=9.60E+01, cal2=1.96E+02,
cal3=2.94E+02, cal4=3.77E+02)

calibCurves <- Calibrate(mrDataOrFile=calibFile,
calibValues=cellConcentrations)

calibratedData <- ApplyCalibration(mrDataOrFile=kineFile,
calibrationCurves=calibCurves)

# Corresponding turbidity values after conversion to Millions cells/mL by
# applying calibration curves
calibratedData[150:160, 1:11]

####################################################################
# 4. Apply calibration curves to kinetics data (multiple devices)
####################################################################
# Here, the same calibration plates (containing a five-points dilution
# series of cell suspension) have been used to calibrate 5 different
# MiniRead devices (named 'A' to 'E')

library(MiniRead)

# Load example kinetics data file path from the package
kineFile <- ExampleFile("KINE_B")

# Load example calibraiton data of devices "A" to "E"
calibFiles <- c(ExampleFile("CALI_A"), ExampleFile("CALI_B"),
ExampleFile("CALI_C"), ExampleFile("CALI_D"),
ExampleFile("CALI_E")
)

# Concatenate all data in a single data frame
allCalis <- NULL
for (file in calibFiles) {
  thisCali <- ReadMiniReadData(file)
  allCalis <- rbind.data.frame(allCalis, thisCali)
```

```
}

# Visualizes all rawcalibration values
PlotAllRepsWellsDevices(mrDataOrFile=allCalis,
xValue="calib",
linReg=FALSE,
pdfName="Raw_Calibration_values_5_devices",
xLabel="Calibration value",
yLabel="MiniRead output")

# Real cell concentrations used for each point of the calibration series
cellConcentrations <- c(cal0=0.00E+00, cal1=9.60E+01, cal2=1.96E+02,
cal3=2.94E+02, cal4=3.77E+02)

# Computes and models calibration curves
calibCurves <- Calibrate(mrDataOrFile=allCalis,
calibValues=cellConcentrations)

# Detailed information of the calibration data (including labels of the
# MiniReadis devices used for calibration) is attached as attributes
# of the 'calibCurves' object.
attributes(calibCurves)

)
```
