## Supplementary figures and images for "MiniRead: a simple and inexpensive do-it-yourself device for multiple analyses of micro-organism growth kinetics"

### interfaceIdle.jpg

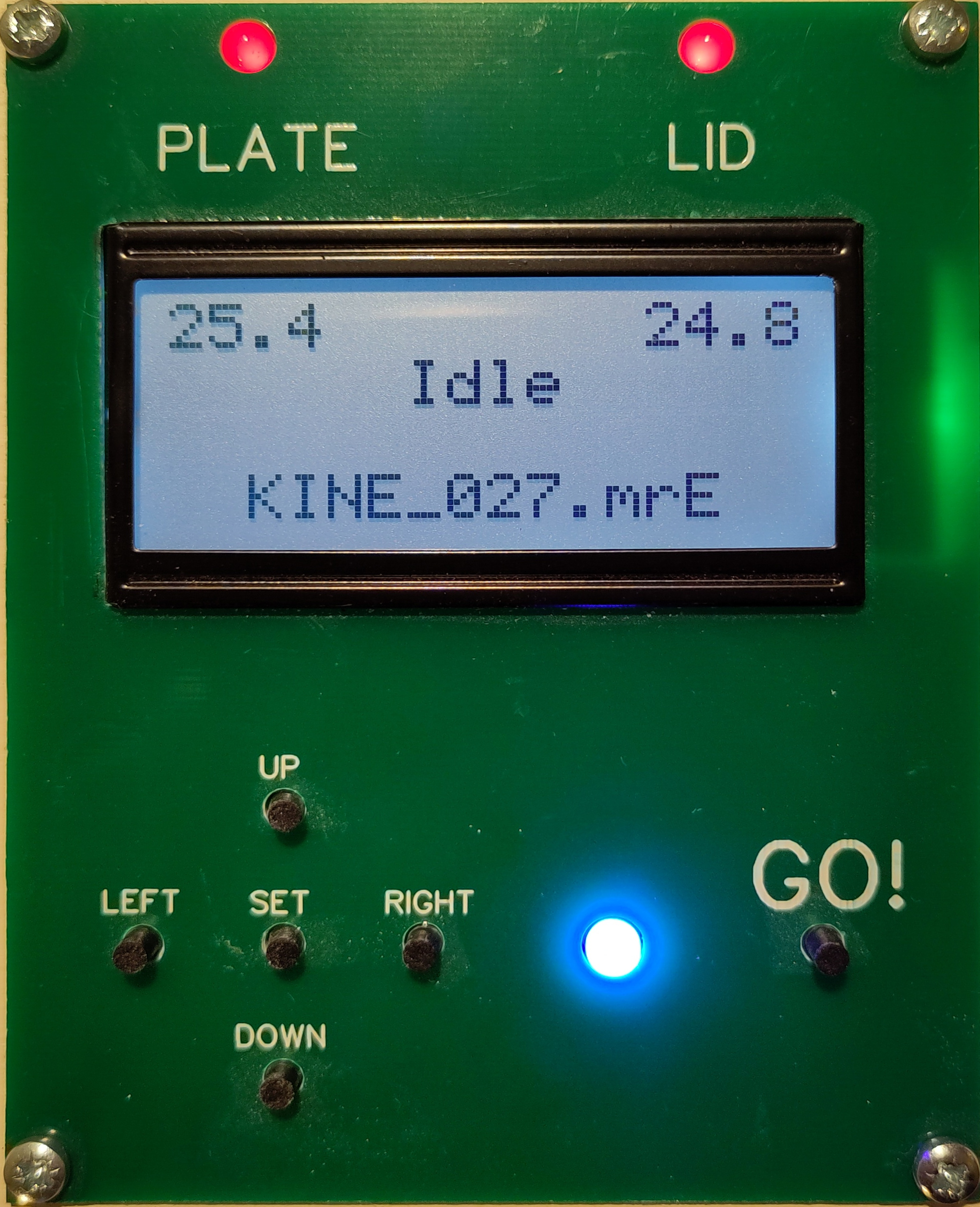

### interfaceRead.jpg

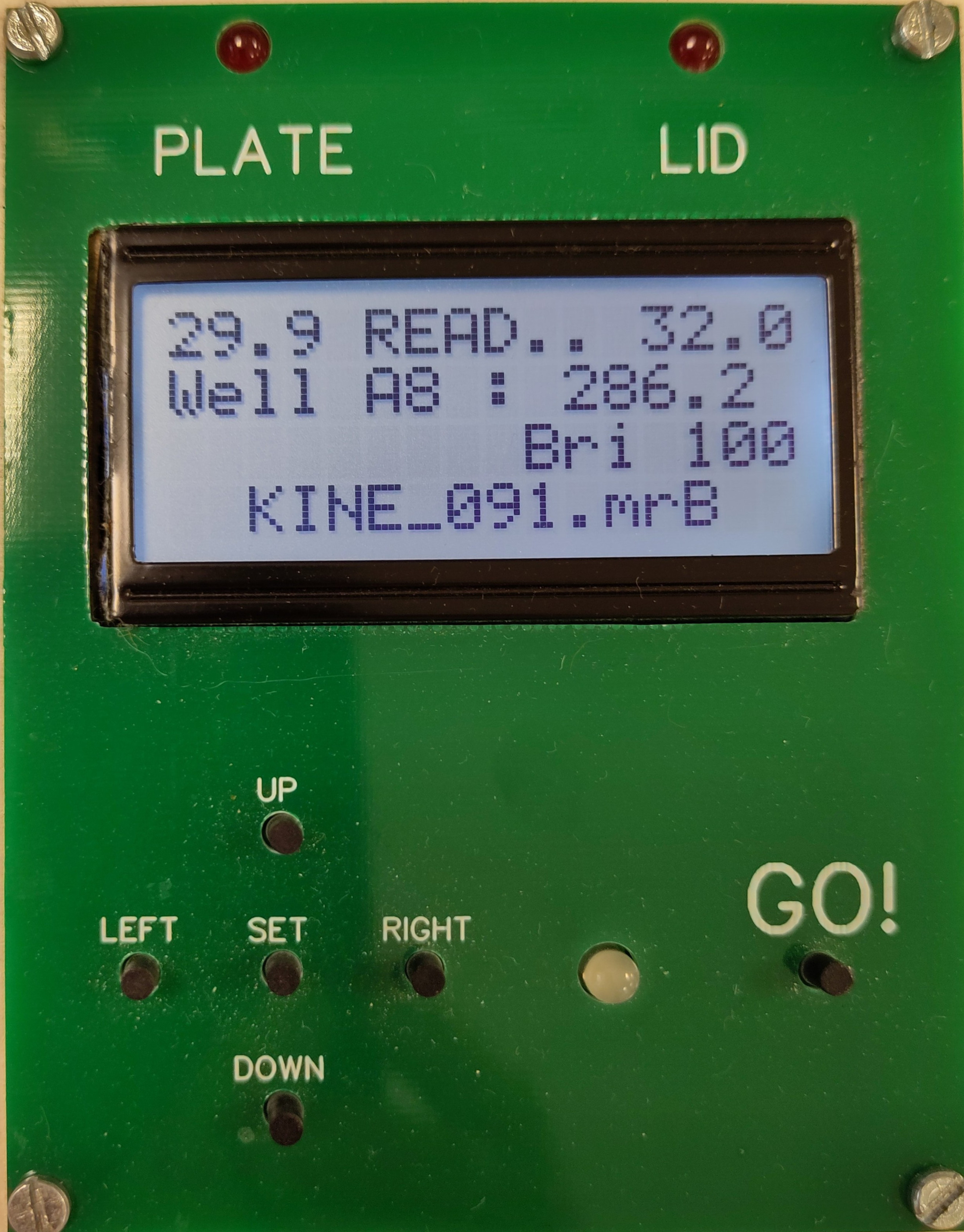

### interfaceSet.jpg

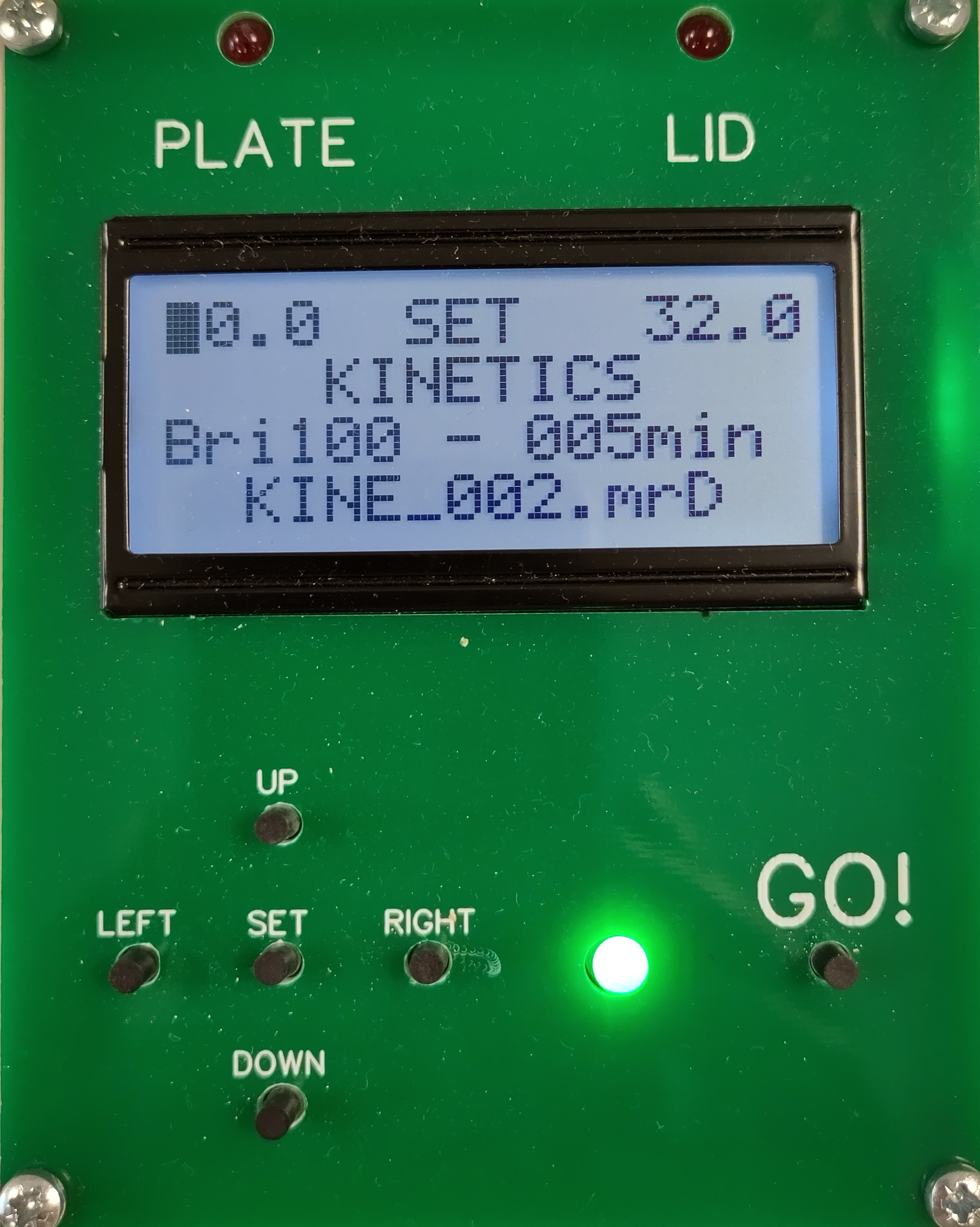

### PCB_PCB_MiniRead_v5_2023-02-15_All_Layers.pdf

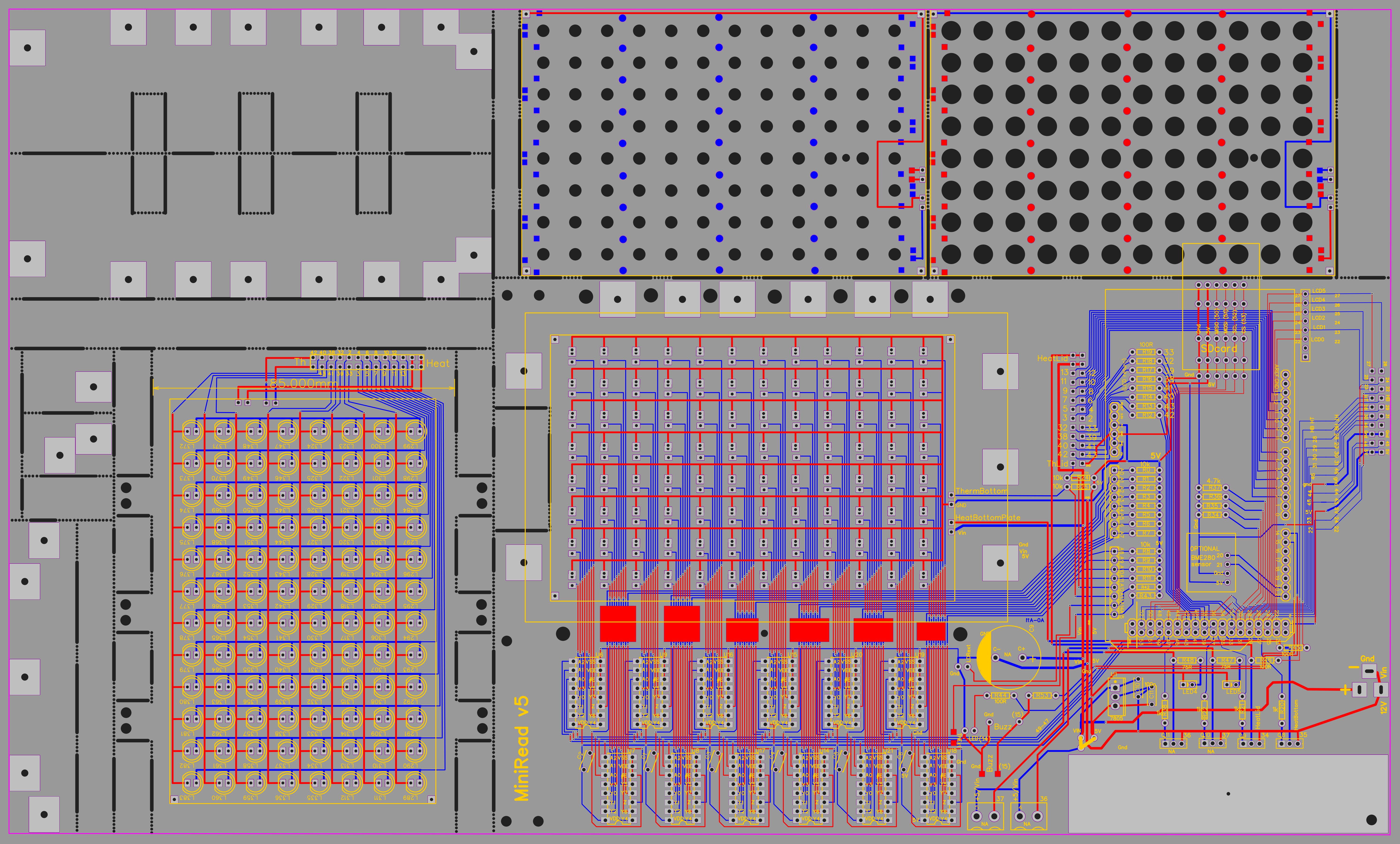

### PCB_PCB_MiniRead_v5_Display_2023-02-15_All_Layers.pdf

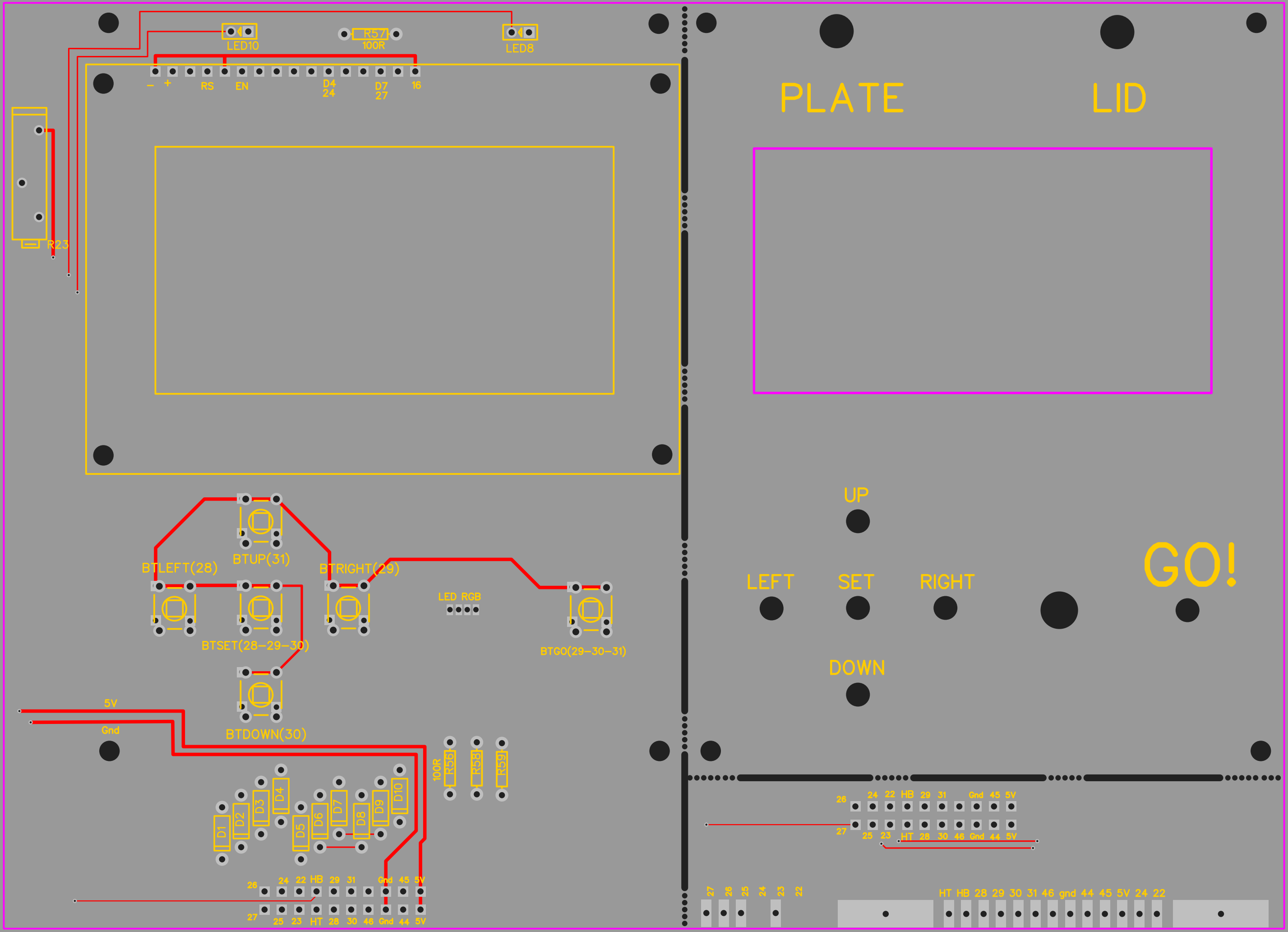

### plateInPlace.jpg

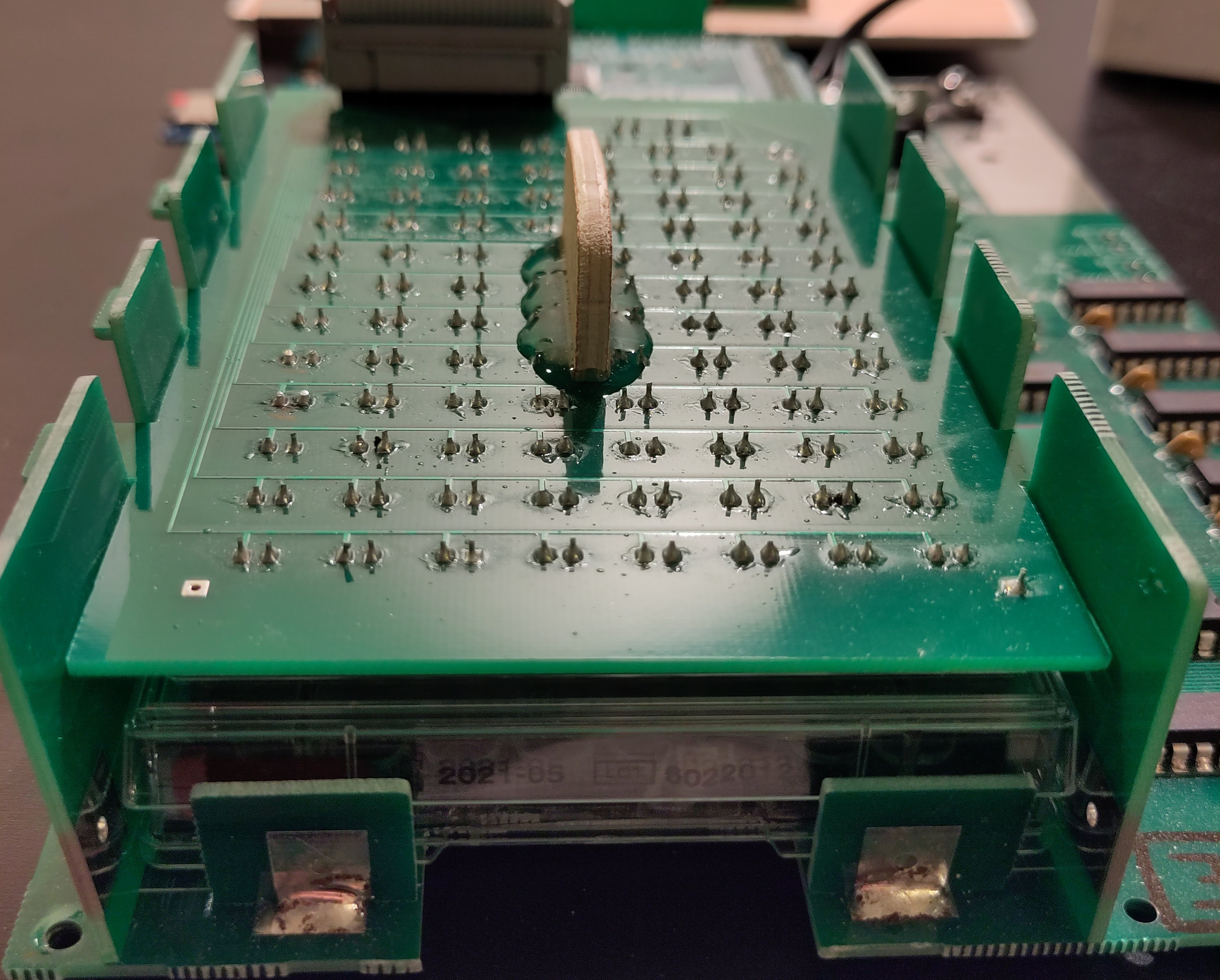

### plateUpperBoardUp.jpg

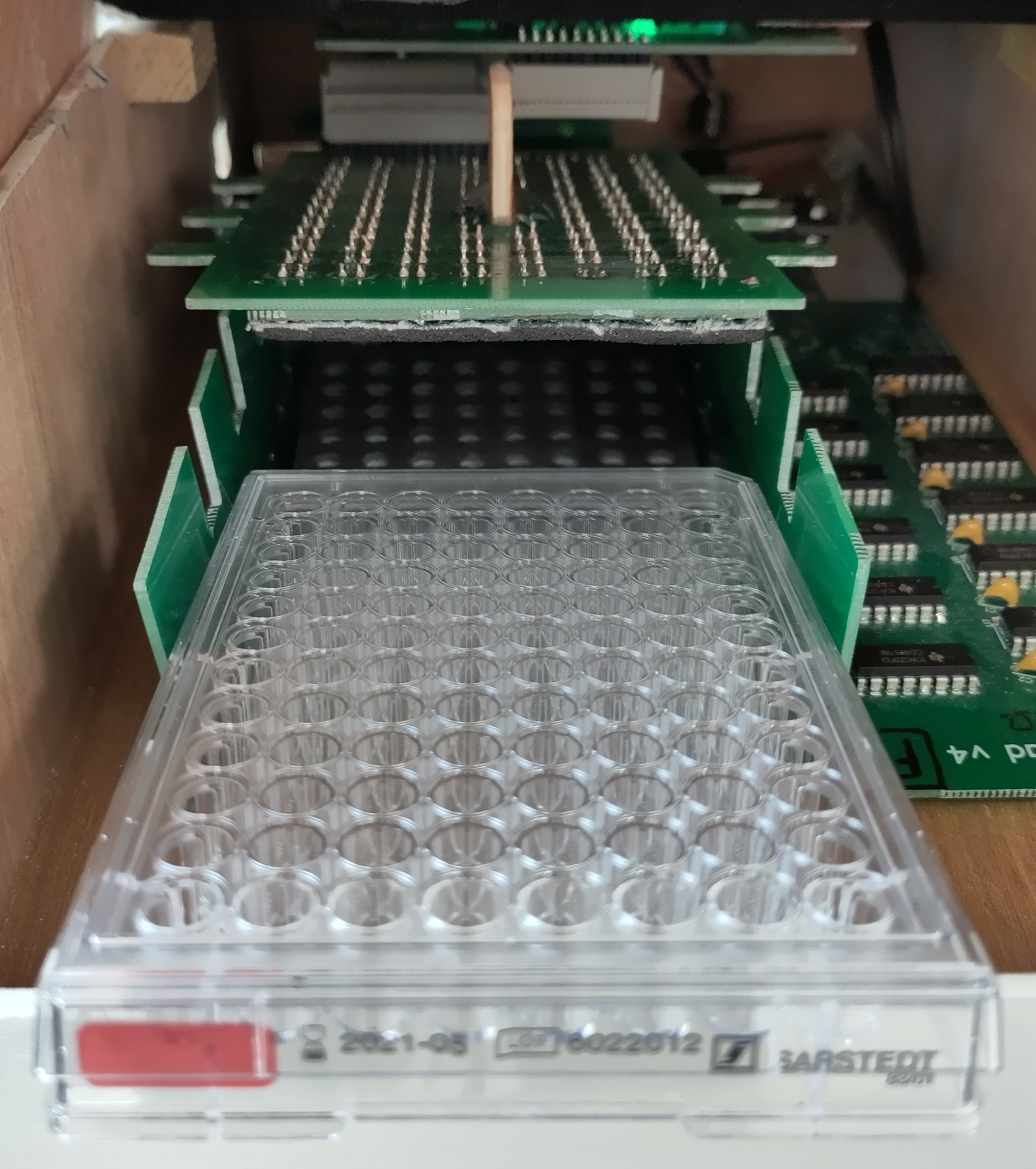

### plateUpperBoardUp.pdf

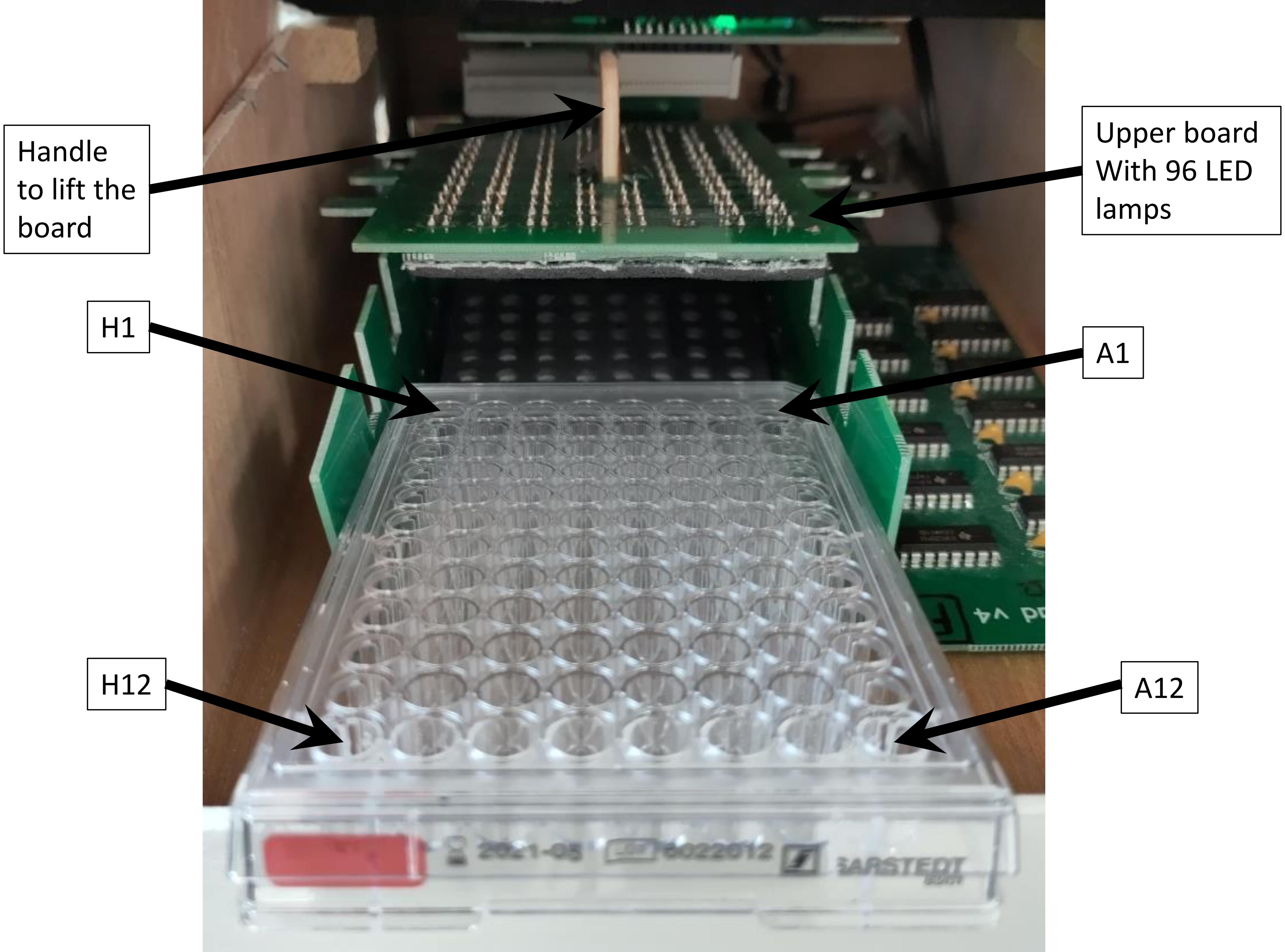

### sd.jpg

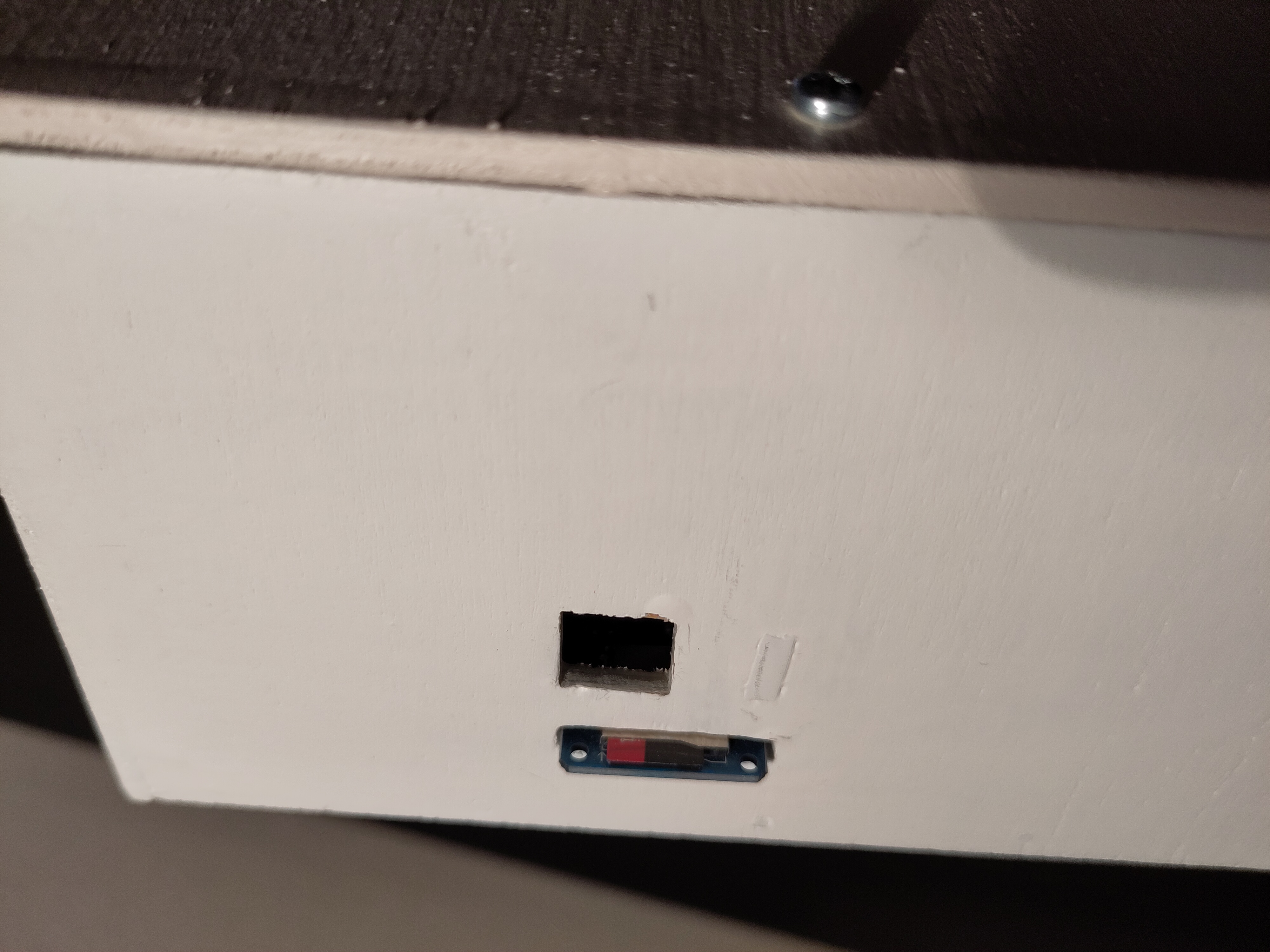
