## Supplementary material for "MiniRead: a simple and inexpensive do-it-yourself device for multiple analyses of micro-organism growth kinetics": Tutorial for R package

### Tutorial of the MiniRead R Package

2023-12-15

This tutorial does not show all possibilities of the MiniRead package. It only gives the most useful examples of analyses that the MiniRead users may need to do to be able to visualize raw data, perform calibration, and analyze growth parameters.

For a complete list of functions and details about their possible arguments, rather use the package documentation and type '? function_name'

#### 1. Visualize raw turbidity data (without calibration)

Load example kinetics data file path from the package

```
library(MiniRead)
kineFile <- ExampleFile("KINE_B")
print(kineFile)
```

```
## [1] "/home/mfalque/R/x86_64-pc-linux-gnu-library/4.3/MiniRead/extdata/KINE_026.MRB"
```

Read MiniRead output file into a data frame

```
kineFile <- ExampleFile("KINE_B")
kineDat <- ReadMiniReadData(kineFile)
```

```
##
## Reading MiniRead output file:
##  /home/mfalque/R/x86_64-pc-linux-gnu-library/4.3/MiniRead/extdata/KINE_026.MRB
```

Here is what the output data frame looks like:

```
head(kineDat, 2)
```

```
##     hours bright thBottom1 thLid1 device rep calib     W1     W2     W3     W4
## 1 0.00000    100     28.76  30.85      B   0     0 217.77 227.55 220.19 202.62
## 2 0.08378    100     30.12  31.87      B   0     0 216.37 227.12 219.22 202.02
##        W5     W6     W7     W8     W9    W10    W11    W12    W13    W14    W15
## 1 214.91 231.09 224.83 204.63 211.23 257.87 200.98 228.96 220.35 207.02 223.80
## 2 214.45 230.02 224.35 204.25 210.24 257.52 200.31 228.15 219.63 207.00 223.01
##       W16    W17    W18    W19    W20    W21    W22    W23    W24    W25    W26
## 1 215.01 268.25 222.18 286.96 290.18 240.60 195.15 199.92 322.43 219.97 210.64
## 2 215.00 268.16 222.41 286.40 289.75 239.97 194.69 199.38 321.69 219.16 209.96
##       W27    W28    W29    W30    W31    W32    W33    W34    W35    W36    W37
## 1 210.21 224.49 195.13 207.81 219.55 230.57 204.98 229.33 218.62 208.42 242.70
## 2 209.99 224.21 194.45 207.69 219.41 229.78 205.05 228.69 218.61 207.37 241.67
##       W38    W39    W40    W41    W42    W43    W44    W45    W46    W47    W48
## 1 235.93 215.11 250.71 197.29 220.24 237.89 208.45 234.05 218.64 200.14 241.28
## 2 235.25 215.06 250.57 197.26 220.91 237.63 209.14 233.87 219.25 200.25 241.81
##       W49    W50    W51    W52    W53    W54    W55    W56    W57    W58    W59
## 1 225.38 214.25 206.59 202.66 215.11 268.91 263.96 212.50 216.78 205.38 270.48
## 2 224.40 214.76 206.22 202.66 215.76 269.09 263.88 213.28 216.74 205.21 270.77
##       W60    W61    W62    W63    W64    W65    W66    W67    W68    W69    W70
```

```
## 1 198.85 311.68 219.34 232.23 228.33 213.04 218.99 227.76 201.94 217.40 237.66
## 2 199.20 310.96 218.71 232.36 228.39 212.72 219.66 227.52 202.41 218.13 237.79
##      W71    W72    W73    W74    W75    W76    W77    W78    W79    W80    W81
## 1 193.23 210.4 203.18 339.84 211.10 215.48 214.39 199.37 205.10 227.57 242.51
## 2 193.33 210.5 202.90 340.23 211.19 215.62 214.57 199.42 205.44 228.13 243.22
##      W82    W83    W84    W85    W86    W87    W88    W89    W90    W91    W92
## 1 197.35 210.99 217.36 292.51 224.54 212.93 220.10 228.16 236.22 209.83 199.77
## 2 197.75 211.16 217.88 292.29 224.30 212.86 220.17 228.25 236.58 209.67 199.79
##      W93    W94    W95    W96 thBottom2 thLid2
## 1 242.32 203.44 201.88 215.75     30.76  33.10
## 2 242.37 203.82 202.15 216.04     30.04  32.06
```

**Overview of 96 raw kinetics curves**

```
kineFile <- ExampleFile("KINE_B")
PlotAllRepsWellsDevices(mrDataOrFile=kineFile,
                        xValue="hours",
                        linReg=FALSE,
                        pdfName=NULL,    # or =name_of_pdf_file
                        xLabel="Calibration value",
                        yLabel="MiniRead output")
```

```
##
## Reading MiniRead output file:
##  /home/mfalque/R/x86_64-pc-linux-gnu-library/4.3/MiniRead/extdata/KINE_026.MRB
```

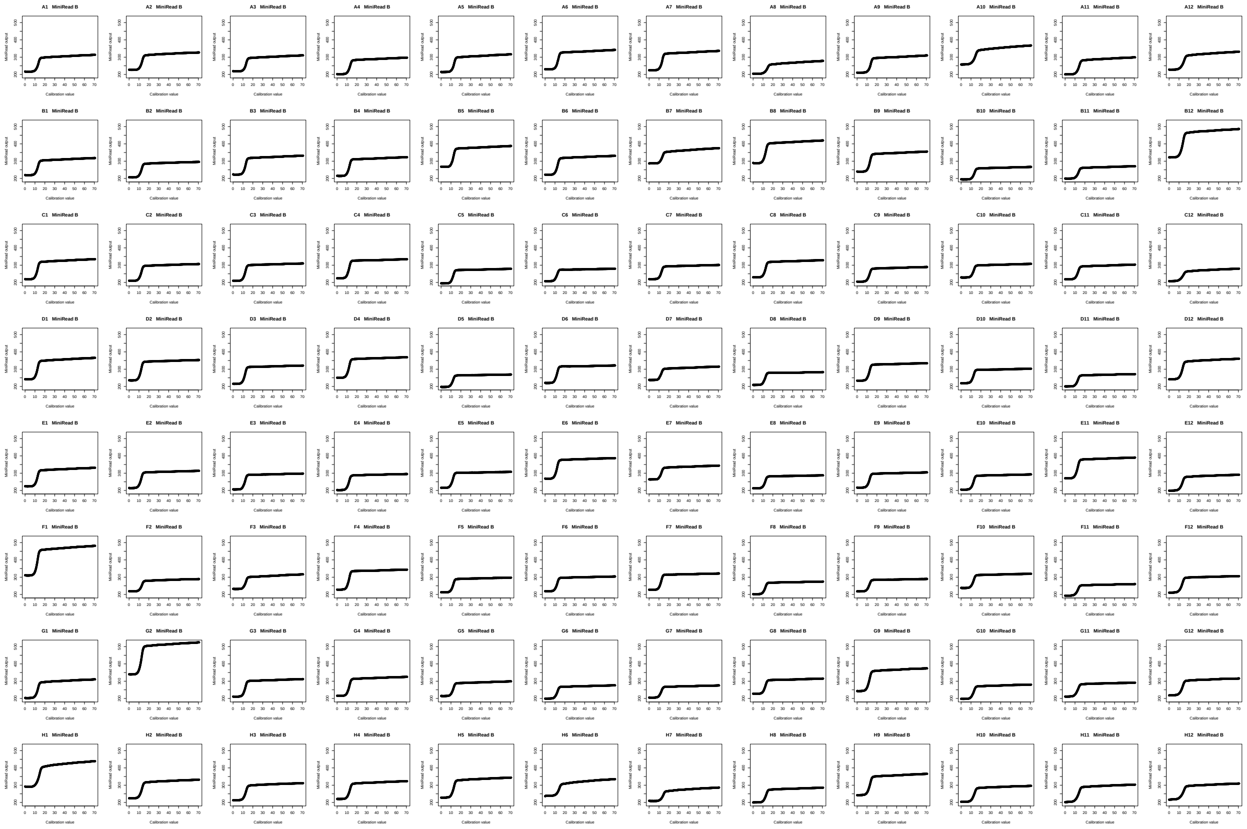

**Plot raw data of two wells**

* without growth parameter analysis
* without correction for evaporation

* without applying calibration curves

```
par(mfrow=c(1,2))
res <- AnalyzeGrowth(mrDataOrFile=kineFile,
                     maxTime=48,
                     wells=c("A1","B2"),
                     useCalibration=FALSE,
                     pdfName=NULL,    # or =name_of_pdf_file
                     flatPlateau=FALSE,
                     writose=FALSE,
                     grOpt=list(text=FALSE, deriv=FALSE, lines=FALSE))
```

```
##
## Reading MiniRead output file:
##  /home/mfalque/R/x86_64-pc-linux-gnu-library/4.3/MiniRead/extdata/KINE_026.MRB
## Checking format of data frame 'mrDataOrFile'
## Preparing plots and analyzing kinetics parameters
```

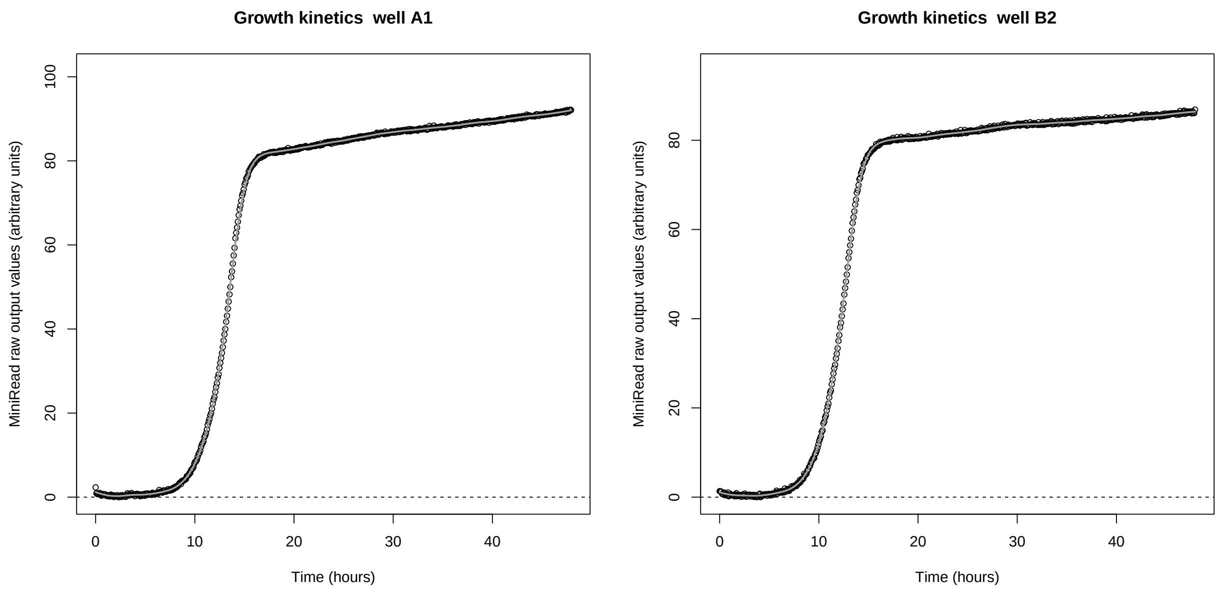


##### Plot raw data of two wells

* *with growth parameter analysis*
* without correction for evaporation
* without applying calibration curves

```
par(mfrow=c(1,2))
res <- AnalyzeGrowth(mrDataOrFile=kineFile,
                     maxTime=48,
                     wells=c("A1","B2"),
                     useCalibration=FALSE,
                     pdfName=NULL,    # or =name_of_pdf_file
                     flatPlateau=FALSE,
                     writose=FALSE,
                     grOpt=list(text=TRUE, deriv=FALSE, lines=TRUE))
```

```
##
## Reading MiniRead output file:
```

```
##  /home/mfalque/R/x86_64-pc-linux-gnu-library/4.3/MiniRead/extdata/KINE_026.MRB
## Checking format of data frame 'mrDataOrFile'
## Preparing plots and analyzing kinetics parameters
```

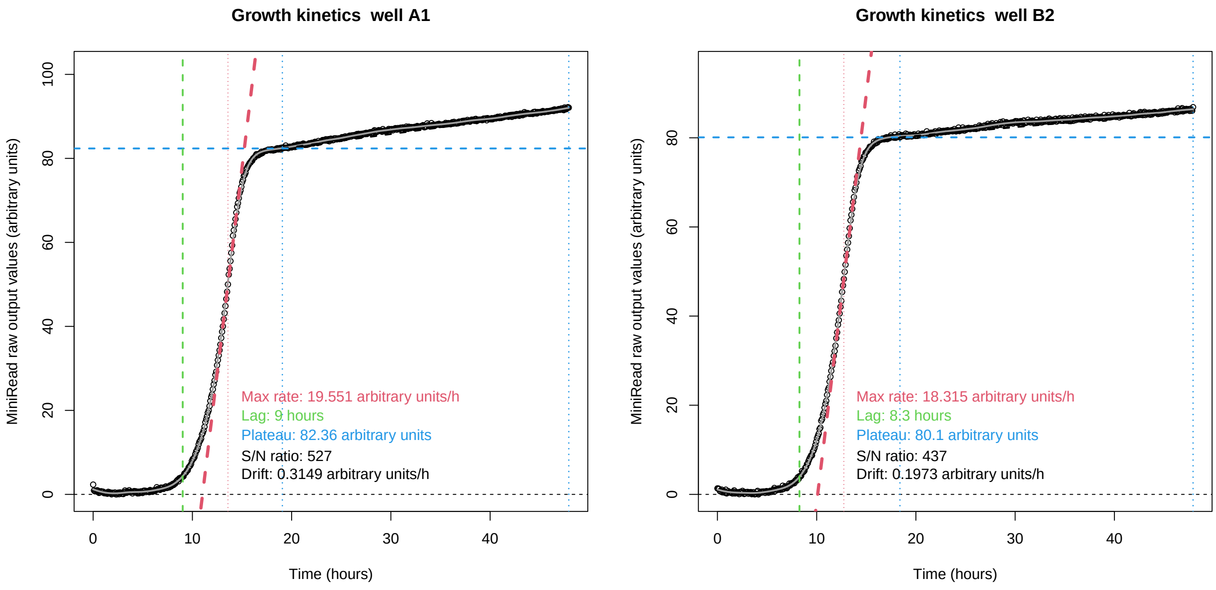


```r
head(res)
```

```
##   device line col maxRate lag plateau snr
## 1      B    A   1  19.551 9.0   82.36 527
## 2      B    B   2  18.315 8.3   80.10 437
```

**Plot raw data of two wells**
* *with growth parameter analysis*
* *with correction for evaporation*
* without applying calibration curves

```r
par(mfrow=c(1,2))
res <- AnalyzeGrowth(mrDataOrFile=kineFile,
                     maxTime=48,
                     wells=c("A1","B2"),
                     useCalibration=FALSE,
                     pdfName=NULL,    # or =name_of_pdf_file
                     flatPlateau=TRUE,
                     writose=FALSE,
                     grOpt=list(text=TRUE, deriv=FALSE, lines=TRUE))
```

```
##
## Reading MiniRead output file:
##  /home/mfalque/R/x86_64-pc-linux-gnu-library/4.3/MiniRead/extdata/KINE_026.MRB
## Checking format of data frame 'mrDataOrFile'
## Preparing plots and analyzing kinetics parameters
```

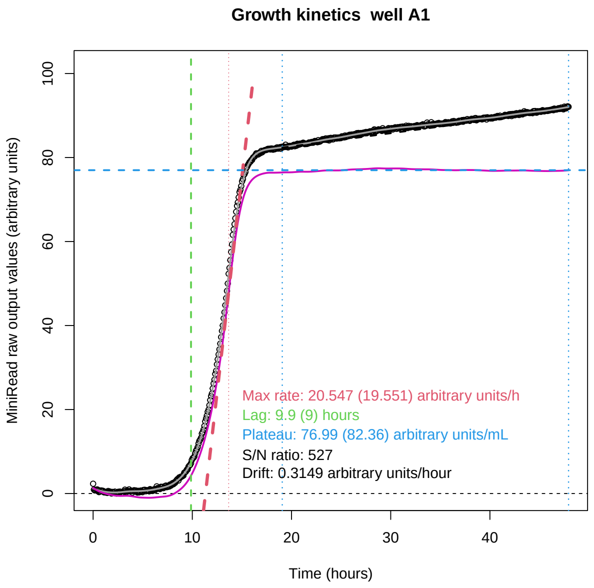


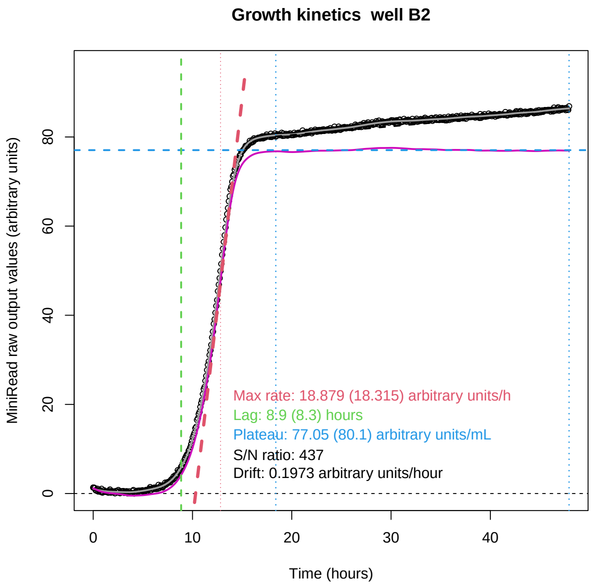


```r
head(res)
```

```
##   device line col maxRateRaw lagRaw plateauRaw snr  drift maxRateCorr lagCorr
## 1      B    A   1     19.551    9.0      82.36 527 0.3149      20.547     9.9
## 2      B    B   2     18.315    8.3      80.10 437 0.1973      18.879     8.9
##   plateauCorr
## 1       76.99
## 2       77.05
```

**Writes in the current directory a pdf file with graphs for all 96 wells**
* *with growth parameter analysis*
* *with correction for evaporation*
* without applying calibration curves

```r
res <- AnalyzeGrowth(mrDataOrFile=kineFile,
                     maxTime=48,
                     wells="all",
                     useCalibration=FALSE,
                     pdfName="Example_output_graphics",
                     flatPlateau=TRUE,
                     writose=TRUE,
                     grOpt=list(text=TRUE, deriv=FALSE, lines=TRUE))
```

```
##
## Reading MiniRead output file:
##  /home/mfalque/R/x86_64-pc-linux-gnu-library/4.3/MiniRead/extdata/KINE_026.MRB
## Checking format of data frame 'mrDataOrFile'
## Preparing plots and analyzing kinetics parameters
##
## Graphs saved as file Example_output_graphics.pdf

##
## Growth parameters saved as: /home/mfalque/DATA/GIT_DEV/MiniRead/user-manual/Example_output_graphics.
## Graphics written in file:
## /home/mfalque/DATA/GIT_DEV/MiniRead/user-manual/Example_output_graphics.pdf
```

#### 2. Analyze calibration data to compute calibration curves (example with only one device)

Load example calibration data file path from the package

```
library(MiniRead)
calibFile <- ExampleFile("CALI_A")
```

Visualize all raw calibration values

```
PlotAllRepsWellsDevices(mrDataOrFile=calibFile,
                        xValue="calib",
                        linReg=FALSE,
                        pdfName=NULL,  # "Raw_Calibration_values_5_devices"
                        xLabel="Calibration value",
                        yLabel="MiniRead output")
```

```
##
## Reading MiniRead output file:
##  /home/mfalque/R/x86_64-pc-linux-gnu-library/4.3/MiniRead/extdata/CALI_053.MRA
```

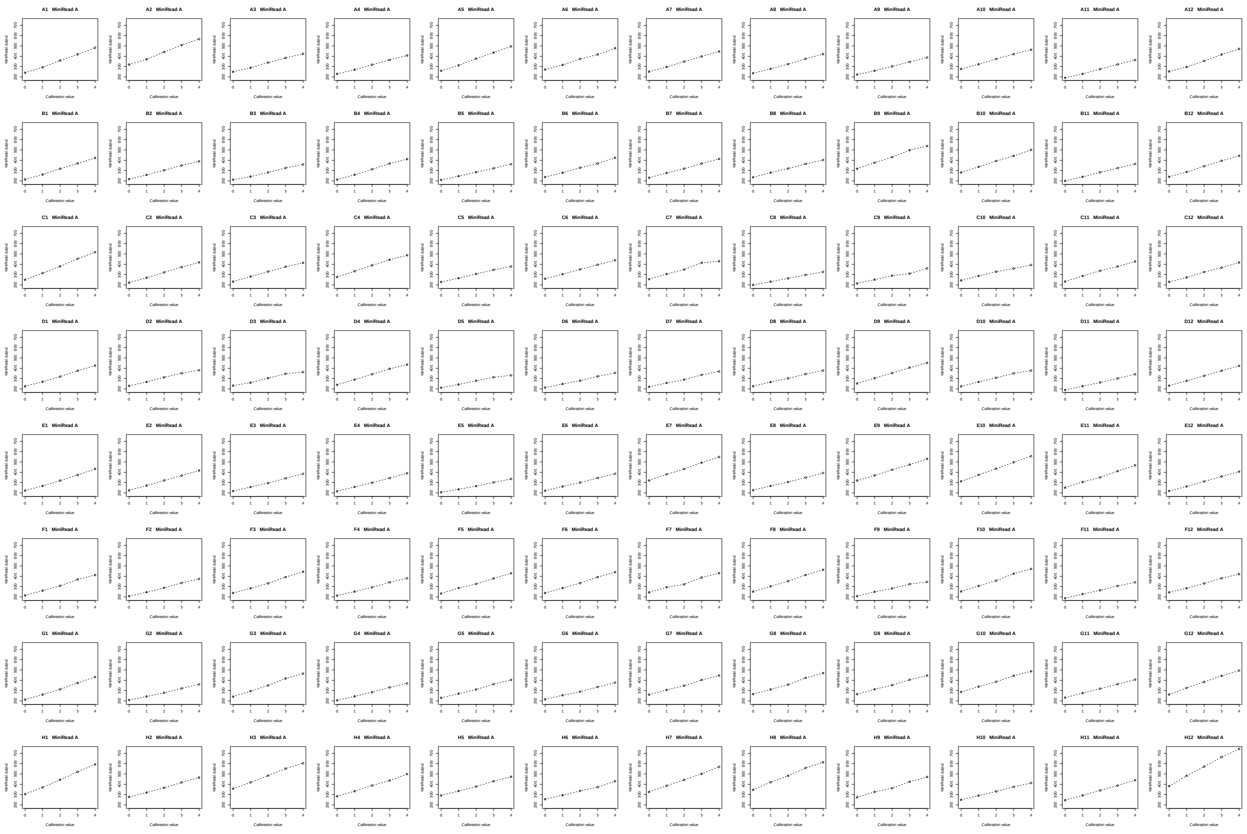

Real cell concentrations used for each point of the calibration series

```
cellConcentrations <- c(cal0=0.00E+00, cal1=9.60E+01, cal2=1.96E+02,
                        cal3=2.94E+02, cal4=3.77E+02)
```

Computes non-linear modelling of calibration curves

```
calibCurves <- Calibrate(mrDataOrFile=calibFile,
                         calibValues=cellConcentrations,
                         pdfName=NULL)    # or =name_of_pdf_file
```

```
##
## Reading MiniRead output file:
##  /home/mfalque/R/x86_64-pc-linux-gnu-library/4.3/MiniRead/extdata/CALI_053.MRA
## Looking for saturated values
```

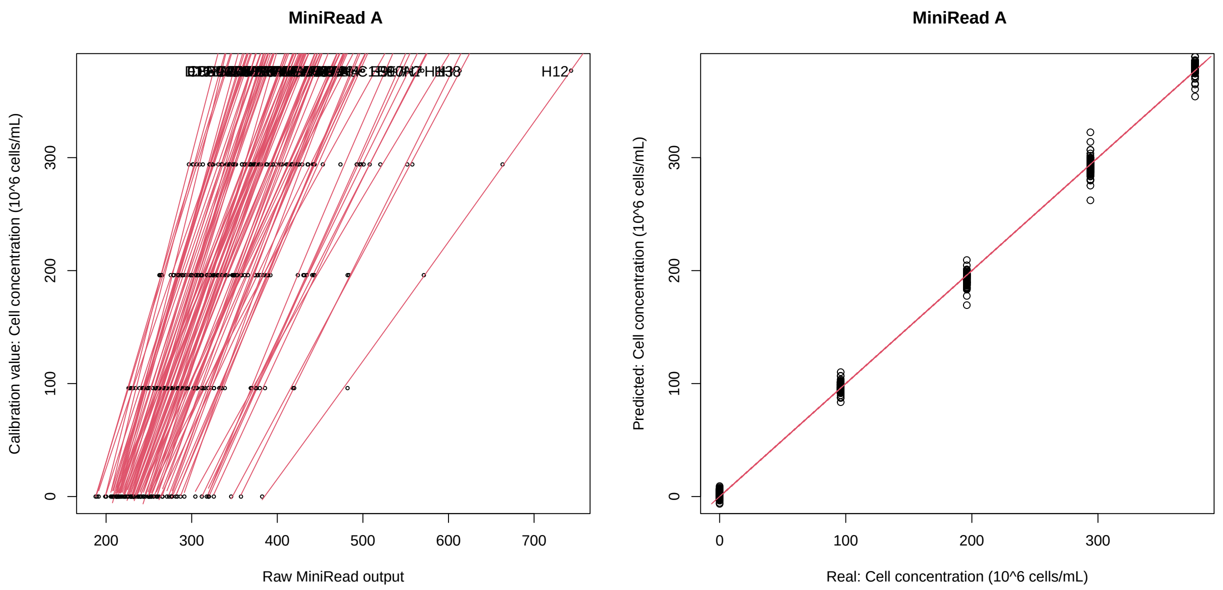


```
##
## Calibration curves saved successfully in:
## /home/mfalque/MiniRead_Calibration/MiniRead_CALIBRATION.rds
## Calibration date 2023-12-20_08h52m48
## Calibration data type: Cell concentration
## Calibration units: 10^6 cells/mL
## File used for calibration: CALI_053.MRA
## Devices used for calibration: A
## Number of wells calibrated: 96
##
## Calibration completed
```

The function Calibrate() writes the computed calibration parameters (calibCurves, which contains the non-linear models of the calibration curves) in the folder 'MiniRead_Calibration' of your home directory.

Some information on the calibration data (including calibration date, labels of the MiniReads devices used for calibration, etc…) is attached as attributes of the 'calibCurves' object:

```r
attributes(calibCurves)
```

```
## $calibDate
## [1] "2023-12-20_08h52m48"
##
## $calibUnit
## [1] "10^6 cells/mL"
##
## $calibType
## [1] "Cell concentration"
##
```

```
## $calibFile
## [1] "CALI_053.MRA"
##
## $calibDevices
## [1] "A"
##
## $calibWells
##  [1] "W1"  "W2"  "W3"  "W4"  "W5"  "W6"  "W7"  "W8"  "W9"  "W10" "W11" "W12"
## [13] "W13" "W14" "W15" "W16" "W17" "W18" "W19" "W20" "W21" "W22" "W23" "W24"
## [25] "W25" "W26" "W27" "W28" "W29" "W30" "W31" "W32" "W33" "W34" "W35" "W36"
## [37] "W37" "W38" "W39" "W40" "W41" "W42" "W43" "W44" "W45" "W46" "W47" "W48"
## [49] "W49" "W50" "W51" "W52" "W53" "W54" "W55" "W56" "W57" "W58" "W59" "W60"
## [61] "W61" "W62" "W63" "W64" "W65" "W66" "W67" "W68" "W69" "W70" "W71" "W72"
## [73] "W73" "W74" "W75" "W76" "W77" "W78" "W79" "W80" "W81" "W82" "W83" "W84"
## [85] "W85" "W86" "W87" "W88" "W89" "W90" "W91" "W92" "W93" "W94" "W95" "W96"
```

#### 3. Apply calibration curves to kinetics data (example with only one device)

Load example kinetics data (device "B")

```
library(MiniRead)
kineFile <- ExampleFile("KINE_B")
nonCalibratedData <- ReadMiniReadData(kineFile)
```

```
##
## Reading MiniRead output file:
##  /home/mfalque/R/x86_64-pc-linux-gnu-library/4.3/MiniRead/extdata/KINE_026.MRB
```

Raw turbidity values (arbitrary units) recorded around the middle of expo phase:

```
nonCalibratedData[150:160, c(1,8:13)]
```

```
##        hours     W1     W2     W3     W4     W5     W6
## 150 12.56899 247.40 263.80 249.84 238.52 249.44 272.55
## 151 12.65336 248.55 265.27 251.26 239.92 251.11 275.39
## 152 12.73773 249.76 266.54 252.99 241.31 252.93 274.86
## 153 12.82212 251.13 268.26 254.57 242.83 254.69 276.75
## 154 12.90649 252.68 269.75 255.96 244.36 256.57 278.92
## 155 12.99087 254.16 271.38 257.73 245.94 258.26 280.30
## 156 13.07524 255.45 273.10 259.47 247.46 260.17 281.91
## 157 13.15963 257.15 274.75 261.19 249.17 262.25 283.68
## 158 13.24401 258.59 276.37 263.09 250.83 264.11 285.78
## 159 13.32838 260.28 278.04 265.14 252.37 266.22 287.65
## 160 13.41276 261.96 279.85 267.12 254.02 268.30 289.24
```

Load calibration data for the same device ("B")

```
calibFile <- ExampleFile("CALI_B")
```

Real cell concentrations (in Million Cells/mL) used for each point of the calibration series

```
cellConcentrations <- c(cal0=0.00E+00, cal1=9.60E+01, cal2=1.96E+02,
                        cal3=2.94E+02, cal4=3.77E+02)
```

Now computing calibration curves

```
calibCurves <- Calibrate(mrDataOrFile=calibFile,
                         calibValues=cellConcentrations,
                         pdfName=NULL)     # or =name_of_pdf_file
```

```
##
## Reading MiniRead output file:
##  /home/mfalque/R/x86_64-pc-linux-gnu-library/4.3/MiniRead/extdata/CALI_025.MRB
## Looking for saturated values
```

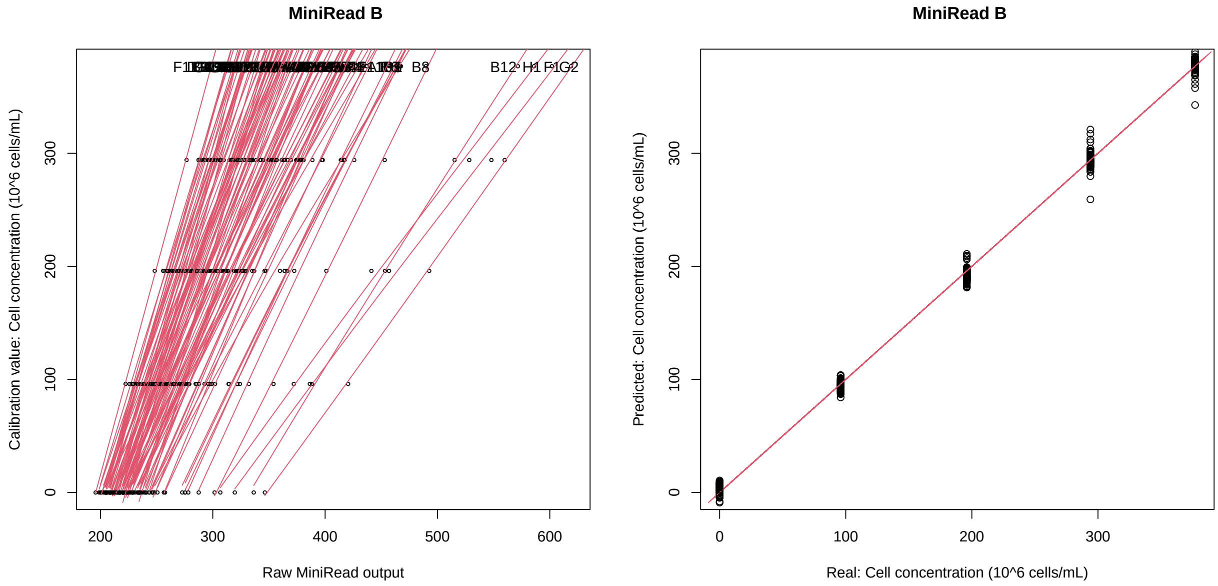


```
##
## Calibration curves saved successfully in:
## /home/mfalque/MiniRead_Calibration/MiniRead_CALIBRATION.rds
## Calibration date 2023-12-20_08h52m50
## Calibration data type: Cell concentration
## Calibration units: 10^6 cells/mL
## File used for calibration: CALI_025.MRB
## Devices used for calibration: B
## Number of wells calibrated: 96
##
## Calibration completed
```

And applying this calibration to the kinetics results:

```
calibratedData <- ApplyCalibration(mrDataOrFile=kineFile,
                                   calibrationCurves=calibCurves)
```

```
##
## Reading MiniRead output file:
##  /home/mfalque/R/x86_64-pc-linux-gnu-library/4.3/MiniRead/extdata/KINE_026.MRB
## Calibration date 2023-12-20_08h52m50
## Calibration data type: Cell concentration
## Calibration units: 10^6 cells/mL
## File used for calibration: CALI_025.MRB
## Devices used for calibration: B
## Number of wells calibrated: 96
##
## Applying calibration to KINETICS file
```

After conversion to Millions cells/mL by applying calibration curves, the turbidity values (same points around the middle of expo phase) are:

```
calibratedData[150:160, c(1,8:13)]
```

```
##        hours        W1        W2        W3        W4        W5        W6
## 150 12.56899  67.47778  76.97530  76.03484  87.88350  74.84608  80.92935
## 151 12.65336  70.18175  80.76543  79.81884  91.38874  78.79292  87.22260
## 152 12.73773  73.02604  84.03584  84.42289  94.86536  83.08659  86.04893
## 153 12.82212  76.24545  88.45872  88.62161  98.66294  87.23062  90.23263
## 154 12.90649  79.88652  92.28399  92.31046 102.48098  91.64793  95.03023
## 155 12.99087  83.36180  96.46177  97.00120 106.41890  95.61039  98.07779
## 156 13.07524  86.38978 100.86212 101.60540 110.20251 100.07893 101.62976
## 157 13.15963  90.37847 105.07530 106.15001 114.45334 104.93328 105.53020
## 158 13.24401  93.75555 109.20393 111.16271 118.57395 109.26351 110.15147
## 159 13.32838  97.71703 113.45152 116.56258 122.39134 114.16338 114.26056
## 160 13.41276 101.65294 118.04518 121.76987 126.47558 118.98050 117.74976
```

#### 4. Apply calibration curves to kinetics data (example with multiple devices)

Here, the same calibration plates (containing a five-points dilution series of cell suspension) have been used to calibrate 5 different MiniRead devices (named 'A' to 'E')

Load example kinetics data file path from the package

```
library(MiniRead)
kineFile <- ExampleFile("KINE_B")
```

Load example calibration data of devices "A" to "E"

```
calibFiles <- c(ExampleFile("CALI_A"), ExampleFile("CALI_B"),
               ExampleFile("CALI_C"), ExampleFile("CALI_D"),
               ExampleFile("CALI_E")
              )
```

Concatenate all data in a single data frame

```
allCalis <- NULL
for (file in calibFiles) {
 thisCali <- ReadMiniReadData(file)
 allCalis <- rbind.data.frame(allCalis, thisCali)
}
```

```
##
## Reading MiniRead output file:
##  /home/mfalque/R/x86_64-pc-linux-gnu-library/4.3/MiniRead/extdata/CALI_053.MRA
##
## Reading MiniRead output file:
##  /home/mfalque/R/x86_64-pc-linux-gnu-library/4.3/MiniRead/extdata/CALI_025.MRB
##
## Reading MiniRead output file:
##  /home/mfalque/R/x86_64-pc-linux-gnu-library/4.3/MiniRead/extdata/CALI_024.MRC
##
## Reading MiniRead output file:
##  /home/mfalque/R/x86_64-pc-linux-gnu-library/4.3/MiniRead/extdata/CALI_026.MRD
##
## Reading MiniRead output file:
##  /home/mfalque/R/x86_64-pc-linux-gnu-library/4.3/MiniRead/extdata/CALI_042.MRE
```

Real cell concentrations used for each point of the calibration series

```
cellConcentrations <- c(cal0=0.00E+00, cal1=9.60E+01, cal2=1.96E+02,
                        cal3=2.94E+02, cal4=3.77E+02)
```

Compute and model calibration curves

```
calibCurves <- Calibrate(mrDataOrFile=allCalis,
                         calibValues=cellConcentrations)
```

```
## Checking format of data frame 'mrDataOrFile'
## Looking for saturated values

## Graphics written in file:
## /home/mfalque/DATA/GIT_DEV/MiniRead/user-manual/Calibration_MiniRead.pdf
## Calibration curves saved successfully in:
## /home/mfalque/MiniRead_Calibration/MiniRead_CALIBRATION.rds
## Calibration date 2023-12-20_08h52m52
## Calibration data type: Cell concentration
## Calibration units: 10^6 cells/mL
## File used for calibration: NA
## Devices used for calibration: A B C D E
## Number of wells calibrated: 96
##
## Calibration completed
```

The list of devices appears in the attributes:

```
attributes(calibCurves)$calibDevices
```

```
## [1] "A" "B" "C" "D" "E"
```

#### 5. Complete automatic analysis

This requires that calibration has already been carried out for the device(s) used, and that the calibCurves files have been written in the folder package.

**Plot well 'A1' of the plate**
* *with growth parameter analysis*
* *with correction for evaporation*
* *with applying calibration curves*

```
res <- AnalyzeGrowth(mrDataOrFile=kineFile,
                     maxTime=48,
                     wells="A1",
                     useCalibration=TRUE,
                     pdfName=NULL,
                     flatPlateau=TRUE,
                     writose=FALSE,
                     grOpt=list(text=TRUE, deriv=FALSE, lines=TRUE))
```

```
##
## Reading MiniRead output file:
##  /home/mfalque/R/x86_64-pc-linux-gnu-library/4.3/MiniRead/extdata/KINE_026.MRB
## Checking format of data frame 'mrDataOrFile'
##
## Reading calibration curves from folder:
## /home/mfalque/MiniRead_Calibration/MiniRead_CALIBRATION.rds
## Calibration date 2023-12-20_08h52m52
```

```
## Calibration data type: Cell concentration
## Calibration units: 10^6 cells/mL
## File used for calibration: NA
## Devices used for calibration: A B C D E
## Number of wells calibrated: 96
##
## Applying calibration to KINETICS file
## Preparing plots and analyzing kinetics parameters
```

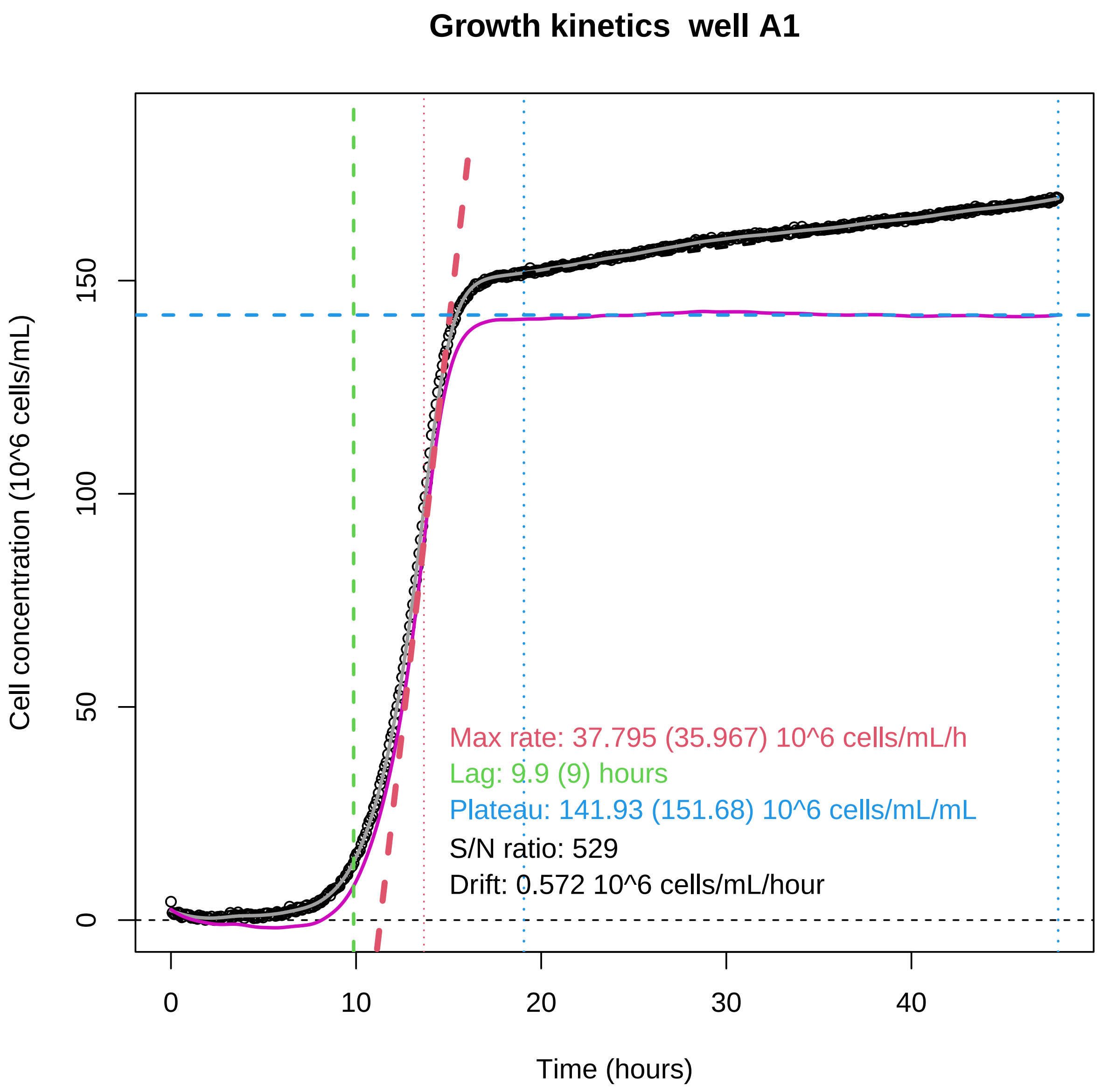


```r
head(res)
```

```
##   device line col maxRateRaw lagRaw plateauRaw snr drift maxRateCorr lagCorr
## 1      B    A   1     35.967      9     151.68 529 0.572      37.795     9.9
##   plateauCorr
## 1      141.93
```
