## Supplementary Figure S1 for "MiniRead: a simple and inexpensive do-it-yourself device for multiple analyses of micro-organism growth kinetics"

**Figure S1.** One of the two heating plates (under the plate and on top of the lid), showing the serpentine of resistive wire.

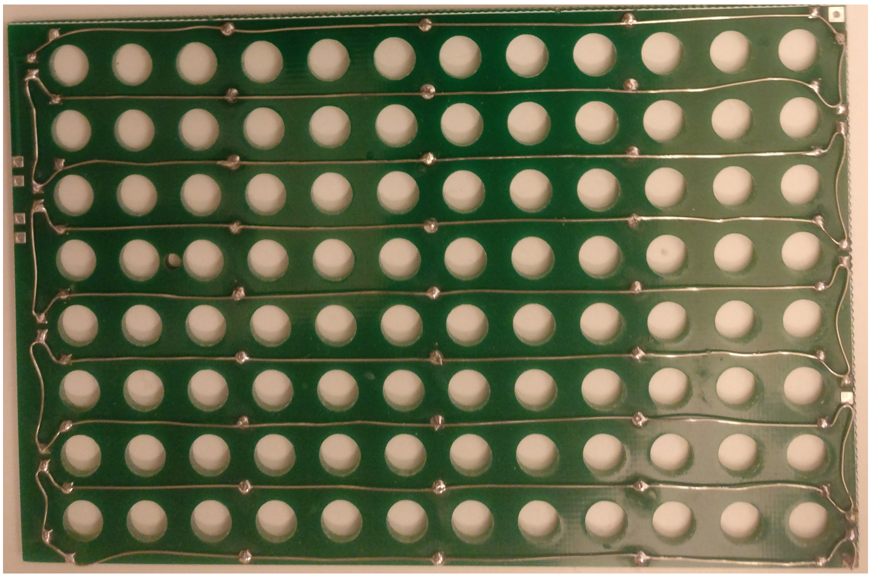
