## Supplementary Figure S2 for "MiniRead: a simple and inexpensive do-it-yourself device for multiple analyses of micro-organism growth kinetics"

**Figure S2**. Top view of the user interface showing real-time display of the parameters (plate and lid temperatures, coordinates of the well being acquired, raw value read, brightness of the LEDs, and name of the current file being written into.

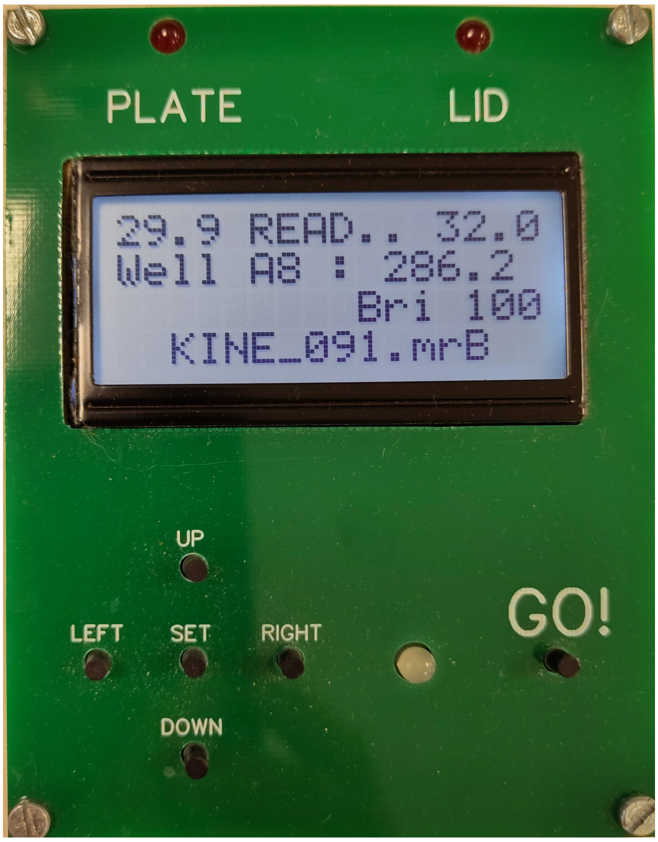
