## Supplementary Figure S3 for "MiniRead: a simple and inexpensive do-it-yourself device for multiple analyses of micro-organism growth kinetics"

**Figure S3.** Kinetics of cell concentration showing the concentration of the liquid medium by evaporation for 60 hours at 30°C in a plate well containing 200 uL of sterile YPD without any inoculation. Black circles: experimental points. Red curve: linear regression. Values of Y-axis are in arbitrary units.

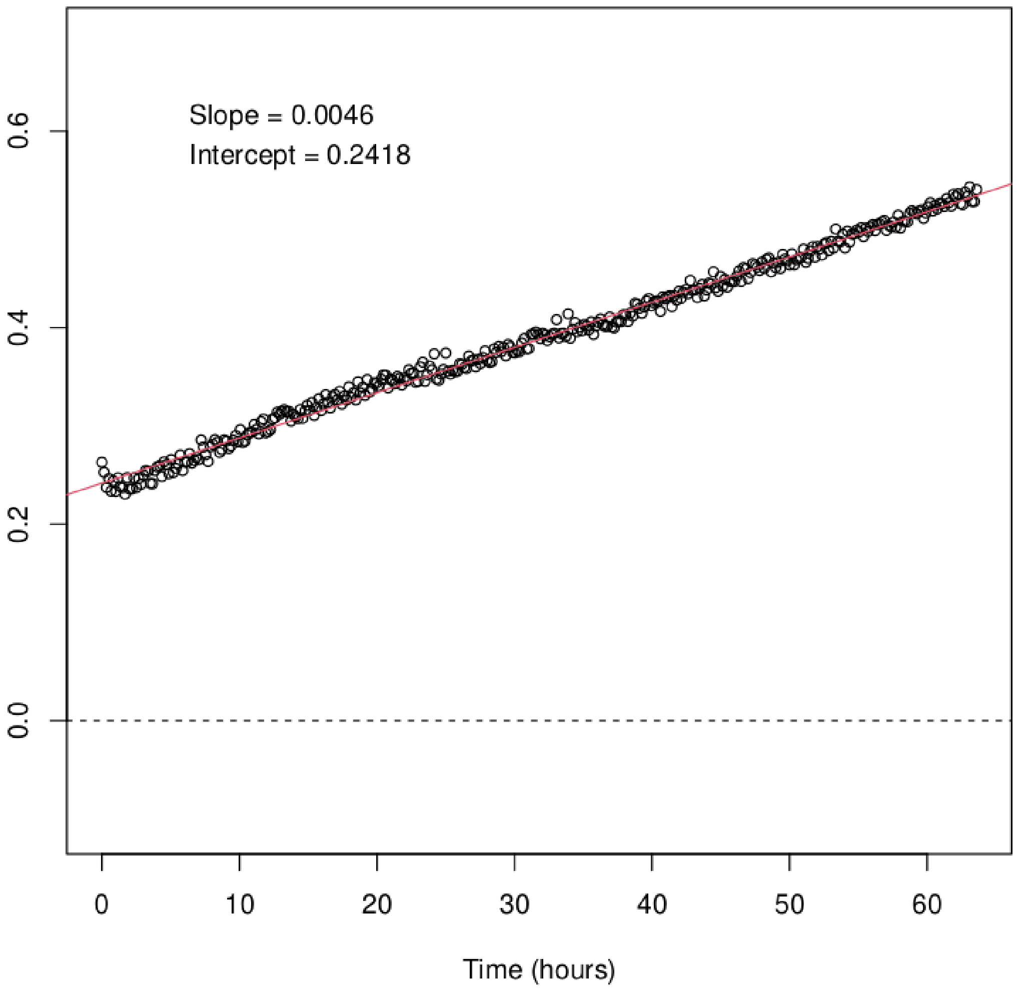
